# Engineering Membrane-Bound Alkane Monooxygenase from *Marinobacter* sp. for Increased Activity in the Selective ω-Hydroxylation of Linear and Branched Aliphatic Esters

**DOI:** 10.1101/2025.08.27.672531

**Authors:** Jelena Spasic, Andrea Nigl, Huijin Cheon, Christine L. Kaiserer, Stela Galušić, Elske van der Pol, Lenny Malihan-Yap, Jin-Byung Park, Robert Kourist

**Affiliations:** Graz University of Technology, Institute of Molecular Biotechnology, Petersgasse 14, 8010 Graz, Austria; Department of Food Science and Biotechnology, Ewha Womans University, Seoul 03760, Republic of Korea; Graz University of Technology, Institute of Organic Chemistry, Stremayrgasse 9, 8010, Graz, Austria

**Author notes:** **Corresponding Authors:** Robert Kourist, Graz University of Technology, Institute of Molecular Biotechnology, Petersgasse 14, 8010 Graz, Austria, Jin-Byung Park, Department of Food Science and Biotechnology, Ewha Womans University, Seoul 03760, Republic of Korea. These authors contributed equally (Jelena Spasic, Andrea Nigl, Huijin Cheon).

**Keywords:** alkane monooxygenase, enzyme engineering, oxyfunctionalization, polymer precursors, regio- and stereoselective hydroxylation

## Abstract

The regio- and stereoselective hydroxylation of unactivated C(sp^3^)-H bonds is an important reaction in organic synthesis. While bacterial alkane monooxygenase AlkB catalyzes the terminal hydroxylation of aliphatic esters with excellent regioselectivity, the molecular principles of substrate recognition and selectivity of this integral membrane enzyme are still poorly understood. In this study, we investigated the substrate scope and engineered the medium-chain alkane monooxygenase from *Marinobacter* sp. (M_AlkB) for the terminal hydroxylation of linear and branched esters of fatty acids and alcohols. For the first time, we demonstrated the stereoselectivity of AlkB towards prochiral substrates containing terminal *gem*-dimethyl groups, leading to the corresponding chiral *β*-methyl primary alcohols in good optical purity (51-79% *ee*). The hydroxylation products can be further derivatized to chiral diols and lactones. Substitution of the highly conserved active site residue F169 to leucine increased the activity towards short and medium-chain esters up to two-fold. While the wildtype enzyme does not accept long-chain substrates, activity towards *n*-dodecyl acetate could be unlocked by reducing the size of the tryptophan residue 60 situated in the putative substrate tunnel. Substitution of the peripheral I238 with valine increased activity regardless of the chain length of the substrate. Our results lay the groundwork for the establishment of a whole-cell process for the regio- and stereoselective hydroxylation of linear and branched esters, leading to valuable bifunctionalized products. The insights gained from mutating key residues and the substrate acceptance of AlkB will guide future protein engineering campaigns.

**SIGNIFICANCE:** We report rational engineering of a membrane-bound alkane monooxygenase in the hydroxylation of aliphatic esters, giving rise to diols, hydroxy acids, and lactones. By site-directed mutagenesis, we increased the activity towards short and medium-chain substrates, and we unlocked the activity towards long-chain substrates. We uncovered the enzyme’s capacity to discriminate between prochiral methyl groups, establishing new routes for the synthesis of chiral diols and lactones, important building blocks for the pharmaceutical industry.

## INTRODUCTION

The selective oxyfunctionalization of inherently inert C(sp^3^)–H bonds into more reactive C–O bonds is a highly important reaction in organic chemistry. It enables the synthesis of value-added compounds such as alcohols, epoxides, aldehydes, ketones, and carboxylic acids (Goldman & Goldberg, 2004; Münch, Püllmann, Zhang, & Weissenborn, 2021; Siedlecka, 2023; White, 2012; Wu, Paul, & Hollmann, 2023), which can also be further functionalized to more complex molecules (Behmagham et al., 2024; Jelen & Tavčar, 2025; Ji et al., 2025; Leech & Lam, 2020; Moon et al., 2023). Challenges associated with the direct oxygenation of aliphatic C–H bonds include regio-, chemo-, and stereoselectivity, product yields, and sustainability (Goldman & Goldberg, 2004; Münch et al., 2021; Siedlecka, 2023; White, 2012; Wu et al., 2023). Nature has evolved numerous enzymes, typically relying on metal cofactors, that are capable of selectively oxyfunctionalizing C(sp^3^)–H bonds of various substrates (Austin & Groves, 2011; Dong et al., 2018; Münch et al., 2021; Wu et al., 2023). Many heme-dependent cytochrome P450s (CYP450) target subterminal ω-1 and ω-2 positions, with only a few being specific for the terminal position (Ebrecht, Aschenbrenner, Gumulya, Smit, & Opperman, 2025; Fiorentini et al., 2018; Funhoff, Bauer, García-Rubio, Witholt, & van Beilen, 2006; Hammerer, Winkler, & Kroutil, 2018; Honda Malca et al., 2012; Meinhold, Peters, Hartwick, Hernandez, & Arnold, 2006; Schultes, Welter, Schmidtke, Tischler, & Mügge, 2024). In contrast, the diiron non-heme alkane monooxygenase AlkB exhibits exceptional regioselectivity for the terminal hydroxylation of linear aliphatic compounds. AlkB and its electron transfer accessory proteins were first isolated from the soil bacterium *Pseudomonas putida* GPo1 (PpAlkB; recently renamed to *Ectopseudomonas oleovorans,* formerly known as *P. oleovorans*) (McKenna & Coon, 1970; Peterson, Basu, & Coon, 1966; Peterson & Coon, 1968). Since then, various homologs from other soil and marine bacteria have been identified (Nie, Liang, Fang, Tang, & Wu, 2011; Ratajczak, Geißdörfer, & Hillen, 1998; Smits, Balada, Witholt, & van Beilen, 2002; Van Beilen et al., 2004, 2002; Williams & Austin, 2022; Williams et al., 2021; Williams, Luongo, Orman, Vizcarra, & Austin, 2022). AlkB is part of the *alk*-operon that enables bacteria to use *n*-alkanes as their sole energy and carbon source. AlkB catalyzes the initial ω-hydroxylation of *n*-alkanes to 1-alkanols, with two electrons provided by rubredoxin AlkG, which is in turn reduced by reductase AlkT (Figure 1A) (Owen et al., 1984; van Beilen et al., 2001; van Beilen, Wubbolts, & Witholt, 1994). In addition to linear medium-chain hydrocarbons (C_6_ to C_12_), AlkB also acts on branched, cyclic, and aromatic compounds. Terminal olefin groups are typically converted to epoxides (May & Abbott, 1973; May, Gordon, & Steltenkamp, 1977; May & Katopodis, 1986; van Beilen, Kingma, & Witholt, 1994). Recently, the dehalogenation of ω-fluorinated hydrocarbons to aldehydes by AlkB has been reported (Hendricks et al., 2025; Xie et al., 2023). The well-studied PpAlkB has been used to hydroxylate esters of fatty acids (FAcE) and fatty alcohols (FAlE) for the production of polyester precursors (Julsing et al., 2012; Schrewe et al., 2014; Schrewe, Magnusson, Willrodt, Bühler, & Schmid, 2011; van Nuland, de Vogel, Eggink, & Weusthuis, 2017; van Nuland, de Vogel, Scott, Eggink, & Weusthuis, 2017; van Nuland, Eggink, & Weusthuis, 2016). The terminal oxyfunctionalization in conjunction with amination yields precursors for polyamides (Herzog et al., 2020; Ladkau et al., 2016; Schrewe, Ladkau, Bühler, & Schmid, 2013). The α,ω-oxyfunctionalized products also find applications in the pharmaceutical (Rivière et al., 2020), cosmetic (Kosmadaki & Katsambas, 2017) and agricultural industry (Z. Wang, Xu, Tian, & Pan, 2007).

**Figure 1.**
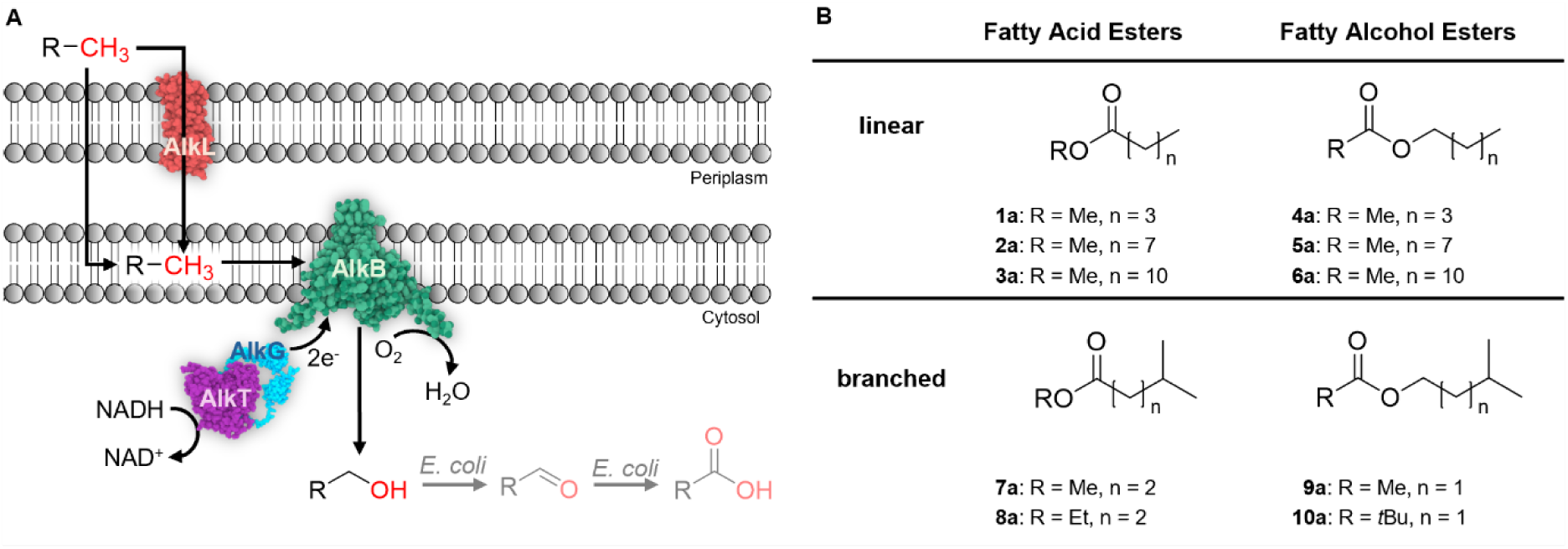
A) Terminal hydroxylation of aliphatic compounds by alkane monooxygenases AlkB and potential oxidation of the resulting alcohol to the aldehyde and carboxylic acid by the host background. B) Linear and branched esters investigated in this study.

The challenges associated with the kinetic characterization of membrane-bound enzymes and the elucidation of their structures have hampered their engineering. The emergence of cryogenic electron microscopy (cryo-EM) (Chai, Guo, McSweeney, Shanklin, & Liu, 2023; Guo et al., 2023; Khanppnavar et al., 2024) and *de novo* structure prediction tools (Abramson et al., 2024; Jumper et al., 2021) have significantly increased our structural understanding of membrane-embedded biocatalysts, allowing their engineering (Choo, Sirota, See, Eisenhaber, & Li, 2023). In the case of AlkB, only recently, the first structures of two homologs from *Fontimonas thermophila* (FtAlkB) were determined by cryo-EM (Chai et al., 2023; Guo et al., 2023). However, the absence of high-throughput screening (HTS) assays for unnatural substrates has limited its engineering to random mutagenesis and selection based on the utilization of natural *n*-alkane substrates (Koch, Chen, van Beilen, & Arnold, 2009; van Beilen et al., 2005).

In a recent study, we engineered PpAlkB to catalyze the crucial terminal hydroxylation of isoprenyl acetate in a multi-enzyme cascade towards the highly interesting polymer precursor tulipalin A (Nigl et al., 2025). AlkB exclusively acted on the terminal methyl group while leaving the adjacent *exo-*olefin intact. The insights from engineering PpAlkB could be transferred to AlkB from *Marinobacter* sp. (M_AlkB), which displayed the highest hydroxylation activity among several homologs (Nigl et al., 2025).

Although AlkB is known to hydroxylate (ω-1)-methyl-substituted alkanes, it prefers the linear terminus (van Beilen, Kingma, et al., 1994). We assumed that the hydroxylation activity could be directed towards the prochiral methyl groups of the alkyl chain by protecting the linear end with a small ester group. The stereoselective hydroxylation of prochiral esters by AlkB leads to optically pure primary alcohols, thereby opening new pathways towards chiral hydroxy acids, diols, and lactones. These chiral compounds present interesting building blocks, not only for polymer production (J. Engel, Cordellier, Huang, & Kara, 2019; Gubbels et al., 2018; Schaffer & Haas, 2014; Spanjers et al., 2016), but also for pharmaceuticals (Suzuki, Usui, Miyake, Namikoshi, & Nakada, 2004; Takahashi, Komine, Yokoi, Ishihara, & Hatakeyama, 2012).

In this work, we aimed to shed light on the substrate scope, activity, and stereoselectivity of M_AlkB. Therefore, we investigated the effect of selected point mutations in the active site, substrate tunnel, and membrane-cytosol interface on the reactivity of M_AlkB towards a set of linear and branched esters of fatty acids and alcohols varying in the length of the alkyl chain (Figure 1B).

## Results and Discussion

### Characterization of the alkane monooxygenase M_AlkB from *Marinobacter* sp

Petroleum-contaminated areas show an enrichment of *alkB* genes and prove to be a vast source of *alkB* sequences (Head, Jones, & Röling, 2006; Vomberg & Klinner, 2000; Yang, Wang, Liao, Xie, & Huang, 2015). Bacteria of the genus *Marinobacter* are highly abundant in such areas, and are known for their ability to grow on alkanes, with most strains showing a preference for medium chain *n*-alkanes ranging from C_6_ to C_8_ (Gauthier et al., 1992; Gunasekera, Bowen, Radwan, Striebich, & Ruiz, 2022; Yang et al., 2015). Here, we characterize an AlkB from *Marinobacter* sp., which is derived from a metagenome sample (Tully, Graham, & Heidelberg, 2018), and was shown to have 1.6 and 2.3-fold higher hydroxylation activity compared to the well-described PpAlkB towards *n*-octane and isoprenyl acetate, respectively (Nigl et al., 2025). M_AlkB shares 78% sequence similarity with PpAlkB (Eggink, Lageveen, Altenburg, & Witholt, 1987; McKenna & Coon, 1970; Owen et al., 1984) and 66% and 51% similarity with the AlkBs from *Fontimonas thermophila* FtAlkB (Guo et al., 2023) and FtAlkBG (Chai et al., 2023), respectively, whose structures have been recently solved by cryo-EM (Figure S2). *M_alkB* was expressed in *E. coli* BL21 (DE3) under the control of the natural alkane-inducible *P_alkB_* promoter encoded on the pCom10 plasmid (Smits et al., 2001) together with the redox partners *alkFGT* and the transporter *alkL* from *P. putida* GPo1 (Figure S5). An initial substrate screening showed that M_AlkB converted esters of different chain lengths (**1a**-**6a**) to their respective hydroxy products (**1b**-**6b**; Figure S9). Although the initial screening was performed over 24 h, product formation reached saturation within 4 h. Therefore, whole-cell activity measurements were performed using a 4 h reaction time. The mass balances of the biotransformations were not fully closed, which could be attributed to substrate or product volatility, hydrolysis, or losses during extraction steps. We observed that the fatty acid methyl esters (FAcME) **1a**-**3a** were partially hydrolyzed to the respective acids **1**-**3** (Figure S9A-C), which prevented efficient quantification of the enzymatic activity for **1a** and **3a**. FAcME hydrolysis was previously reported and attributed to endogenous esterases from *E. coli* (Julsing et al., 2012; Kadisch, Schmid, & Bühler, 2017; Schrewe et al., 2013; van Nuland et al., 2016). In whole-cell reactions with the acetate esters **4a**-**6a**, less hydrolysis was observed (Figure S9D-E). The conversion by the wildtype (WT) PpAlkB and M_AlkB of the C_12_ chain esters **3a** and **6a** was very low. This follows the previously reported substrate preference for PpAlkB (Schrewe et al., 2011; van Beilen, Kingma, et al., 1994; van Nuland et al., 2016). The substrate chain length accepted by M_AlkB fits in the range of medium-chain monooxygenases, which is in line with the reported activity towards *n*-octane and isoprenyl acetate (Nigl et al., 2025).

We determined the whole-cell activity for the nonanoic acid methyl ester **2a** (van Nuland et al., 2016) and three fatty alcohol acetates (FAlAc) with varying chain lengths (**4a**, **5a**, and **6a**). M_AlkB showed comparable activity to PpAlkB towards **2a** (5.8 U/g_CDW_), whereas the activity towards medium-chain acetates was up to 8-fold higher, reaching 2.7 U/g_CDW_ and 4.7 U/g_CDW_ for **4a** and **5a**, respectively (Figure 2B-E). For **6a**, we observed very low conversions with both enzymes (Figure S9F), limiting activity calculations. Interestingly, when comparing the whole-cell oxygenation rates of methyl nonanoate **2a** with the nonyl acetate **5a** by PpAlkB, the conversion of **2a** was significantly faster than that of **5a** (Figure 2CE). In reactions with M_AlkB such a notable difference was not observed. Even though **2a** and **5a** have the same alkyl chain length and are sterically very similar, their hydrophobicity differs (Table S6). The logP value of **2a** and **5a** is 4.32 and 3.95, respectively. We assumed that hydrophobicity and partition coefficient of the substrates could influence the whole-cell hydroxylation rates, especially when considering that the substrate is expected to enter the substrate tunnel leading to the enzyme active site via the lipid bilayer (Figure 1A) (Mikulska-Ruminska et al., 2025). Both enzymes’ tunnels differ in the second-shell residues neighboring the amino acids forming the tunnel, which are almost identical (Figure S3C-D). This makes it very difficult to rationalize the different acceptance of methyl alcohol esters and acetate esters by both membrane monooxygenases.

**Figure 2.**
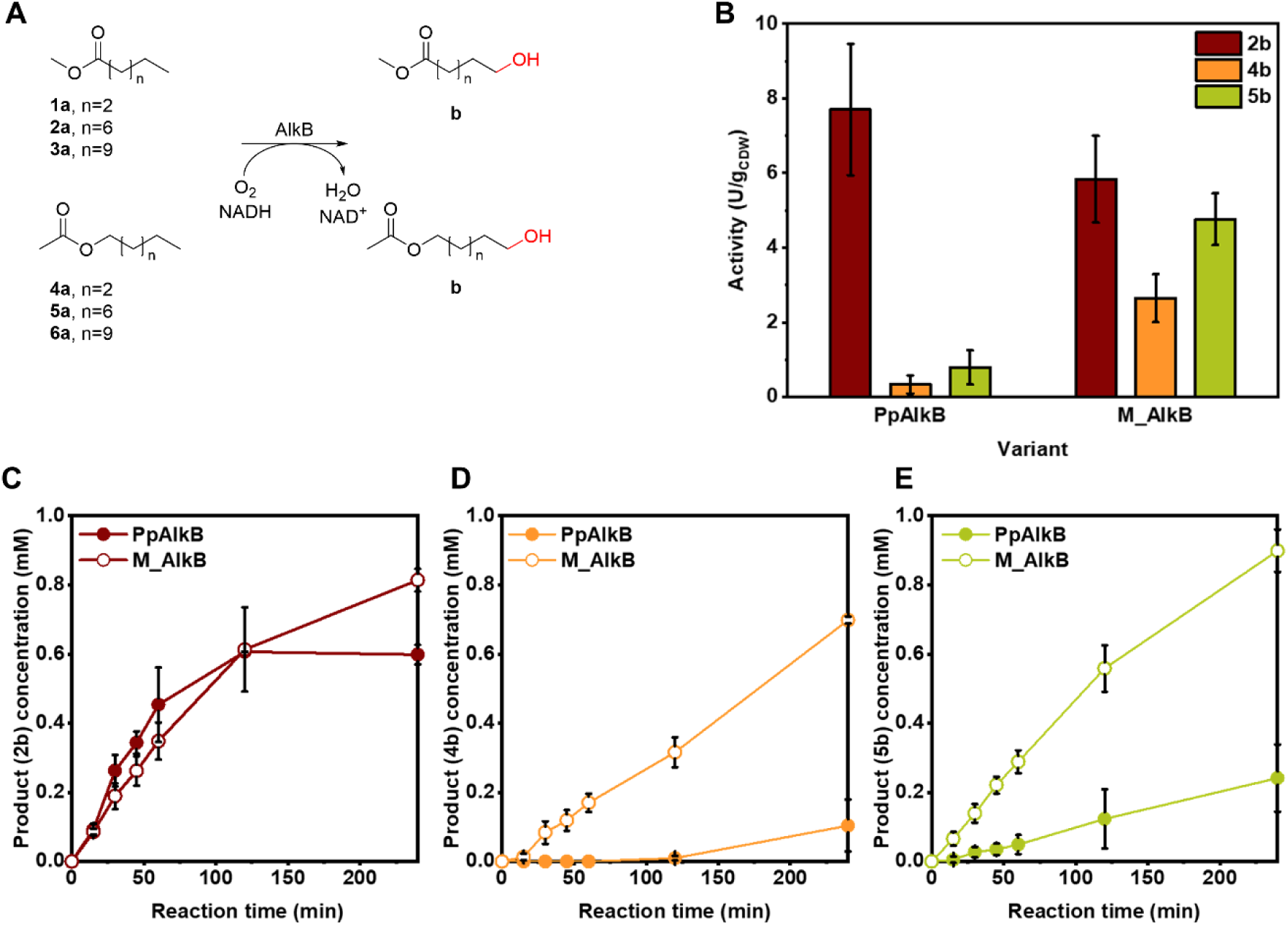
Biotransformation of linear FAcME and FAlAc by PpAlkB and M_AlkB. A) Tested substrates and expected products; only residual (<2%) conversion of substrates **1a**, **3a**, and **6a** was observed. B) Activity of PpAlkB and M_AlkB towards FAcME **2a** and FAlAc **4a** and **5a**. Time-course of product formation for C) **2b**, D) **4b**, and E) **5b** by PpAlkB (filled circles) and M_AlkB (empty circles). The reactions were performed for 4h at 1 g_CDW_/L (mean ± SD; *n* = 3).

### Enzyme engineering for increased activity towards linear esters

The growth assay developed by van Beilen (Koch et al., 2009; van Beilen et al., 2005). is not applicable for the esters **1a**-**6a** since they can be hydrolyzed by the cells and directly fed into the β−oxidation pathway, rendering their terminal hydroxylation unnecessary for their utilization as a carbon source. Therefore, we investigated the effect of point mutations of residues in the active site, the substrate tunnel, and the periphery. Figure 3 highlights the residues targeted by mutagenesis in the 3D structure of M_AlkB generated by AlphaFold 3 (Abramson et al., 2024). The putative substrate tunnel was predicted using CAVER Web 2.0 (Figure 3A) (Marques et al., 2025). The modeled enzyme consists of six transmembrane helices arranged in a slightly cone-like shape, opening towards the cytosol. Helices 2, 4, and 6 are close to the predicted substrate tunnel, similar to those described for FtAlkB (Guo et al., 2023) and FtAlkBG (Chai et al., 2023). The predicted active site cavity is formed between the transmembrane helices and resembles a typical AlkB active site with the nine distinct, highly conserved histidines coordinating the two iron atoms essential for catalytic activity (Chai et al., 2023; Guo et al., 2023; van Beilen et al., 2005). The distance between two iron atoms in the active site is 5.9 Å and is in accordance with distances previously observed in the cryo-EM structures of two homologues (5.4 Å for FtAlkB and 6.1 Å for FtAlkBG) (Chai et al., 2023; Guo et al., 2023). Interestingly, the two iron atoms are unusually far apart (>5 Å) with no apparent bridging ligand between them, still raising questions regarding the reaction mechanism (Austin et al., 2008; Chai et al., 2023; Groves, Feng, & Austin, 2023; Guo et al., 2023; Mikulska-Ruminska et al., 2025; Reinhardt et al., 2025; Y. Wang & Liu, 2024).

**Figure 3.**
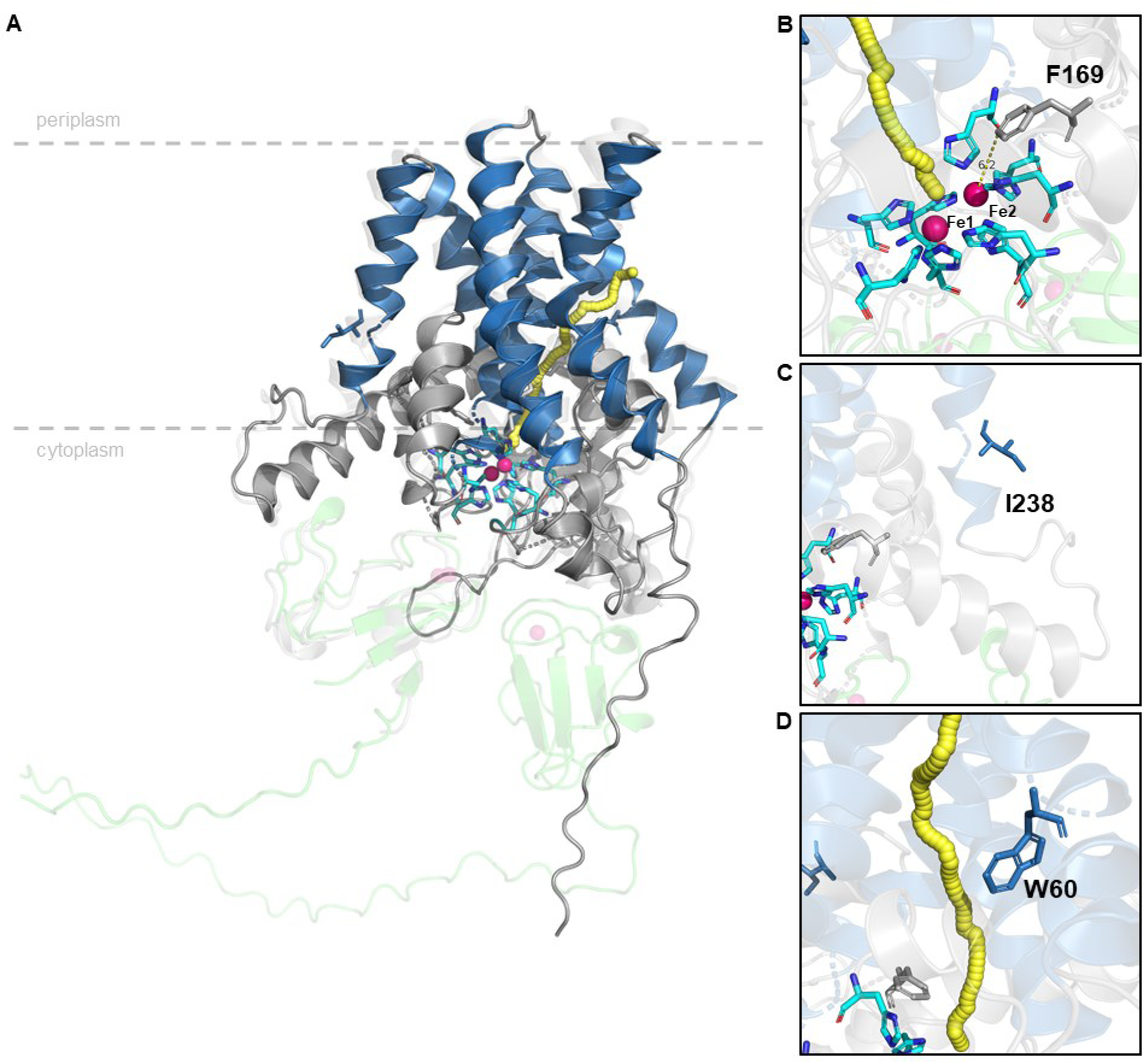
A) AlphaFold 3 predicted structure of M_AlkB (dark grey) with highlighted transmembrane domains (dark blue), active site histidine residues (cyano) coordinating two iron atoms (pink), and substrate tunnel (yellow). The interaction with PpAlkG (UniProt P00272, light green) was predicted by AlphaFold 3. The predicted structure of the protein complex was aligned with FtAlkBG (PDB 8f6t, light grey). AA residues targeted for site-directed mutagenesis: B) F169 close to the active site, C) I238 in the transmembrane domain facing the lipid bilayer, and D) W60 at the entrance of the predicted substrate tunnel.

To improve the enzyme’s activity, several point mutations were tested (Figure 3B-D). Residue I238 is located close to the periplasm in the fifth predicted transmembrane helix, distant from the active site (Figure 3C). The analog substitution of I233 to valine in PpAlkB was identified in a random mutagenesis-based directed evolution as part of a triple mutant, which showed improved activity towards butane and pentane (Koch et al., 2009). M_AlkB I238V showed higher activity towards *n-*octane and isoprenyl acetate (Nigl et al., 2025). As shown in Figure 4A, at a cell density of 1 g_CDW_/L, the mutation I238V increased the hydroxylation activity towards **2a**, **4a**, and **5a**, resulting in 9.7, 3.5, and 5.6 U/g_CDW_, respectively. Increasing the cell density to 3.1 g_CDW_/L still led to higher activity compared to the wildtype enzyme, reaching 6.7, 6.4, and 7.4 U/g_CDW_ for **2a**, **4a**, and **5a**, respectively (Figure 4B). At higher cell density, product formation rate for acetates increased up to 2-fold, while for methyl ester **2a**, the improvement was only 1.37-fold. Overall, this M_AlkB variant led to the highest product yields obtained after 2 h, resulting in 0.92 mM **2b** (Table S18). These results indicate that the I238V mutation indeed increases activity of AlkB towards linear esters in the whole-cell biocatalysts, independent of cell density and substrate chain length, which confirms results from previous engineering studies (Koch et al., 2009; Nigl et al., 2025).

**Figure 4.**
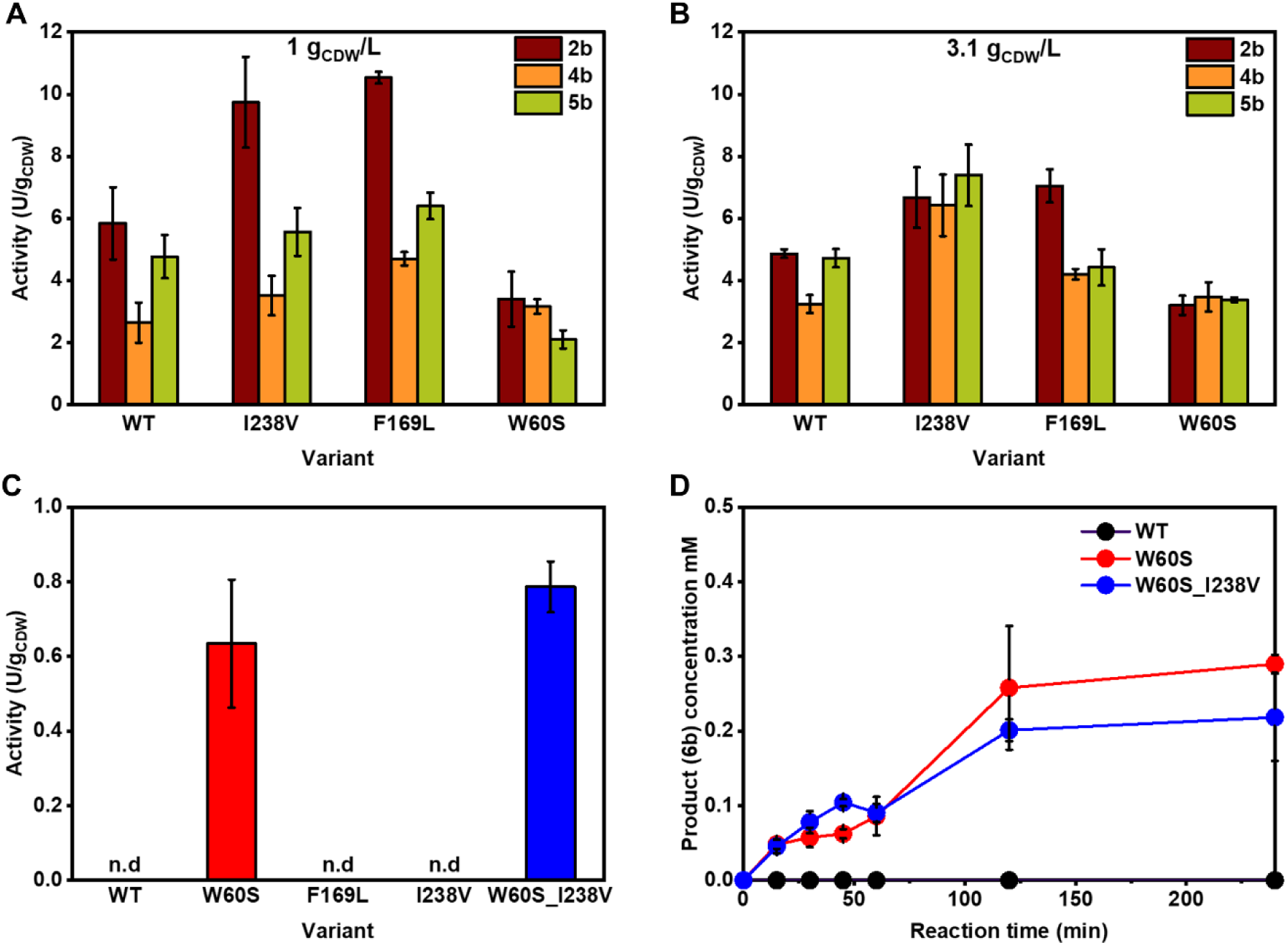
Hydroxylation activities of M_AlkB wildtype (WT) and variants using linear esters **2a**, **4a**, and **5a** at A) 1 g_CDW_/L and B) 3.1 g_CDW_/L. C) Activities and D) time-course of M_AlkB wildtype (WT) and W60S and W60S_I238V with **6a** as substrate at 3.1 g_CDW_/L. The reaction was performed for 4h (mean ± SD; *n* = 3). n.d. not determined.

The residue F169 is situated in the active site, being 6.2 Å distant from the second Fe atom (Fe2 in Figure 3B). Its substitution to leucine was recently reported to improve the activity towards isoprenyl acetate (Nigl et al., 2025). This residue is highly conserved among homologs (99% F, Table S1). It is, therefore, interesting that a change to leucine led to improved activity for **2a**, with up to a 1.8-fold increase reaching 10.5 U/g_CDW_. In contrast to I238V, the activity of F169L strongly depended on cell density, with a pronounced increase in activity at lower cell densities (1 g_CDW_/L). The dependence of enzyme activity on cell density could arise from transport limitations, substrate availability, or metabolic background. We expect that these are comparable between wt and the tested mutants. However, oxygen availability at different cell densities might have an effect on the reaction rates of wt and selected mutants.

In other AlkB homologs, the substitution of the tryptophan located at the entrance of the substrate tunnel (Figure 3D) in the second predicted transmembrane helix (W55 in PpAlkB, W62 in FtAlkB, W60 in M_AlkB) to a smaller amino acid, such as serine or valine, shifted the substrate acceptance of the monooxygenases towards longer substrates (> C_12_) (Guo et al., 2023; van Beilen et al., 2005). The position of this tryptophan in the predicted tunnel leading from the membrane as a possible substrate reservoir to the active site suggests that this residue might act as a gate-keeper for long-chain substrates (Guo et al., 2023; van Beilen et al., 2005). For the medium-chain esters **2a**, **4a**, and **5a**, the analogous mutation W60S in *M_alkB* did not lead to improved activity, but rather to a decreased hydroxylation rate, more pronounced at a lower cell density. However, this residue was tested, knowing that the acceptance of longer chain substrates might be unlocked. The variants I238V and F169L showed only traces of product formation in the conversion of **6a** (not shown), similar to the WT. As anticipated, M_AlkB W60S converted **6a**, reaching 0.63 U/g_CDW_ after 2 h of the reaction at 3.1 g_CDW_/L (Figure 4C, Table S18). In further testing with the M_AlkB double variant W60S_I238V, the activity gain with an initial activity of 0.79 U/g_CDW_ at 3.1 g_CDW_/L yielded 0.20 mM **6b** after 2h (Figure 4D, Table S18). Although the increase in the initial activity by the double mutation was modest (1.25-fold), the overall product concentration was comparable to the single mutant (0.26 mM **6b** after 2h).

Our results highlight that the knowledge gained by rational engineering can be transferred among AlkB homologs and that the effect on the activity towards alkanes (Guo et al., 2023; Koch et al., 2009; Nigl et al., 2025; van Beilen et al., 2005) can also influence the activity towards esters. For instance, the mutation I238V in *M_alkB* seems to improve the catalytic efficiency of the whole-cell biocatalyst independently of the substrate. The molecular effect of the peripheral substitution I238V remains to be clarified. Nevertheless, I238 is only 8.2 Å distant from a flexible loop closely situated to R215 and E232, which potentially play a role in the binding of AlkG (Groves et al., 2023; Mikulska-Ruminska et al., 2025). The limitation of M_AlkB towards longer chains could be unlocked via the single mutation W60S (0.63 U/g_CDW_) and further enhanced in the double variant W60S_I238V, reaching an initial activity of 0.79 U/g_CDW_ in the hydroxylation of the C12 substrate **6a**. This underlines the potential function of W60 as a gatekeeper in the substrate access tunnel (Guo et al., 2023; van Beilen et al., 2005). Overall, our rate improvement of whole-cell biocatalysts by enzyme engineering resulted in up to 2-fold improvements and is similar to fold-changes obtained previously, reporting 2.6-fold improved butane conversion (Koch et al., 2009) and 6-fold improved isoprenyl acetate conversion (Nigl et al., 2025). These results demonstrate that the protein engineering of M_AlkB increased the conversion of linear C_5_-C_12_ esters, representing a new starting point to produce ω-hydroxy acetates and diols.

### Activity and stereoselectivity of M_AlkB WT and mutants towards branched esters

AlkB monooxygenase has been previously reported to catalyze the allylic oxidation of sterically hindered substrates (Table S19), such as isobutene (Engel et al., 2015) and isoprenyl acetate (Nigl et al., 2025). We hypothesized that structurally similar substrates with a terminal *gem-*dimethyl group should also be converted, even though the steric hindrance is higher than that of the *exo*-olefin group. Due to the prochiral nature of the *gem-*dimethyl group, hydroxylation of one of the methyl groups would give rise to a chiral primary alcohol. May et al. reported the formation of enantiomerically enriched (*R*)*-*epoxides from 1-alkenes by PpAlkB (May & Abbott, 1972, 1973). In the conversion of terminally branched dimethyl alkanes, such as 2-methyl octane (**11a**), PpAlkB strongly preferred hydroxylation of the linear terminus (van Beilen, Kingma, et al., 1994). Indeed, the selectivity of M_AlkB for the hydroxylation of the linear over the branched end was 36-fold, leading to **11b** with only traces of chiral **11c** observed (Figure S12, Table S17). van Nuland et al. showed that PpAlkB does not act on terminal methyl ester or acetyl ester groups (van Nuland, de Vogel, Scott, et al., 2017; van Nuland et al., 2016). We hypothesized that in an ester composed of a germinal dimethyl end and a methyl ester or acetyl ester end, the activity of AlkB would be directed to the prochiral group, albeit with lower activity. This would allow us to determine the capacity of AlkB to discriminate between the two methyl groups (Figure 5A). To confirm our hypothesis, we investigated the hydroxylation of several prochiral esters **7a** to **10a**, varying in their chain length and the protective ester group (Figure 5A). The terminal hydroxylation of the branched FAcME **7a** would result in the chiral hydroxy fatty acid ester **7b**, which can easily be converted to the corresponding lactone **7e**. Similarly, terminal hydroxylation of the ethyl ester **8a** would result in the chiral ω-hydroxy fatty acid ester **8b**, which can be lactonized to **7e** (Figure 5A). On the other hand, terminal hydroxylation of the FAlE **9a** and **10a** would form the corresponding monoester of 2-methyl-1,4-butanediol (**9b**) (Figure 5A). We were pleased to see the formation of the corresponding hydroxy products **7b** and **9b** starting from prochiral **7a** and **9a**, respectively (Figure S13 and S14). The formation of **7b** was further confirmed by the presence of the corresponding lactone **7e**. The hydroxylation of **9a** to **9b** was confirmed by comparison with an authentic standard produced by the esterification of the diol **9g** using a *Pseudomonas cepacia* lipase. Initial testing in 1.5 mL vials at a cell density of 1 g_CDW_/L led to very low conversion of the sterically demanding substrates. Henceforth, the biotransformations were conducted at 3.1 g_CDW_/L in sealed glass vials of 20 mL, leading to increased product formation. To facilitate dissolution of the hydrophobic substrates, stocks were prepared in water-miscible cosolvents, ethanol or DMSO (both 2.5% (v/v)). Interestingly, in biotransformations with DMSO as a cosolvent, we observed further oxidation of the alcohol **9b** to the corresponding aldehyde **9c** and carboxylic acid **9d**, which was further lactonized to **9e** (74% *ee*) under acidic conditions (Figure S14B and S16C-D). The overoxidation is probably caused by alcohol dehydrogenases present in *E. coli* (van Nuland, de Vogel, Eggink, et al., 2017). Contrarily, no overoxidation was observed when ethanol was used as a cosolvent.

**Figure 5.**
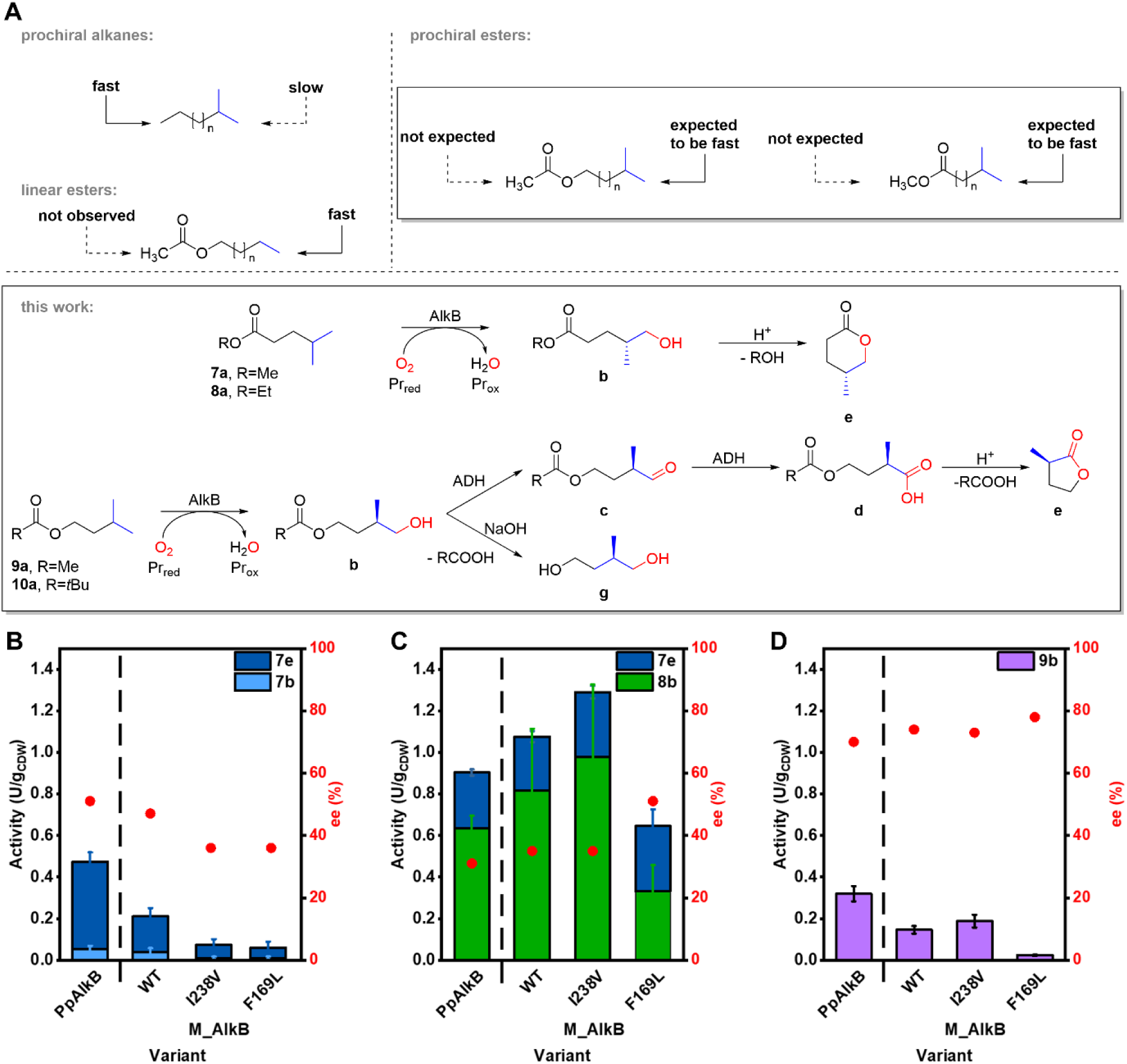
A) Rationale behind the route towards optically pure hydroxy acids, diols, and lactones via protection of the linear terminus with an ester group leading to stereoselective terminal hydroxylation of the prochiral *gem*-dimethyl group by AlkB. Hydroxylation activity and stereoselectivity of PpAlkB and M_AlkB variants performed at 3.1 g_CDW_/L towards the branched esters B) **7a**, C) **8a**, and D) **9a**. The reaction was performed for 24h and data are represented as arithmetic mean ± SD (*n* = 3).

Under optimized reaction conditions, M_AlkB WT showed an activity of 0.27 U/g_CDW_ and 0.15 U/g_CDW_ for the hydroxylation of **7a** and **9a**, which is 1.6-fold and 2.1-fold lower compared to PpAlkB (0.42 and 0.32 U/g_CDW_, respectively). Interestingly, by extending the methyl to an ethyl moiety in **8a**, the activity was higher, reaching 1.08 U/g_CDW_ for M_AlkB and 0.90 U/g_CDW_ for PpAlkB. This could be attributed to the higher logP of **8a** (2.44), compared to **7a** (2.08). Even though an even longer alkyl chain of the alcohol moiety of the ester of branched carboxylic acids might improve activity (van Nuland et al., 2016), a longer group than ethyl might be itself hydroxylated by the enzyme, leading to product mixtures. Therefore, we considered further increasing the size of the ester moiety as a strategy to increase the activity and selectivity. However, the ester **10a** containing a *tert*-butyl group was barely converted (< 2%) (Figure S17), indicating that M_AlkB is rather limited in the conversion of bulky substrates.

Further on, we investigated the influence on the activity and selectivity of several M_AlkB mutants, which increased the enzymatic activity with linear esters (Figure 4). The M_AlkB variant I238V was slightly less active towards **7a**, reaching 0.17 U/g_CDW_, and was more active towards **8a**, reaching 1.29 U/g_CDW_. For **9a**, it displayed comparable activity with a WT of 0.19 U/g_CDW_. Interestingly, the variant M_AlkB F169L, which showed higher activity with the sterically demanding isoprenyl acetate (Nigl et al., 2025) and linear esters, showed only 0.06 U/g_CDW_ for **7a**, while it reached 0.64 U/g_CDW_ for **8a**, still less than the wildtype enzyme. While the phenylalanine to leucine substitution changes size and hydrophobicity,^81^ this observed effect on the activity on the prochiral substrates is difficult to rationalize. Overall, the activity of M_AlkB and its variants was about 20-fold lower towards the sterically hindered esters compared to the conversion of linear esters of the same length, with activities ranging from 0.02 to 1.29 U/g_CDW_. Although activities for all tested AlkB variants were low, using PpAlkB, we produced 0.54 mM of lactone **7e** (starting from 2 mM **7a**) after 24 h (Table 1).

**Table 1.**
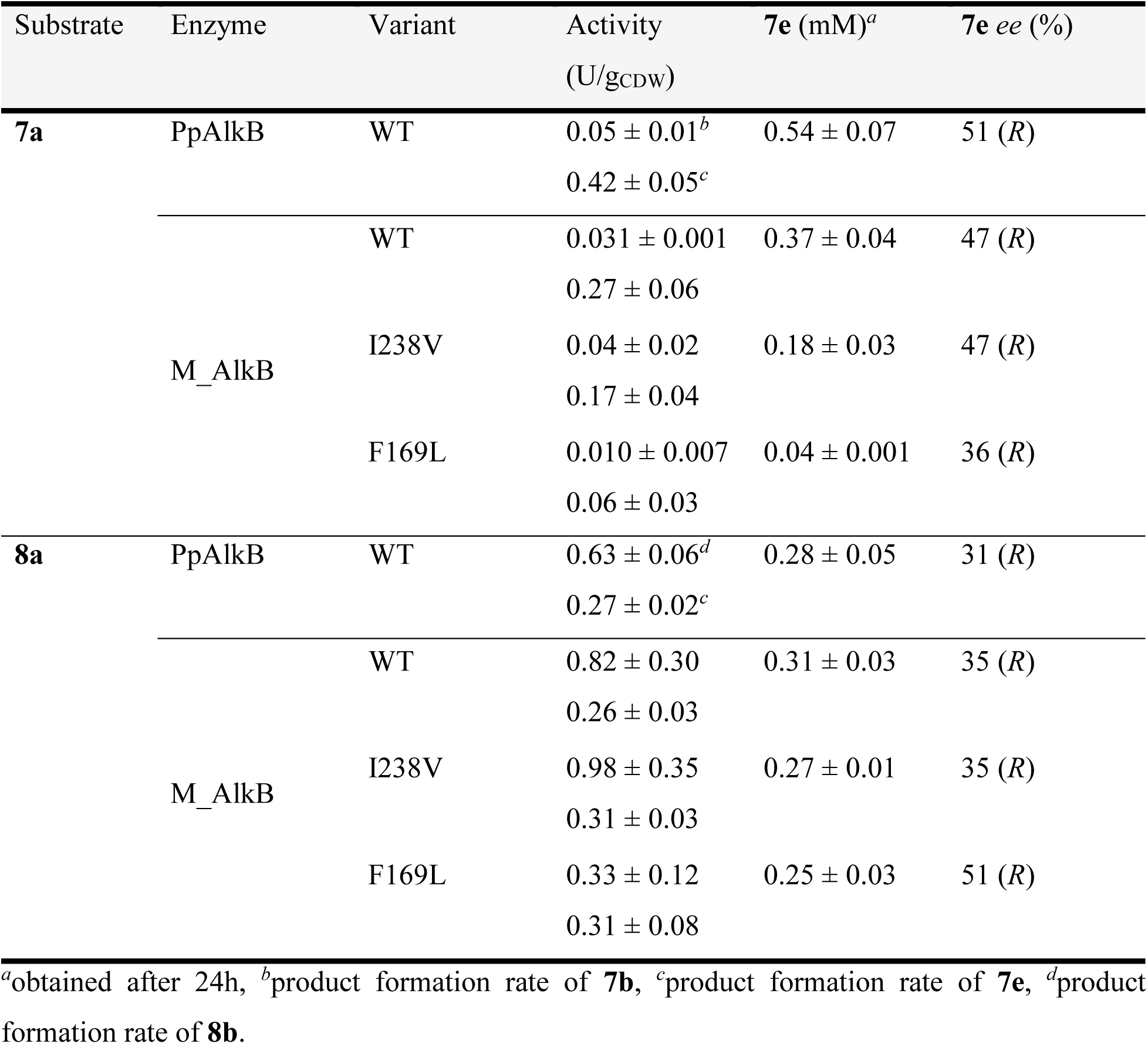
Summary of screening of AlkB WT and variants with the prochiral ester 7a and 8a at 3.1 g_CDW_/L (mean ± SD; *n* = 3).

PpAlkB and M_AlkB displayed similar % *ee* (51 and 47, respectively) with both preferentially forming (*R*)-**7e** lactone (Figure S15 and Table 1). The enantiomeric excess of the lactone **7e** formed by the M_AlkB variants I238V and F169L was comparable with 47% and 36% *ee*, respectively. We were wondering whether prolonging the methyl ester to an ethyl group affects the enantioselectivity. Reactions with **8a** still resulted in the preferred formation of the (*R*)-enantiomer. However, compared to the methyl ester **7a**, the *ee* decreased to 35% for the WT, while for the F169L variant, the *ee* increased to 51% (Table 1). Higher stereoselectivity was observed for M_AlkB-catalyzed hydroxylation of the branched FAlAc **9a**, yielding 79% *ee* of (*R*)**-9b**. Base-catalyzed hydrolysis of **9b** to **9g** slightly reduced the *ee* to 74%, which is similar to results obtained with PpAlkB (81% *ee* (*R*)-**9b** and 74% *ee* (*R*)-**9g**). All tested variants showed similar stereoselectivity in the hydroxylation of **9a** (73-78% *ee*) compared to the WT. Table 1, and Table 2. provide a detailed overview of the activity and stereoselectivity of PpAlkB and M_AlkB wildtype and variants with the branched esters **7a**, **8a**, and **9a**.

**Table 2.** Summary of screening of AlkB WT and variants with the prochiral ester 9a at 3.1 g_CDW_/L (mean ± SD; *n* = 3).

| Substrate | Enzyme | Variant | Activity<br>(U/g<sub>CDW</sub>) | **9b** (mM)<sup>a</sup> | **9b ee**<br>(%) | **9g ee**<br>(%) |
| --- | --- | --- | --- | --- | --- | --- |
| **9a** | PpAlkB | WT | $0.32 \pm 0.04$ | $0.59 \pm 0.02$ | 81 (R) | 74 (R) |
| | M_AlkB | WT | $0.15 \pm 0.02$ | $0.22 \pm 0.02$ | 79 (R) | 74 (R) |
| | | I238V | $0.19 \pm 0.03$ | $0.34 \pm 0.01$ | 79 (R) | 73 (R) |
| | | F169L | $0.02 \pm 0.01$ | $0.04 \pm 0.01$ | 67 (R) | 78 (R) |

<sup>a</sup>obtained after 24h.

Stereoselectivity of PpAlkB was previously shown only for epoxidation of terminal alkenes with *pro*-*R* stereoselectivity (May & Abbott, 1972, 1973), whereas its capacity to discriminate between different alkyl groups of branched hydrocarbons remained unknown. By using a protection strategy, we suppressed the competing hydroxylation of the linear terminus of the substrate and directed the hydroxylating activity towards the prochiral group. PpAlkB and M_AlkB are *pro*-*R* selective in the hydroxylation of the *gem-*dimethyl group, leading to the production of chiral lactone **7e** and chiral diol **9g**. The chiral synthon **7e** is used for the synthesis of active enantiomers of the pheromones of pine sawflies, important for pesticide control (Z. Wang et al., 2007). Interestingly, a previously reported chemosynthetic synthesis of (*R*)-**7e** was based on functionalizing common natural monoterpenoid L-(-)-menthol (Ishmuratov et al., 2004). Our results demonstrate a new biocatalytic route starting from cheap branched esters towards chiral diols and lactones that can be further derivatized into a vast number of chiral building blocks. The obtained results indicated that fatty acid alkyl esters showed better activity than the esters of medium-chain length alcohols. This could be attributed to the difference in the steric hindrance of the methyl ester or acetate group or the difference in hydrophobicity between both. Recent studies proposed the reaction mechanism of alkane monooxygenases for terminal hydroxylation of linear alkanes (Chai et al., 2023; Guo et al., 2023). The molecular mechanism for the stereoselective hydroxylation of prochiral esters is still unknown. Our results indicate two possible binding modes, with *pro*-*R* being preferential. Future efforts will be directed to the identification of amino acid residues directing activity and selectivity towards these substrates and further improving the activity of the monooxygenase by enzyme engineering.

## Conclusion

Alkane monooxygenases show unique selectivity in the terminal hydroxylation of a wide range of substrates. The lack of accurate structural models has so far hampered the engineering of these interesting biocatalysts. Herein, we demonstrated the rational engineering of M_AlkB, isolated from a metagenomic dataset, to improve enzymatic activity towards the synthesis of valuable hydroxylated aliphatic esters. With these results, we gained insight into the substrate preference and stereoselectivity of M_AlkB, an enzyme that appears to be particularly interesting for the conversion of short to medium-chain esters. Using a protection strategy enabled us, for the first time, to characterize the stereoselectivity of alkane monooxygenase towards prochiral terminal *gem*-dimethyl group. This provided a new enzymatic reaction pathway for the synthesis of chiral lactones, hydroxy acids, and diols in good optical purity. We expect that the herein presented results on the substrate scope and mutagenesis of M_AlkB will guide further enzyme engineering studies of bacterial membrane-bound alkane monooxygenases.

## MATERIALS AND METHODS

### Chemicals

If not stated otherwise, all commercially available chemicals and compounds used in this work were purchased either from Merck/Sigma-Aldrich (Darmstadt, Germany), TCI chemicals (Tokyo, Japan), or Carl Roth (Karlsruhe, Germany), usually in the highest purity available. The standard methyl 9-hydroxynonanoate was purchased from Ambeed (Arlington Heights, IL, USA). 5-methyltetrahydropyran-2-one and (5*R*)-tetrahydro-5-methyl-2H-pyran-2-one were purchased from Angene Chemical (Nacharam, India), 2-methylbutane-1,4-diol was purchased from Enamine Ltd. (Kyiv, Ukraine), and (2*R*)-2-methylbutane-1,4-diol was purchased from ABCR (Karlsruhe, Germany).

### Cloning of the *alk*-operon

The plasmid for the expression of *M_alkB* from *Marinobacter* sp. (M_AlkB; NCBI protein: MAB50652.1) was previously prepared by cloning synthetic DNA fragments from IDT into the *pPpalkBFGTL* vector via Gibson Assembly, replacing the *alkB*, while leaving the electron transfer system *alkFGT* from *P. putida* GPo1 intact, as these redox partners proved to be well accepted by the homolog from *Marinobacter* sp. (Nigl et al., 2025). Single and combinatorial mutants of *M_alkB* were either available in-house (Nigl et al., 2025) or generated by site-directed mutagenesis (SDM) (Table S8). Successful mutagenesis was verified by Sanger sequencing (Microsynth, Vienna, Austria). Table S7 lists all the strains used in this study.

### Heterologous protein production and whole-cell biotransformations

*E. coli* BL21(DE3) cells were transformed with the p*M_alkB*(mutant)*FGTL* expression constructs based on the pCom10 backbone, where the expression of the *alkBFGTL* operon is under the control of the natural alkane-inducible promoter *P_alkB_* (Smits et al., 2001). All substrates were tested in whole-cell biotransformation, using *E. coli* BL21(DE3) expressing *M_alkB*(mutant)*FGTL*, similar to previous work (Nigl et al., 2025). Briefly, a single colony of freshly plated *E. coli* containing the respective construct was used to inoculate Lennox lysogeny broth (LB; 5 mL) and incubated overnight at 37 °C with constant agitation (120 rpm). Adapted M9 minimal media (M9MM; 20 mL) (Nigl et al., 2025) was inoculated with seed culture (200 µL) and incubated at 30 °C for 18 to 20 h. For heterologous gene expression, M9MM (200 mL) was inoculated to an OD_600_ of 0.15 with the respective preculture, grown at 30 °C until an OD_600_ of 0.4 to 0.5, and induced by adding 0.05% (v/v) dicyclopropyl ketone (DCPK). After four hours of expression at 30 °C, the cells were harvested (4,400 x *g*, 4 °C, 15 min) and resuspended in resting cell (RC) buffer (50 mM KPi, pH 7.4, 2 mM MgSO4, 1% glucose) to an OD_600_ of 3.2 or 10 (corresponding to 1 g_CDW_/L, and 3.1 g_CDW_/L, respectively). The reaction was initiated by adding 2 mM of substrate (from an 80 mM stock solution in EtOH or DMSO) to the RCs and performed at 25 °C, 180 rpm, with a total volume of either 300 µL in 1.5 mL or 1 mL in 20 mL tightly sealed glass vials. The initial screening of substrates was performed for 24h, while for the quantification, the reactions were performed for 4h and 24h for linear and branched esters, respectively. For each sampling point, a separate reaction was prepared from a master mix. Samples (250 µL) were taken and quenched by adding 2 M HCl (25 µL) and stored at -20 °C until further analysis.

### Gas chromatography (GC) analysis

To analyze biotransformation samples, the quenched samples (275 µL) were extracted with EtOAc (250 µL) containing 1 mM methyl benzoate as internal standard (ISTD) by vigorous shaking for 1 min. After phase separation by centrifugation (16,000 x *g*, 4 °C, 5 min), the organic layer was dried over Na_2_SO_4_, and the extract (200 µL) was directly subjected to GC analysis. To obtain the lactone **9e**, the biotransformation sample (300 µL) was quenched with 2 M HCl (30 µL), centrifuged (16,000 x *g*, 4 °C, 5 min), and the supernatant was incubated for 1 h at 80 °C, 200 rpm in a thermoshaker. 275 µL was then extracted with EtOAc (250 µL) containing ISTD as described. Chiral diols were analyzed by extracting non-quenched biotransformation samples (500 µL) with MTBE (250 µL). After the phases were separated by centrifugation (16,000 x *g*, 4 °C, 5 min), the organic phase was mixed with 10 M NaOH (2.5 µL), and the sample was incubated at 40 °C and 900 rpm for 24 h in a thermoshaker. The sample was dried over Na_2_SO_4_ and directly analyzed by GC.

A GC coupled to a mass spectrometer (GC-MS) was used for qualitative analysis of the reactions and product identification. The measurements were performed on a Shimadzu GC-MS-QP2010 SE (Shimadzu, Kyoto, Japan) equipped with a Zebron ZB-5Plus GC column (30 m × 0.25 mm × 0.25 μm, Phenomenex (Torrance, CA, USA)), using the method presented in Table S9. For quantitative analysis, the extracted reaction samples were analyzed on a GC with a flame ionization detector (GC-FID) from Shimadzu Nexis GC-2030 equipped with a Zebron ZB-5 capillary column (30 m × 0.25 mm × 0.25 μm) from Phenomenex (Torrance, CA, USA). Details on the GC-FID methods used for achiral and chiral analysis are provided in Tables S10-12. Due to the lack of authentic standards for **1b** and **3b**, their hydroxy products and overoxidized products were only tentatively identified by GC-MS. The concentrations of the analytes were calculated by external calibration curves generated by measuring samples with known concentrations of the pure compounds extracted from the RC buffer and treated equally to reaction samples. Enzymatic activities (U/g_CDW_) were obtained from biological triplicates and calculated as µmol of product(s) formed per min and g_CDW_. The data is represented as the arithmetic mean ± standard deviation (SD).

Chiral compounds were analyzed by GC-FID (Shimadzu Nexis GC-2030, Shimadzu, Kyoto, Japan) using either a Hydrodex βTBDAc (50 m × 0.25 mm × 0.25 μm, Macherey-Nagel, GmbH & Co. KG, Düren, Germany) or a Hydrodex βTBDM (25 m × 0.25 mm × 0.25 μm, Macherey-Nagel, GmbH & Co. KG, Düren, Germany) column. Tables S11 and S12 provide detailed descriptions of the methods used.

## SUPPLEMENTARY MATERIAL

Includes additional experimental details, data on modeling and activities, as well as supporting GC chromatograms.

## AUTHOR INFORMATION

### Author Contributions

**Jelena Spasic**: Conceptualization (equal); Formal analysis (lead); Writing – original draft (lead); Writing – review and editing (equal). **Andrea Nigl**: Conceptualization (equal); Formal analysis (lead); Writing – original draft (lead); Writing – review and editing (equal). **Huijin Cheon**: Investigation (equal); Formal analysis (lead); Writing – original draft (lead); Writing – review and editing (equal). **Christine L. Kaiserer**: Investigation (equal); Formal analysis (equal). **Stela Galusic**: Investigation (supporting); Formal analysis (supporting). **Elske van der Pol**: Investigation (supporting); Writing – review and editing (supporting); Methodology (supporting). **Lenny Malihan-Yap**: Investigation (supporting); Writing – review and editing (supporting). **Jin-Byung Park**: Conceptualization (lead); Writing – original draft (supporting); Writing – review and editing (equal). **Robert Kourist**: Conceptualization (lead); Writing – original draft (supporting); Writing – review and editing (equal); Funding acquisition.

## Acknowledgments

This research was funded in part or as whole by the Austrian Science Funds (Project IDs 10.55776/COE17, 10.55776/P34280, 10.55776/P 36614-B). For Open Access purposes, the author has applied a CC BY public copyright license to any author who accepted manuscript version arising from this submission. J.S. and A.N. acknowledge funding from Land Steiermark (PN 44). This project has received funding from the European Union’s Horizon Europe research and innovation program under the Marie Skłodowska-Curie (grant No 101072686). This work was supported by the National Research Foundation (NRF) of Korea funded by the Ministry of Science and ICT (grant No.: 2022M3J4A1066173 and RS-2025-02217886).

## ABBREVIATIONS

AA: amino acid
CDW: cell dry weight
cryo-EM: cryogenic electron microscopy
CYP450: cytochrome P450
DCM: dichloromethane
DCPK: dicyclopropyl ketone
DMAP: 4-dimethylaminopyridine
DMSO: dimethyl sulfoxide
*E. coli*: *Escherichia coli*
EtOAc: ethyl acetate
EtOH: ethanol
FAcE: esters of fatty acids
FAcMe: fatty acid methyl ester
FAlAc: fatty alcohol acetate
FAlE: esters of fatty alcohols
FID: flame ionization detector
FtAlkB: AlkB from *Fontimonas thermophila*
FtAlkBG: fused AlkB AlkG from *Fontimonas thermophila*
GC: gas chromatography
HTS: high-throughput screening
ISTD: internal standard
LB: Lennox lysogeny broth
LPS: Pseudomonas cepacia lipase
M_AlkB: AlkB from *Marinobacter* sp.
M9MM: adapted M9 minimal media
MS: mass spectrometer
NADH: nicotinamide adenine dinucleotide
NMR: nuclear Magnetic Resonance
OD_600_: optical density at 600 nm
PpAlkB: AlkB from *Pseudomonas putida* GPo1
RC: resting cell
ROP: ring opening polymerization
Rt: retention time
SD: standard deviation
SDM: site-directed mutagenesis
WT: wildtype

## REFERENCES

Abramson, J., Adler, J., Dunger, J., Evans, R., Green, T., Pritzel, A., … Jumper, J. M. (2024). Accurate structure prediction of biomolecular interactions with AlphaFold 3. Nature, 630(8016), 493–500. doi: 10.1038/s41586-024-07487-w

Austin, R. N. & Groves, J. T. (2011). Alkane-oxidizing metalloenzymes in the carbon cycle. Metallomics, 3(8), 775. doi: 10.1039/c1mt00048a

Austin, R. N., Luddy, K., Erickson, K., Pender-Cudlip, M., Bertrand, E., Deng, D., … Groves, J. T. (2008). Cage escape competes with geminate recombination during alkane hydroxylation by the diiron oxygenase AlkB. Angew Chem Int Ed, 47(28), 5232–5234. doi: 10.1002/anie.200801184

Behmagham, F., Tawfiq, M. M., Poor Heravi, M. R., Alsultany, N. M. A., Ballal, S., Bahair, H., … Vessally, E. (2024). Advancements in double decarboxylative coupling reactions of carboxylic acids. RSC Adv, 14(21), 14919–14933. doi: 10.1039/D4RA01747A

Chai, J., Guo, G., McSweeney, S. M., Shanklin, J., & Liu, Q. (2023). Structural basis for enzymatic terminal C–H bond functionalization of alkanes. Nat Struct Mol Biol, 30(4), 521–526. doi: 10.1038/s41594-023-00958-0

Choo, J. P. S., Sirota, F. L., See, W. W. L., Eisenhaber, B., & Li, Z. (2023). Engineering of styrene oxide isomerase for enhanced production of 2-arylpropionaldehydes: chemoenzymatic synthesis of (*S*)-profens. ACS Catal, 13(17), 11268–11276. doi: 10.1021/acscatal.3c02777

Dong, J., Fernández-Fueyo, E., Hollmann, F., Paul, C. E., Pesic, M., Schmidt, S., … Zhang, W. (2018). Biocatalytic oxidation reactions: a chemist’s perspective. Angew Chem Int Ed, 57(30), 9238–9261. doi: 10.1002/anie.201800343

Ebrecht, A. C., Aschenbrenner, J. C., Gumulya, Y., Smit, M. S., & Opperman, D. J. (2025). Ancestral sequence reconstruction reveals determinants of regioselectivity in C(sp^3^)-H oxyfunctionalization reactions by CYP505Es. ACS Catal, 15(1), 595–600. doi: 10.1021/acscatal.4c06260

Eggink, G., Lageveen, R. G., Altenburg, B., & Witholt, B. (1987). Controlled and functional expression of the *Pseudomonas oleovorans* alkane utilizing system in *Pseudomonas putida* and *Escherichia coli*. J Biol Chem, 262(36), 17712–17718. doi: 10.1016/S0021-9258(18)45437-3

Engel, J., Cordellier, A., Huang, L., & Kara, S. (2019). Enzymatic ring-opening polymerization of lactones: traditional approaches and alternative strategies. ChemCatChem, 11(20), 4983– 4997. doi: 10.1002/cctc.201900976

Engel, P., Haas, T., Pfeffer, J. C., Thum, O., & Gehring, C. (2015). Patent No. US20150010968A1. United States. Retrieved from https://patents.google.com/patent/US20150010968/ar

Fiorentini, F., Hatzl, A.-M., Schmidt, S., Savino, S., Glieder, A., & Mattevi, A. (2018). The extreme structural plasticity in the CYP153 subfamily of P450s directs development of designer hydroxylases. Biochemistry, 57(48), 6701–6714. doi: 10.1021/acs.biochem.8b01052

Funhoff, E. G., Bauer, U., García-Rubio, I., Witholt, B., & van Beilen, J. B. (2006). CYP153A6, a soluble P450 oxygenase catalyzing terminal-alkane hydroxylation. J Bacteriol, 188(14), 5220–5227. doi: 10.1128/JB.00286-06

Gauthier, M. J., Lafay, B., Christen, R., Fernandez, L., Acquaviva, M., Bonin, P., & Bertrand, J.-C. (1992). *Marinobacter hydrocarbonoclasticus* gen. nov., sp. nov., a new, extremely halotolerant, hydrocarbon-degrading marine bacterium. Int J Syst Bacteriol, 42(4), 568–576. doi: 10.1099/00207713-42-4-568

Goldman, A. S., & Goldberg, K. I. (2004). Organometallic C—H bond activation: an introduction. In K. I. Goldberg & A. S. Goldman (Eds.), Activation and Functionalization of C—H Bonds (Vol. 885, pp. 1–43). Washington, DC: American Chemical Society. doi: 10.1021/bk-2004-0885.ch001

Groves, J. T., Feng, L., & Austin, R. N. (2023). Structure and function of alkane monooxygenase (AlkB). Acc Chem Res, 56(24), 3665–3675. doi: 10.1021/acs.accounts.3c00590

Gubbels, E., Heitz, T., Yamamoto, M., Chilekar, V., Zarbakhsh, S., Gepraegs, M., … Kaminsky, W. (2018). Polyesters. In Ullmann’s Encyclopedia of Industrial Chemistry (pp. 1–30). Wiley. doi: 10.1002/14356007.a21_227.pub2

Gunasekera, T. S., Bowen, L. L., Radwan, O., Striebich, R. C., & Ruiz, O. N. (2022). Genomic and transcriptomic characterization revealed key adaptive mechanisms of *Marinobacter hydrocarbonoclasticus* NI9 for proliferation and degradation of jet fuel. Int Biodeterior Biodegrad, 175, 105502. doi: 10.1016/j.ibiod.2022.105502

Guo, X., Zhang, J., Han, L., Lee, J., Williams, S. C., Forsberg, A., … Feng, L. (2023). Structure and mechanism of the alkane-oxidizing enzyme AlkB. Nat Commun, 14(1), 2180. doi: 10.1038/s41467-023-37869-z

Hammerer, L., Winkler, C. K., & Kroutil, W. (2018). Regioselective biocatalytic hydroxylation of fatty acids by cytochrome P450s. Catal Lett, 148(3), 787–812. doi: 10.1007/s10562-017-2273-4

Head, I. M., Jones, D. M., & Röling, W. F. M. (2006). Marine microorganisms make a meal of oil. Nat Rev Microbiol, 4(3), 173–182. doi: 10.1038/nrmicro1348

Hendricks, L., Reinhardt, C. R., Green, T., Kunczynski, L., Roberts, A. J., Miller, N., … Austin, R. N. (2025). *Fontimonas thermophila* alkane monooxygenase (FtAlkB) is an alkyl fluoride dehalogenase. J Am Chem Soc, 147(11), 9085–9090. doi: 10.1021/jacs.5c00386

Herzog, B., Kohan, M. I., Mestemacher, S. A., Pagilagan, R. U., Redmond, K., & Sarbandi, R. (2020). Polyamides. In Ullmann’s Encyclopedia of Industrial Chemistry (pp. 1–47). Wiley. doi: 10.1002/14356007.a21_179.pub4

Honda Malca, S., Scheps, D., Kühnel, L., Venegas-Venegas, E., Seifert, A., Nestl, B. M., & Hauer, B. (2012). Bacterial CYP153A monooxygenases for the synthesis of omega-hydroxylated fatty acids. Chem Commun, 48(42), 5115. doi: 10.1039/c2cc18103g

Ishmuratov, G. Yu., Yakovleva, M. P., Zaripova, G. V., Botsman, L. P., Muslukhov, R. R., & Tolstikov, G. A. (2004). Novel synthesis of (4*R*)-4-methylpentanolide from (L)-(−)-menthol. Chem Nat Compd, 40(6), 548–551. doi: 10.1007/s10600-005-0033-y

Jelen, J., & Tavčar, G. (2025). Deoxyfluorination: a detailed overview of recent developments. Synthesis, 57(09), 1517–1541. doi: 10.1055/a-2412-1398

Ji, C., Lu, Y., Xia, S., Zhu, C., Zhu, C., Li, W., & Xie, J. (2025). Photoinduced late-stage radical decarboxylative and deoxygenative coupling of complex carboxylic acids and their derivatives. Angew Chem Int Ed, 64(9). doi: 10.1002/anie.202423113

Julsing, M. K., Schrewe, M., Cornelissen, S., Hermann, I., Schmid, A., & Bühler, B. (2012). Outer membrane protein AlkL boosts biocatalytic oxyfunctionalization of hydrophobic substrates in *Escherichia coli*. Appl Environ Microbiol, 78(16), 5724–5733. doi: 10.1128/aem.00949-12

Jumper, J., Evans, R., Pritzel, A., Green, T., Figurnov, M., Ronneberger, O., … Hassabis, D. (2021). Highly accurate protein structure prediction with AlphaFold. Nature, 596(7873), 583–589. doi: 10.1038/s41586-021-03819-2

Kadisch, M., Schmid, A., & Bühler, B. (2017). Hydrolase BioH knockout in *E. coli* enables efficient fatty acid methyl ester bioprocessing. J Ind Microbiol Biotechnol, 44(3), 339–351. doi: 10.1007/s10295-016-1890-z

Khanppnavar, B., Choo, J. P. S., Hagedoorn, P.-L., Smolentsev, G., Štefanić, S., Kumaran, S., … Li, X. (2024). Structural basis of the Meinwald rearrangement catalysed by styrene oxide isomerase. Nat Chem, 16(9), 1496–1504. doi: 10.1038/s41557-024-01523-y

Koch, D. J., Chen, M. M., van Beilen, J. B., & Arnold, F. H. (2009). In vivo evolution of butane oxidation by terminal alkane hydroxylases AlkB and CYP153A6. Appl Environ Microbiol, 75(2), 337–344. doi: 10.1128/aem.01758-08

Kosmadaki, M., & Katsambas, A. (2017). Topical treatments for acne. Clin Dermat, 35(2), 173–178. doi: 10.1016/j.clindermatol.2016.10.010

Ladkau, N., Assmann, M., Schrewe, M., Julsing, M. K., Schmid, A., & Bühler, B. (2016). Efficient production of the nylon 12 monomer ω-aminododecanoic acid methyl ester from renewable dodecanoic acid methyl ester with engineered *Escherichia coli*. Metab Eng, 36, 1–9. doi: 10.1016/j.ymben.2016.02.011

Leech, M. C., & Lam, K. (2020). Electrosynthesis using carboxylic acid derivatives: new tricks for old reactions. Acc Chem Res, 53(1), 121–134. doi: 10.1021/acs.accounts.9b00586

Marques, S. M., Borko, S., Vavra, O., Dvorsky, J., Kohout, P., Kabourek, P., … Bednar, D. (2025). Caver Web 2.0: analysis of tunnels and ligand transport in dynamic ensembles of proteins. Nucleic Acids Res, 53(W1), W132–W142. doi: 10.1093/nar/gkaf399

May, S. W., & Abbott, B. J. (1972). Enzymatic epoxidation I. Alkene epoxidation by the ω-hydroxylation system of *Pseudomonas oleovorans*. Biochem Biophys Res Commun, 48(5), 1230–1234. doi: 10.1016/0006-291X(72)90842-X

May, S. W., & Abbott, B. J. (1973). Enzymatic Epoxidation: II. Comparison between the epoxidation and hydroxylation reactions catalyzed by the ω-hydroxylation system of *Pseudomonas olevorans*. J Biol Chem, 248(5), 1725–1730. doi: 10.1016/s0021-9258(19)44251-8

May, S. W., Gordon, S. L., & Steltenkamp, M. S. (1977). Enzymic epoxidation of trans,trans-1,8-dideuterio-1,7-octadiene. Analysis using partially relaxed proton Fourier transform NMR. J Am Chem Soc, 99(7), 2017–2024. doi: 10.1021/ja00449a001

May, S. W., & Katopodis, A. G. (1986). Oxygenation of alcohol and sulphide substrates by a prototypical non-haem iron monooxygenase: catalysis and biotechnological potential. Enzyme Microb Technol, 8(1), 17–21. doi: 10.1016/0141-0229(86)90004-9

McKenna, E. J., & Coon, M. J. (1970). Enzymatic ω-oxidation: IV purification and properties of the ω-hydroxylase of *Pseudomonas oleovorans*. J Biol Chem, 245(15), 3882–3889. doi: 10.1016/S0021-9258(18)62932-1

Meinhold, P., Peters, M. W., Hartwick, A., Hernandez, A. R., & Arnold, F. H. (2006). Engineering cytochrome P450 BM3 for terminal alkane hydroxylation. Adv Synth Catal, 348(6), 763–772. doi: 10.1002/adsc.200505465

Mikulska-Ruminska, K., Licht, M., Ertem, M. Z., Shanklin, J., Liu, Q., & Bahar, I. (2025, June 27). Substrate binding and channeling allosterically modulate the interactions within the AlkB-AlkG electron transfer complex. BioRxiv, p. 2025.06.23.661152. Cold Spring Harbor Laboratory. doi: 10.1101/2025.06.23.661152

Moon, H. W., Lavagnino, M. N., Lim, S., Palkowitz, M. D., Mandler, M. D., Beutner, G. L., … Radosevich, A. T. (2023). Deoxyfluorination of 1°, 2°, and 3° alcohols by nonbasic O–H activation and Lewis acid-catalyzed fluoride shuttling. J Am Chem Soc, 145(41), 22735–22744. doi: 10.1021/jacs.3c08373

Münch, J., Püllmann, P., Zhang, W., & Weissenborn, M. J. (2021). Enzymatic hydroxylations of sp^3^-carbons. ACS Catal, 11(15), 9168–9203. doi: 10.1021/acscatal.1c00759

Nie, Y., Liang, J., Fang, H., Tang, Y.-Q., & Wu, X.-L. (2011). Two novel alkane hydroxylase-rubredoxin fusion genes isolated from a *Dietzia* Bacterium and the functions of fused rubredoxin domains in long-chain *n*-alkane degradation. Appl Environ Microbiol, 77(20), 7279–7288. doi: 10.1128/aem.00203-11

Nigl, A., Delsoglio, V., Sovic, L., Grgić, M., Malihan-Yap, L., Myrtollari, K., … Kourist, R. (2025). Engineering of transmembrane alkane monooxygenases to improve a key reaction step in the synthesis of polymer precursor tulipalin A. Angew Chem Int Ed, 64(25). doi: 10.1002/anie.202503464

Owen, D. J., Eggink, G., Hauer, B., Kok, M., McBeth, D. L., Yang, Y. L., & Shapiro, J. A. (1984). Physical structure, genetic content and expression of the *alkBAC operon*. Mol Gen Genet, 197(3), 373–383. doi: 10.1007/bf00329932

Peterson, J. A., Basu, D., & Coon, M. J. (1966). Enzymatic ω-oxidation: I: Electron carriers in fatty acid and hydrocarbon hydroxylation. J Biol Chem, 241(21), 5162–5164. doi: 10.1016/S0021-9258(18)99684-5

Peterson, J. A., & Coon, M. J. (1968). Enzymatic ω-oxidation: III. Purification and properties of rubredoxin, a component of the ω-hydroxylation system of *Pseudomonas oleovorans*. J Biol Chem, 243(2), 329–334. doi: 10.1016/S0021-9258(18)99296-3

Ratajczak, A., Geißdörfer, W., & Hillen, W. (1998). Alkane hydroxylase from *Acinetobacter* sp. strain ADP1 is encoded by *alkM* and belongs to a new family of bacterial integral-membrane hydrocarbon hydroxylases. Appl Environ Microbiol, 64(4), 1175–1179. doi: 10.1128/aem.64.4.1175-1179.1998

Reinhardt, C. R., Lee, J. A., Hendricks, L., Green, T., Kunczynski, L., Roberts, A. J., … Austin, R. N. (2025). No bridge between us: EXAFS and computations confirm two distant iron ions comprise the active site of alkane monooxygenase (AlkB). J Am Chem Soc, 147(3), 2432– 2443. doi: 10.1021/jacs.4c12633

Rivière, S., Vielmuth, C., Ennenbach, C., Abdelrahman, A., Lemke, C., Gütschow, M., … Menche, D. (2020). Design, synthesis and biological evaluation of highly potent simplified archazolids. ChemMedChem, 15(14), 1348–1363. doi: 10.1002/cmdc.202000154

Schaffer, S., & Haas, T. (2014). Biocatalytic and fermentative production of α,ω-bifunctional polymer precursors. Org Process Res Dev, 18(6), 752–766. doi: 10.1021/op5000418

Schrewe, M., Julsing, M. K., Lange, K., Czarnotta, E., Schmid, A., & Bühler, B. (2014). Reaction and catalyst engineering to exploit kinetically controlled whole-cell multistep biocatalysis for terminal FAME oxyfunctionalization. Biotechnol Bioeng, 111(9), 1820–1830. doi: 10.1002/bit.25248

Schrewe, M., Ladkau, N., Bühler, B., & Schmid, A. (2013). Direct terminal alkylamino-functionalization *via* multistep biocatalysis in one recombinant whole-cell catalyst. Adv Synth Catal, 355(9), 1693–1697. doi: 10.1002/adsc.201200958

Schrewe, M., Magnusson, A. O., Willrodt, C., Bühler, B., & Schmid, A. (2011). Kinetic analysis of terminal and unactivated C‒H bond oxyfunctionalization in fatty acid methyl esters by monooxygenase-based whole-cell biocatalysis. Adv Synth Catal, 353(18), 3485–3495. doi: 10.1002/adsc.201100440

Schultes, F. P. J., Welter, L., Schmidtke, M., Tischler, D., & Mügge, C. (2024). A tailored cytochrome P450 monooxygenase from *Gordonia rubripertincta* CWB2 for selective aliphatic monooxygenation. Biol Chem, 405(9–10), 677–689. doi: 10.1515/hsz-2024-0041

Siedlecka, R. (2023). Selectivity in the aliphatic C–H bonds oxidation (hydroxylation) catalyzed by heme-and non-heme metal complexes—recent advances. Catalysts, 13(1), 121. doi: 10.3390/catal13010121

Smits, T. H. M., Balada, S. B., Witholt, B., & van Beilen, J. B. (2002). Functional analysis of alkane hydroxylases from Gram-negative and Gram-positive bacteria. J Bacteriol, 184(6), 1733–1742. doi: 10.1128/JB.184.6.1733-1742.2002

Smits, T. H. M., Seeger, M. A., Witholt, B., & van Beilen, J. B. (2001). New alkane-responsive expression vectors for *Escherichia coli* and *Pseudomonas*. Plasmid, 46(1), 16–24. doi: 10.1006/plas.2001.1522

Spanjers, C. S., Schneiderman, D. K., Wang, J. Z., Wang, J., Hillmyer, M. A., Zhang, K., & Dauenhauer, P. J. (2016). Branched diol monomers from the sequential hydrogenation of renewable carboxylic acids. ChemCatChem, 8(19), 3031–3035. doi: 10.1002/cctc.201600710

Suzuki, T., Usui, K., Miyake, Y., Namikoshi, M., & Nakada, M. (2004). First total synthesis of antimitotic compound, (+)-phomopsidin. Org Lett, 6(4), 553–556. doi: 10.1021/ol036338q

Takahashi, K., Komine, K., Yokoi, Y., Ishihara, J., & Hatakeyama, S. (2012). Stereocontrolled total synthesis of (−)-englerin A. J Org Chem, 77(17), 7364–7370. doi: 10.1021/jo301145r

Tully, B. J., Graham, E. D., & Heidelberg, J. F. (2018). The reconstruction of 2,631 draft metagenome-assembled genomes from the global oceans. Sci Data, 5(1), 170203. doi: 10.1038/sdata.2017.203

van Beilen, J. B., Kingma, J., & Witholt, B. (1994). Substrate specificity of the alkane hydroxylase system of *Pseudomonas oleovorans* GPo1. Enzyme Microb Technol, 16(10), 904–911. doi: 10.1016/0141-0229(94)90066-3

Van Beilen, J. B., Marín, M. M., Smits, T. H. M., Röthlisberger, M., Franchini, A. G., Witholt, B., & Rojo, F. (2004). Characterization of two alkane hydroxylase genes from the marine hydrocarbonoclastic bacterium *Alcanivorax borkumensis*. Environ Microbiol, 6(3), 264–273. doi: 10.1111/j.1462-2920.2004.00567.x

van Beilen, J. B., Panke, S., Lucchini, S., Franchini, A. G., Röthlisberger, M., & Witholt, B. (2001). Analysis of *Pseudomonas putida* alkane-degradation gene clusters and flanking insertion sequences: evolution and regulation of the *alk* genes. Microbiology, 147(6), 1621–1630. doi: 10.1099/00221287-147-6-1621

van Beilen, J. B., Smits, T. H. M., Roos, F. F., Brunner, T., Balada, S. B., Röthlisberger, M., & Witholt, B. (2005). Identification of an amino acid position that determines the substrate range of integral membrane alkane hydroxylases. J Bacteriol, 187(1), 85–91. doi: 10.1128/jb.187.1.85-91.2005

Van Beilen, J. B., Smits, T. H. M., Whyte, L. G., Schorcht, S., Röthlisberger, M., Plaggemeier, T., … Witholt, B. (2002). Alkane hydroxylase homologues in Gram-positive strains. Environ Microbiol, 4(11), 676–682. doi: 10.1046/j.1462-2920.2002.00355.x

van Beilen, J. B., Wubbolts, M. G., & Witholt, B. (1994). Genetics of alkane oxidation by *Pseudomonas oleovorans*. Biodegradation, 5(3–4), 161–174. doi: 10.1007/bf00696457

van Nuland, Y. M., de Vogel, F. A., Eggink, G., & Weusthuis, R. A. (2017). Expansion of the ω- oxidation system AlkBGTL of *Pseudomonas putida* GPo1 with AlkJ and AlkH results in exclusive mono-esterified dicarboxylic acid production in *E. coli*. Microb Biotechnol, 10(3), 594–603. doi: 10.1111/1751-7915.12607

van Nuland, Y. M., de Vogel, F. A., Scott, E. L., Eggink, G., & Weusthuis, R. A. (2017). Biocatalytic, one-pot diterminal oxidation and esterification of n-alkanes for production of α,ω-diol and α,ω-dicarboxylic acid esters. Metab Eng, 44, 134–142. doi: 10.1016/j.ymben.2017.10.005

van Nuland, Y. M., Eggink, G., & Weusthuis, R. A. (2016). Application of AlkBGT and AlkL from *Pseudomonas putida* GPo1 for selective alkyl ester ω-oxyfunctionalization in *Escherichia coli*. Appl Environ Microbiol, 82(13), 3801–3807. doi: 10.1128/aem.00822-16

Vomberg, A., & Klinner, U. (2000). Distribution of alkB genes within *n*-alkane-degrading bacteria. J Appl Microbiol, 89(2), 339–348. doi: 10.1046/j.1365-2672.2000.01121.x

Wang, Y., & Liu, Y. (2024). Computational insights into the non-heme diiron alkane monooxygenase enzyme AlkB: electronic structures, dioxygen activation, and hydroxylation mechanism of liquid alkanes. Inorg Chem, 63(37), 17056–17066. doi: 10.1021/acs.inorgchem.4c02721

Wang, Z., Xu, Q., Tian, W., & Pan, X. (2007). Stereoselective synthesis of (2*S*,3*S*,7*S*)-3,7-dimethylpentadec-2-yl acetate and propionate, the sex pheromones of pine sawflies. Tetrahedron Lett, 48(42), 7549–7551. doi: 10.1016/j.tetlet.2007.06.123

White, M. C. (2012). Adding aliphatic C–H bond oxidations to synthesis. Science, 335(6070), 807–809. doi: 10.1126/science.1207661

Williams, S. C., & Austin, R. N. (2022). An overview of the electron-transfer proteins that activate alkane monooxygenase (AlkB). Front Microbiol, 13. doi: 10.3389/fmicb.2022.845551

Williams, S. C., Forsberg, A. P., Lee, J., Vizcarra, C. L., Lopatkin, A. J., & Austin, R. N. (2021). Investigation of the prevalence and catalytic activity of rubredoxin-fused alkane monooxygenases (AlkBs). J Inorg Biochem, 219, 111409. doi: 10.1016/j.jinorgbio.2021.111409

Williams, S. C., Luongo, D., Orman, M., Vizcarra, C. L., & Austin, R. N. (2022). An alkane monooxygenase (AlkB) family in which all electron transfer partners are covalently bound to the oxygen-activating hydroxylase. J Inorg Biochem, 228, 111707. doi: 10.1016/j.jinorgbio.2021.111707

Wu, Y., Paul, C. E., & Hollmann, F. (2023). Mirror, mirror on the wall, which is the greenest of them all? A critical comparison of chemo- and biocatalytic oxyfunctionalisation reactions. Green Carbon, 1(2), 227–241. doi: 10.1016/j.greenca.2023.10.004

Xie, Y., Ramirez, D., Chen, G., He, G., Sun, Y., Murdoch, F. K., & Löffler, F. E. (2023). Genome-wide expression analysis unravels fluoroalkane metabolism in *Pseudomonas* sp. strain 273. Environ Sci Technol, 57(42), 15925–15935. doi: 10.1021/acs.est.3c03855

Yang, Y., Wang, J., Liao, J., Xie, S., & Huang, Y. (2015). Abundance and diversity of soil petroleum hydrocarbon-degrading microbial communities in oil exploring areas. Appl Microbiol Biotechnol, 99(4), 1935–1946. doi: 10.1007/s00253-014-6074-z

