## Supplemental information for "Engineering Membrane-Bound Alkane Monooxygenase from *Marinobacter* sp. for Increased Activity in the Selective ω-Hydroxylation of Linear and Branched Aliphatic Esters"

AUTHOR ADDRESS

### 1 Computational methods

#### 1.1 AlkB protein sequence comparisons

Table S1 provides an amino acid distribution according to the 3DM database (Kuipers et al., 2010) of the residues selected for mutation in M_AlkB (W60, F169 and I238). The M_AlkB protein sequence was aligned with FtAlkB (Guo et al., 2023), FtAlkBG (Chai, Guo, McSweeney, Shanklin, & Liu, 2023), and PpAlkB using Jalview (Figure S1) (Waterhouse, Procter, Martin, Clamp, & Barton, 2009) and the sequence homology was calculated using the web tool Expasy SIM (Expasy - SIM Alignment Tool; Gasteiger, 2003). For homology calculations, only the monooxygenase part of FtAlkBG was taken into account.

**Table S1.** Amino acid distribution for selected positions based on data retrieved from the 3DM database (Kuipers et al., 2010). Alignment of 16,104 sequences (18 sequences with gaps).

| Position | Amino acid occurrence (%) |
|---|---|
| **60** | Valine (28.01), Leucine (18.96), Alanine (16.98), Isoleucine (13.65), Threonine (7.90), Tryptophan (1.48), others (13.12) |
| **169** | Phenylalanine (98.97), Tryptophan (0.35), Tyrosine (0.29), others (0.39) |
| **238** | Alanine (37.78), Leucine (28.85), Glycine (8.87), Isoleucine (6.71), Serine (5.61), others (12.64) |

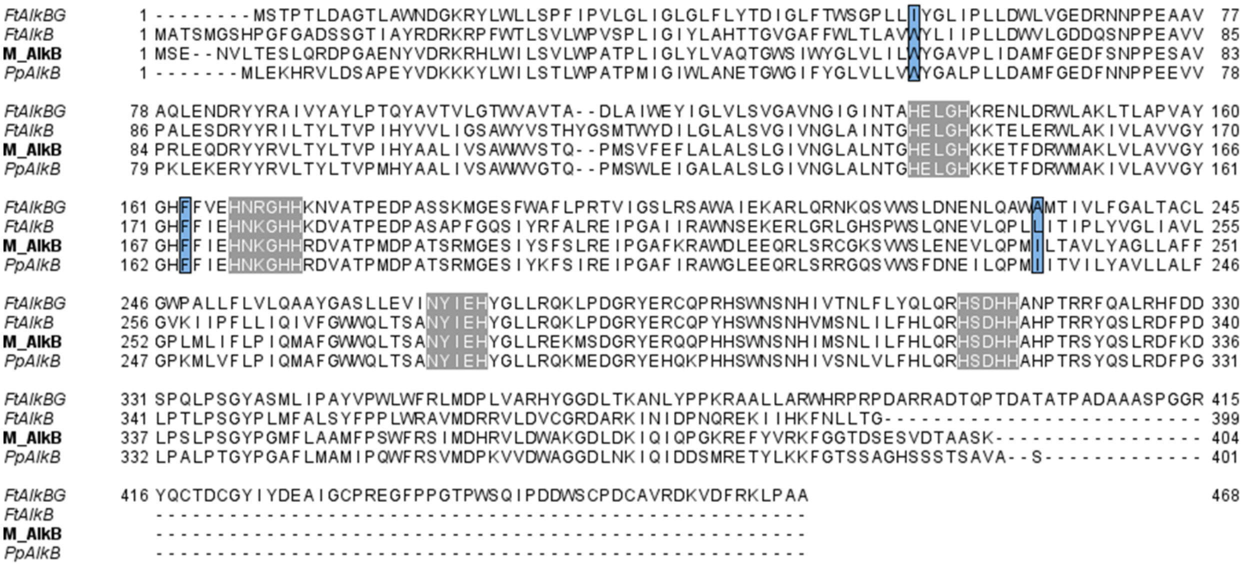

**Figure S1.** Multiple sequence alignment of M_AlkB, FtAlkB, FtAlkBG, and PpAlkB. The grey boxes highlight the four highly conserved sequence motifs containing the nine histidine residues coordinating the Fe-atoms in the active site of alkane monooxygenases (van Beilen et al., 2005). The three residues investigated in this work are highlighted in blue.

#### 1.2 AlkB protein structure prediction and comparison

3D models of AlkB from *Marinobacter sp*. (M_AlkB) and *Pseudomonas putida* GPo1 (recently renamed to *Ectopseudomonas oleovorans*, formerly *Pseudomonas oleovorans*; PpAlkB) were predicted using AlphaFold 3 (AF3) with two $Fe^{3+}$ ions set as ligands (Abramson et al., 2024). The structures were refined with the Protein Preparation Wizard module in Schrödinger Maestro 14.2 (Schrödinger LLC, 2025), minimizing only hydrogen atoms using the OPLS3e force field (RMSD was set to 0.3 Å). The predicted structures were compared with each other and the published cryo-EM structures of FtAlkB (PDB 8SBB) and FtAlkBG (PDB 8F6T), as shown in Figure S2. Putative tunnels were predicted with the online tool CAVER web 2.0 (https://loschmidt.chemi.muni.cz/caverweb/; (Marques et al., 2025; Stourac et al., 2019) using the minimized structures as PDB source files. Since the catalytic mechanism of AlkB has not been fully established, two starting points were compared: (i) the midpoint between the two Fe ions and (ii) the Fe ion reported to be involved in the oxidation reaction (Groves, Feng, & Austin, 2023). The midpoint was set by entering the direct coordinates and the Fe-atom coordinated by four histidines (Fe1; chain C) was selected as the ligand. Standard amino acids and Fe-atoms were included in the tunnel detection. Default molecular dynamics settings were used (CAVER parameters: minimum probe radius: 0.9, shell depth: 4, radius: 3, clustering threshold: 3.5, max. distance: 3, desired radius: 5; YASARA parameters: density: 0.997, pH: 7.4, temperature: 298. K, duration: 5000 ps, save interval: 10 ps, snapshots: 500). All predicted tunnels of PpAlkB (Table S1) and M_AlkB (Table S2) were analyzed and compared to the cryo-EM structures of FtAlkB and FtAlkBG (Figure S2). The top five tunnels for both enzymes and starting points are highlighted in the tables and visualized in Figure S3. Upon analysis, tunnel 5, both in PpAlkB and M_AlkB, starting from the midpoint of the iron ions showed the highest resemblance to the substrate tunnel described by Guo et al., 2023 and were used for further analysis and visualizations. The analysis starting from the Fe-atom gave similar results (Table S2-S3 and Figure S3).

**Table S2.** Summary of predicted tunnels for PpAlkB using CAVER (Marques et al., 2025; Stourac et al., 2019). The highlighted tunnels are presented in Figure S3A.

| ID | No | No_snaps | Avg_BR | SD | Max_BR | Avg_L | SD | Avg_C | SD | Priority | Avg_throughput | SD |
|---|---|---|---|---|---|---|---|---|---|---|---|---|
| **Midpoint of Fe-atoms as a starting point** | | | | | | | | | | | | |
| 1 | 440 | 440 | 1.120 | 0.111 | 1.43 | 20.103 | 1.763 | 1.575 | 0.073 | 0.38263 | 0.43568 | 0.03166 |
| 2 | 429 | 429 | 1.066 | 0.144 | 1.67 | 21.976 | 2.227 | 1.261 | 0.080 | 0.33412 | 0.39019 | 0.06726 |
| 3 | 122 | 122 | 0.974 | 0.057 | 1.15 | 20.151 | 1.875 | 1.420 | 0.090 | 0.08834 | 0.36279 | 0.04632 |
| 4 | 50 | 50 | 0.979 | 0.059 | 1.15 | 8.416 | 1.774 | 1.134 | 0.078 | 0.06023 | 0.60346 | 0.04121 |
| 5 | 36 | 36 | 0.972 | 0.068 | 1.13 | 27.787 | 2.446 | 1.334 | 0.118 | 0.02354 | 0.32760 | 0.06288 |
| 6 | 31 | 31 | 0.927 | 0.026 | 0.99 | 23.045 | 2.900 | 1.791 | 0.245 | 0.01942 | 0.31385 | 0.05340 |
| 7 | 32 | 32 | 0.947 | 0.049 | 1.13 | 28.555 | 1.752 | 1.354 | 0.051 | 0.01742 | 0.27267 | 0.04442 |
| 8 | 29 | 29 | 0.924 | 0.022 | 0.97 | 30.437 | 1.868 | 1.502 | 0.105 | 0.01040 | 0.17970 | 0.02970 |
| 9 | 17 | 17 | 0.944 | 0.051 | 1.10 | 32.432 | 2.507 | 1.816 | 0.154 | 0.00698 | 0.20561 | 0.04324 |
| 10 | 9 | 9 | 0.920 | 0.019 | 0.97 | 31.667 | 4.943 | 1.752 | 0.344 | 0.00316 | 0.17613 | 0.06267 |
| 11 | 4 | 4 | 0.918 | 0.021 | 0.95 | 18.326 | 0.097 | 1.265 | 0.029 | 0.00262 | 0.32765 | 0.02347 |
| 12 | 4 | 4 | 0.918 | 0.010 | 0.93 | 30.353 | 0.691 | 1.303 | 0.034 | 0.00140 | 0.17595 | 0.01701 |
| 13 | 2 | 2 | 0.919 | 0.012 | 0.93 | 32.334 | 2.084 | 1.386 | 0.100 | 0.00083 | 0.20747 | 0.03390 |
| 14 | 2 | 2 | 0.914 | 0.012 | 0.93 | 31.964 | 0.853 | 1.836 | 0.058 | 0.00075 | 0.18811 | 0.01488 |
| 15 | 2 | 2 | 0.963 | 0.061 | 1.02 | 37.608 | 2.343 | 1.754 | 0.129 | 0.00072 | 0.18117 | 0.01269 |
| 16 | 1 | 1 | 0.964 | 0.000 | 0.96 | 30.816 | 0.000 | 1.312 | 0.000 | 0.00050 | 0.25233 | 0.00000 |
| 17 | 1 | 1 | 0.909 | 0.000 | 0.91 | 37.627 | 0.000 | 1.814 | 0.000 | 0.00024 | 0.12260 | 0.00000 |
| 18 | 1 | 1 | 0.935 | 0.000 | 0.93 | 50.613 | 0.000 | 2.241 | 0.000 | 0.00020 | 0.09807 | 0.00000 |
| **Fe1 (Chain C) set as a starting point** | | | | | | | | | | | | |
| 1 | 429 | 429 | 1.071 | 0.154 | 1.69 | 18.94 | 2.234 | 1.257 | 0.079 | 0.36665 | 0.42819 | 0.07519 |
| 2 | 440 | 440 | 1.12 | 0.111 | 1.43 | 22.435 | 1.929 | 1.537 | 0.071 | 0.3652 | 0.41583 | 0.03294 |
| 3 | 122 | 122 | 0.974 | 0.057 | 1.15 | 17.473 | 2.192 | 1.384 | 0.104 | 0.09594 | 0.39398 | 0.05371 |
| 4 | 50 | 50 | 0.979 | 0.06 | 1.15 | 10.757 | 1.851 | 1.182 | 0.071 | 0.05548 | 0.55593 | 0.04264 |
| 5 | 36 | 36 | 0.972 | 0.068 | 1.13 | 26.814 | 2.932 | 1.402 | 0.147 | 0.02418 | 0.33644 | 0.06444 |
| 6 | 32 | 32 | 0.928 | 0.027 | 0.99 | 22.204 | 2.805 | 1.651 | 0.229 | 0.02074 | 0.32467 | 0.05499 |
| 7 | 32 | 32 | 0.947 | 0.049 | 1.13 | 27.384 | 2.057 | 1.314 | 0.065 | 0.01806 | 0.28277 | 0.04308 |
| 8 | 29 | 29 | 0.924 | 0.022 | 0.97 | 30.189 | 2.083 | 1.526 | 0.108 | 0.01068 | 0.18457 | 0.02972 |
| 9 | 17 | 17 | 0.944 | 0.051 | 1.10 | 34.912 | 2.807 | 1.75 | 0.138 | 0.00661 | 0.19494 | 0.04123 |
| 10 | 4 | 4 | 0.918 | 0.021 | 0.95 | 23.409 | 1.219 | 1.439 | 0.039 | 0.00231 | 0.28984 | 0.0171 |

| | | | | | | | | | | | | |
|---|---|---|---|---|---|---|---|---|---|---|---|---|
| 11 | 8 | 8 | 0.914 | 0.009 | 0.93 | 34.663 | 2.651 | 1.851 | 0.143 | 0.00229 | 0.14312 | 0.01 523 |
| 12 | 4 | 4 | 0.918 | 0.01 | 0.93 | 32.274 | 1.264 | 1.352 | 0.016 | 0.00135 | 0.1686 | 0.01 963 |
| 13 | 2 | 2 | 0.919 | 0.012 | 0.93 | 30.838 | 0.548 | 1.393 | 0.063 | 0.00086 | 0.21489 | 0.02 43 |
| 14 | 1 | 1 | 0.964 | 0.000 | 0.96 | 33.302 | 0.000 | 1.287 | 0.000 | 0.00048 | 0.24222 | 0.00 000 |
| 15 | 1 | 1 | 0.903 | 0.000 | 0.90 | 38.676 | 0.000 | 1.972 | 0.000 | 0.00035 | 0.17511 | 0.00 000 |
| 16 | 1 | 1 | 0.911 | 0.000 | 0.91 | 30.42 | 0.000 | 1.463 | 0.000 | 0.00032 | 0.15844 | 0.00 000 |
| 17 | 1 | 1 | 0.909 | 0.000 | 0.91 | 34.637 | 0.000 | 1.871 | 0.000 | 0.00027 | 0.13707 | 0.00 000 |
| 18 | 1 | 1 | 0.935 | 0.000 | 0.93 | 46.172 | 0.000 | 2.105 | 0.000 | 0.00023 | 0.11691 | 0.00 000 |

**Table S3.** Summary of predicted tunnels for M_AlkB using CAVER (Marques et al., 2025; Stourac et al., 2019). The highlighted tunnels are presented in Figure S3B.

| ID | No | Nosnaps | Avg_BR | SD | Max_BR | Avg_L | SD | Avg_C | SD | Priority | Avg_throughput | SD |
|---|---|---|---|---|---|---|---|---|---|---|---|---|
| **Midpoint of Fe-atoms as a starting point** | | | | | | | | | | | | |
| 1 | 458 | 458 | 1.111 | 0.122 | 1.44 | 20.681 | 2.562 | 1.494 | 0.236 | 0.37422 | 0.40935 | 0.05573 |
| 2 | 184 | 184 | 0.989 | 0.085 | 1.38 | 22.864 | 2.371 | 1.321 | 0.149 | 0.13374 | 0.36414 | 0.06974 |
| 3 | 154 | 154 | 0.969 | 0.056 | 1.15 | 20.637 | 2.431 | 1.367 | 0.093 | 0.11381 | 0.37025 | 0.05699 |
| 4 | 93 | 93 | 0.968 | 0.065 | 1.24 | 24.535 | 3.546 | 1.802 | 0.298 | 0.05999 | 0.32316 | 0.07337 |
| 5 | 47 | 47 | 0.960 | 0.049 | 1.15 | 28.285 | 2.879 | 1.409 | 0.119 | 0.02736 | 0.29168 | 0.04893 |
| 6 | 6 | 6 | 0.946 | 0.048 | 1.05 | 31.653 | 6.667 | 1.994 | 0.381 | 0.00292 | 0.24400 | 0.08979 |
| 7 | 4 | 4 | 0.914 | 0.014 | 0.94 | 33.639 | 1.641 | 1.497 | 0.188 | 0.00144 | 0.18098 | 0.02751 |
| 8 | 2 | 2 | 0.939 | 0.005 | 0.94 | 26.111 | 3.130 | 2.417 | 0.022 | 0.00122 | 0.30451 | 0.03075 |
| 9 | 4 | 4 | 0.912 | 0.003 | 0.91 | 33.284 | 1.750 | 1.596 | 0.034 | 0.00111 | 0.13854 | 0.01263 |
| 10 | 1 | 1 | 0.914 | 0.000 | 0.91 | 9.085 | 0.000 | 1.220 | 0.000 | 0.00101 | 0.50639 | 0.00000 |
| 11 | 1 | 1 | 0.932 | 0.000 | 0.93 | 11.970 | 0.000 | 1.318 | 0.000 | 0.00095 | 0.47346 | 0.00000 |
| 12 | 2 | 2 | 0.903 | 0.003 | 0.91 | 37.557 | 1.194 | 1.880 | 0.025 | 0.00057 | 0.14279 | 0.01900 |
| 13 | 2 | 2 | 0.907 | 0.005 | 0.91 | 42.286 | 2.647 | 2.169 | 0.073 | 0.00046 | 0.11495 | 0.01187 |
| 14 | 1 | 1 | 0.902 | 0.000 | 0.90 | 28.387 | 0.000 | 1.456 | 0.000 | 0.00042 | 0.21117 | 0.00000 |
| 15 | 1 | 1 | 0.923 | 0.000 | 0.92 | 35.534 | 0.000 | 1.977 | 0.000 | 0.00038 | 0.18833 | 0.00000 |
| 16 | 1 | 1 | 0.973 | 0.000 | 0.97 | 34.523 | 0.000 | 1.423 | 0.000 | 0.00036 | 0.18244 | 0.00000 |
| 17 | 1 | 1 | 0.945 | 0.000 | 0.95 | 33.797 | 0.000 | 1.457 | 0.000 | 0.00033 | 0.16540 | 0.00000 |
| 18 | 1 | 1 | 0.911 | 0.000 | 0.91 | 37.299 | 0.000 | 1.507 | 0.000 | 0.00026 | 0.13090 | 0.00000 |
| 19 | 1 | 1 | 1.027 | 0.000 | 1.03 | 52.299 | 0.000 | 2.781 | 0.000 | 0.00021 | 0.10281 | 0.00000 |
| 20 | 1 | 1 | 0.938 | 0.000 | 0.94 | 62.858 | 0.000 | 2.909 | 0.000 | 0.00011 | 0.05606 | 0.00000 |
| **Fe1 (Chain C) set as a starting point** | | | | | | | | | | | | |
| 1 | 461 | 461 | 1.138 | 0.131 | 1.51 | 22.813 | 2.012 | 1.525 | 0.092 | 0.36406 | 0.39565 | 0.05435 |
| 2 | 185 | 185 | 0.991 | 0.086 | 1.38 | 18.112 | 1.849 | 1.236 | 0.071 | 0.15733 | 0.42606 | 0.0671 |
| 3 | 155 | 155 | 0.975 | 0.061 | 1.17 | 15.01 | 1.795 | 1.302 | 0.092 | 0.13962 | 0.4513 | 0.06407 |
| 4 | 96 | 96 | 0.968 | 0.064 | 1.24 | 22.928 | 3.221 | 1.82 | 0.252 | 0.06682 | 0.34869 | 0.07751 |
| 5 | 50 | 50 | 0.962 | 0.048 | 1.11 | 27.384 | 2.537 | 1.433 | 0.13 | 0.03063 | 0.30691 | 0.05006 |
| 6 | 4 | 4 | 0.912 | 0.003 | 0.91 | 28.876 | 1.795 | 1.471 | 0.058 | 0.00132 | 0.16569 | 0.01043 |
| 7 | 3 | 3 | 0.906 | 0.005 | 0.91 | 31.507 | 1.717 | 1.302 | 0.099 | 0.00125 | 0.20955 | 0.03393 |
| 8 | 1 | 1 | 0.932 | 0.000 | 0.93 | 12.264 | 0.000 | 1.294 | 0.000 | 0.00097 | 0.48347 | 0.00000 |
| 9 | 1 | 1 | 0.914 | 0.000 | 0.91 | 10.773 | 0.000 | 1.128 | 0.000 | 0.00079 | 0.39767 | 0.00000 |
| 10 | 2 | 2 | 0.942 | 0.004 | 0.95 | 31.685 | 0.043 | 1.439 | 0.034 | 0.00067 | 0.16900 | 0.01455 |
| 11 | 2 | 2 | 0.903 | 0.003 | 0.91 | 36.712 | 1.280 | 2.072 | 0.136 | 0.0006 | 0.15056 | 0.02181 |
| 12 | 1 | 1 | 0.981 | 0.000 | 0.98 | 31.373 | 0.000 | 1.422 | 0.000 | 0.00047 | 0.2336 | 0.00000 |
| 13 | 2 | 2 | 0.907 | 0.005 | 0.91 | 43.674 | 1.557 | 1.995 | 0.021 | 0.00046 | 0.11512 | 0.00991 |
| 14 | 1 | 1 | 0.923 | 0.000 | 0.92 | 31.553 | 0.000 | 2.252 | 0.000 | 0.00043 | 0.21313 | 0.00000 |
| 15 | 1 | 1 | 0.902 | 0.000 | 0.9 | 35.667 | 0.000 | 1.609 | 0.000 | 0.00031 | 0.15637 | 0.00000 |
| 16 | 1 | 1 | 0.9 | 0.000 | 0.9 | 36.126 | 0.000 | 1.516 | 0.000 | 0.00023 | 0.11460 | 0.00000 |
| 17 | 1 | 1 | 1.027 | 0.000 | 1.03 | 51.610 | 0.000 | 2.834 | 0.000 | 0.00021 | 0.10547 | 0.00000 |
| 18 | 1 | 1 | 0.938 | 0.000 | 0.94 | 62.127 | 0.000 | 3.05 | 0.000 | 0.00011 | 0.05750 | 0.00000 |

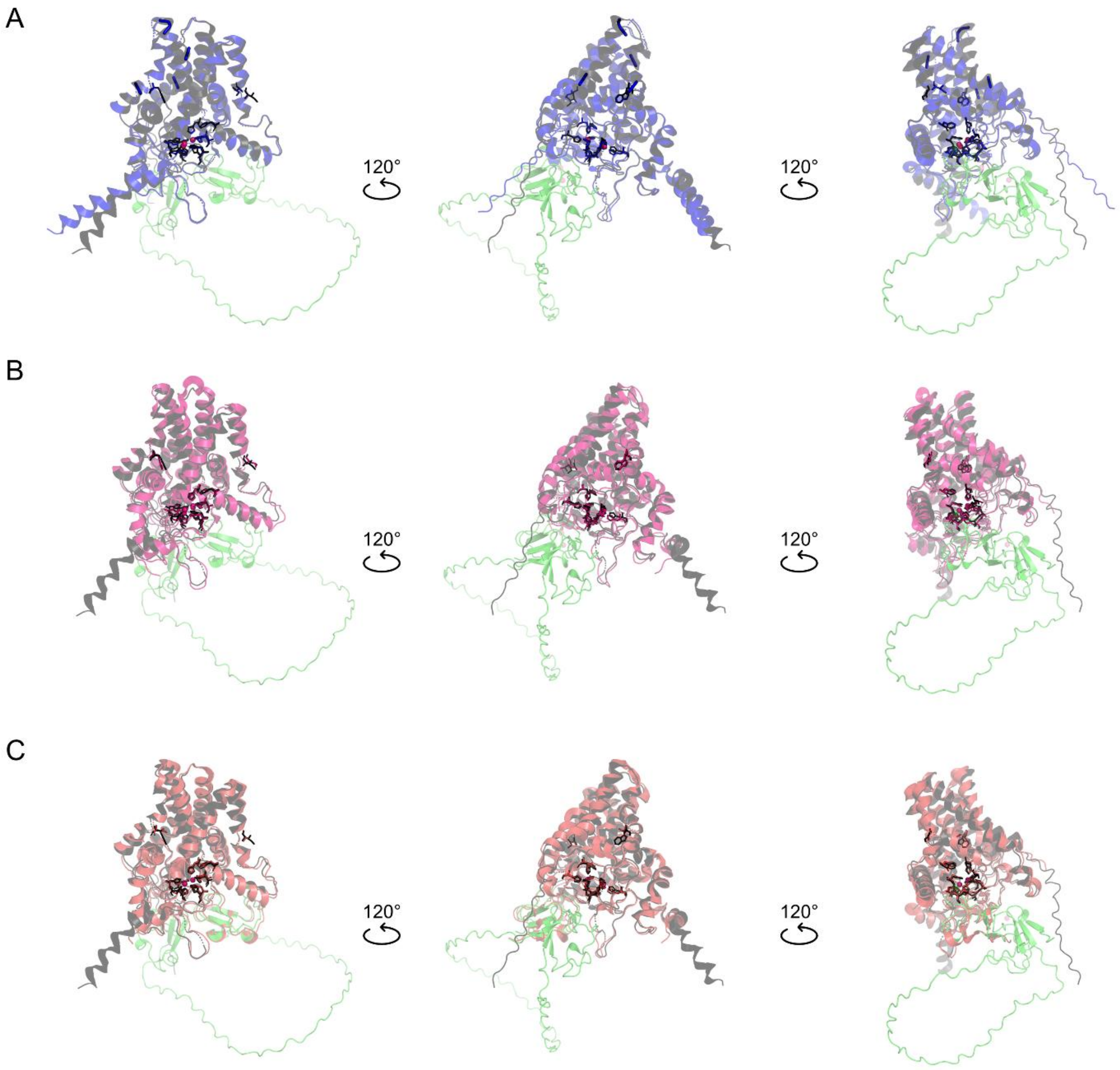

**Figure S2.** Structural comparison of M_AlkB (grey) with A) PpAlkB (blue), B) FtAlkB (PDB 8SBB, pink (Guo et al., 2023)) and C) FtAlkBG (PDB 8F6T, red (Chai et al., 2023)). Structures of M_AlkB, PpAlkB, PpAlkG (UniProt P00272, light green) and the interactions of M_AlkB and PpAlkG were predicted using AF3 (Abramson et al., 2024). The conserved histidines in the active center coordinating the two Fe-atoms (pink spheres) and the three residues subjected to mutation in this work are represented as sticks.

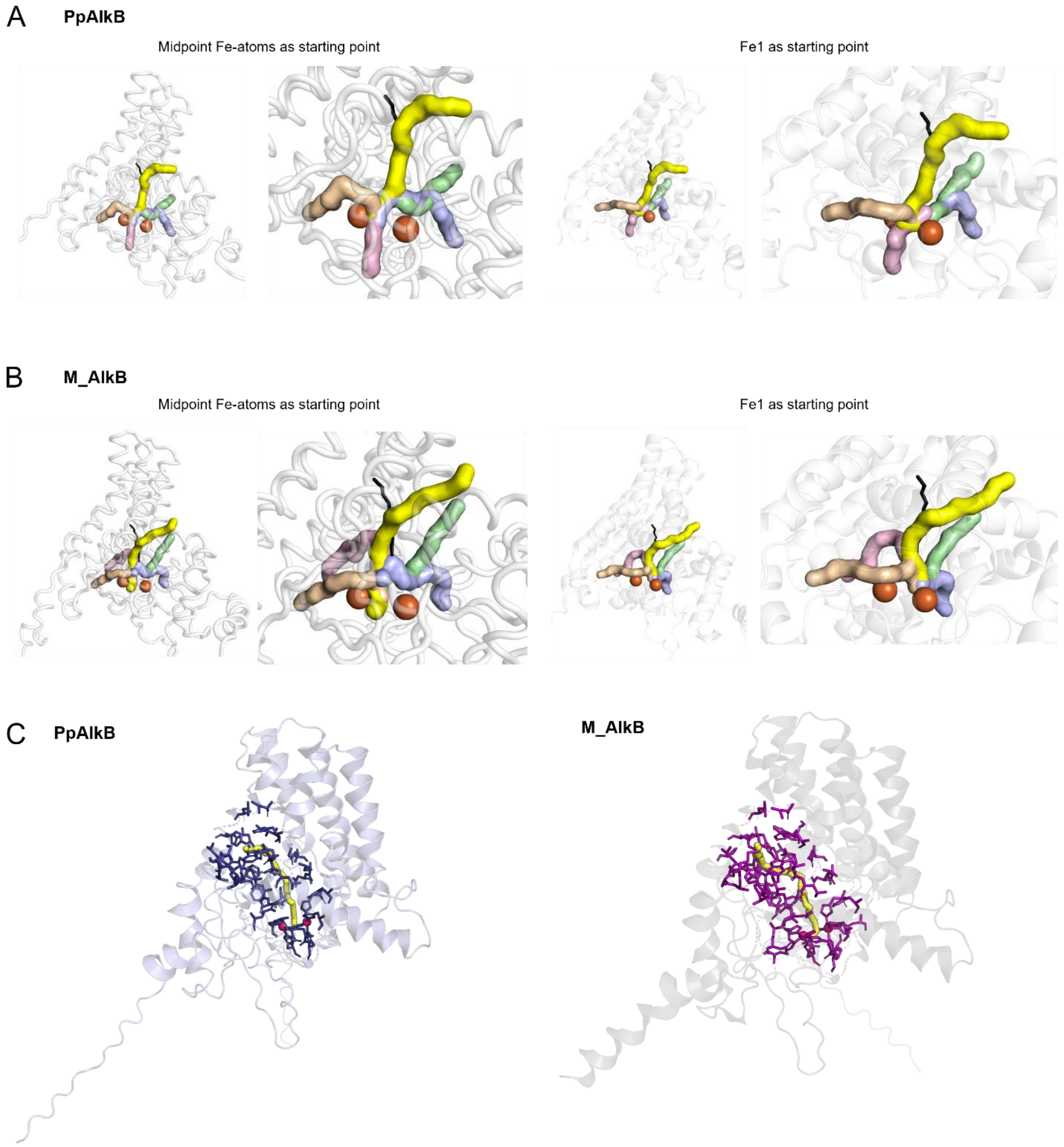

**Figure S3.** Top 5 tunnels identified by CAVER 2.0 (Marques et al., 2025; Stourac et al., 2019) with midpoint between the two Fe-atoms and Fe1 as a starting point in A) PpAlkB (tunnels 1-5) and B) M_AlkB (tunnels 1-5) with the selected tunnel 5 (yellow) used for further analysis and *n*-dodecane from PDB 82BB shown as a black stick. C) Selected substrate tunnels of PpAlkB (tunnel 5) and M_AlkB (tunnel 5) with residues lining the tunnel highlighted as sticks. D) Difference in residues between M_AlkB and PpAlkB located in four transmembrane helices around the predicted substrate tunnels.

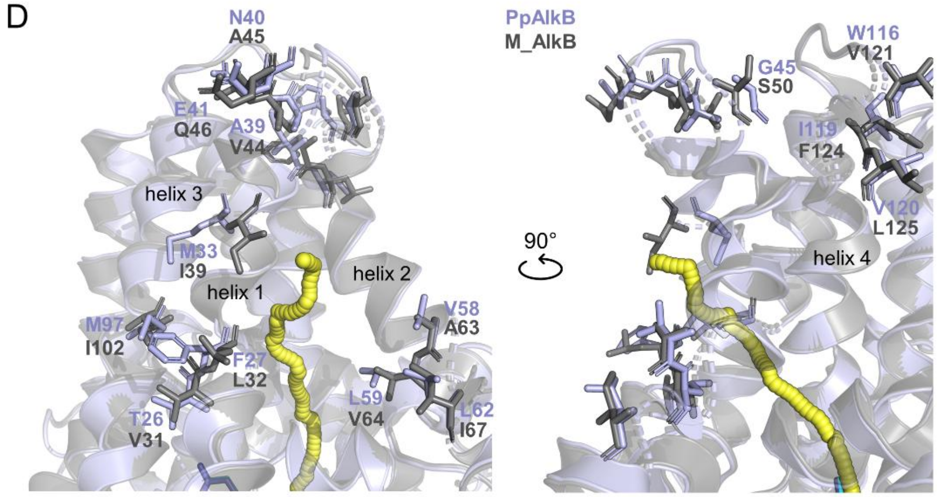

| PpAlkB | M_AlkB | helix |
|---|---|---|
| T26 | V31 | 1 |
| F27 | L32 | 1 |
| M33 | I39 | 1 |
| A39 | V44 | 1 |
| N40 | A45 | 1 |
| E41 | Q46 | 2 |
| G45 | S50 | 2 |
| V58 | A63 | 2 |
| L59 | V64 | 2 |
| L62 | I67 | 2 |
| M97 | I102 | 3 |
| W116 | V121 | 4 |
| I119 | F124 | 4 |
| V120 | L125 | 4 |

**Figure S3.** – continued.

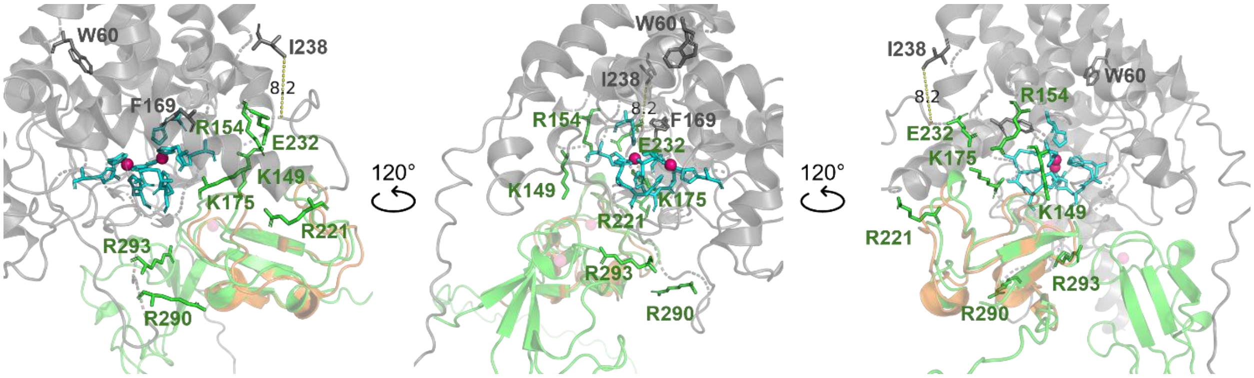

**Figure S4.** Interaction of M_AlkB and PpAlkG (light green) as predicted by AF3 (Abramson et al., 2024) and alignment with FtAlkG (PDB 8F6T, orange) (Chai et al., 2023). Selected M_AlkB residues located on cytosolic loops, which might be involved in the binding of AlkG via salt-bridge formation, are highlighted as green sticks. The residues are based on the AlkB-AlkG interactions in the structure of FtAlkBG (8F6T) (Groves et al., 2023; Mikulska-Ruminska et al., 2025). The residues R221 and E232 are located on a loop, which is in close proximity to residue I238 (8.2 Å).

### 2 Substrates and products

**Table S4.** List of products derived from conversions of linear fatty acid and fatty alcohol esters used in this work.

| Substrates and products based on linear fatty acids | | | | | |
|---|---|---|---|---|---|
| **C5 FAc** | | **C9FAc** | | **C12FAc** | |
| **1** | 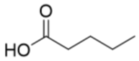 | **2** | 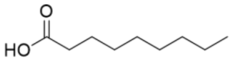 | **3** | 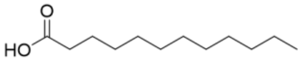 |
| **1a** | 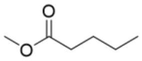 | **2a** | 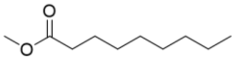 | **3a** | 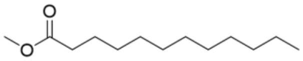 |
| **1b** | 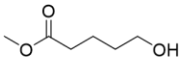 | **2b** | 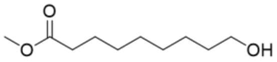 | **3b** | 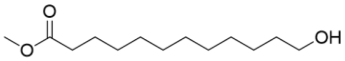 |
| **1c** | 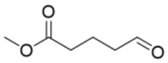 | **2c** | 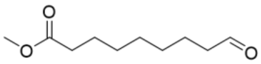 | **3c** | 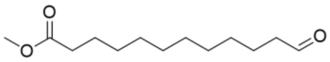 |
| **1d** | 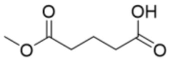 | **2d** | 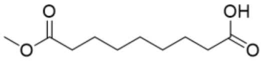 | **3d** | 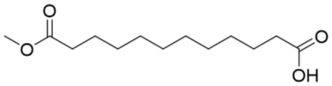 |
| **1i** | 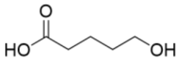 | **2i** | 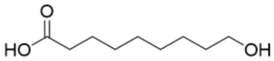 | **3i** | 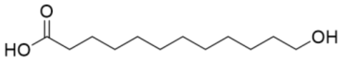 |

| Substrates and products based on linear fatty alcohols | | | | | |
|---|---|---|---|---|---|
| **C5 FAl** | | **C9FAl** | | **C12FAl** | |
| **4** | 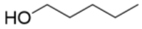 | **5** | 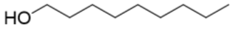 | **6** | 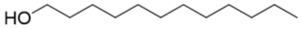 |
| **4a** | 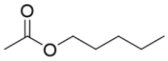 | **5a** | 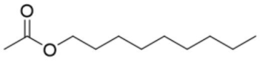 | **6a** | 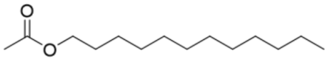 |
| **4b** | 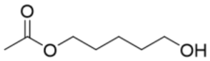 | **5b** | 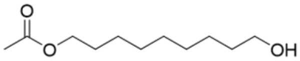 | **6b** | 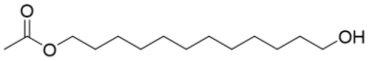 |
| **4c** | 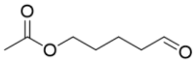 | **5c** | 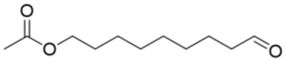 | **6c** | 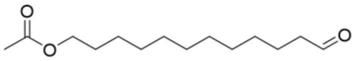 |
| **4d** | 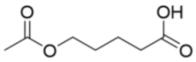 | **5d** | 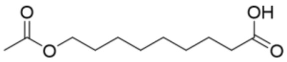 | **6d** | 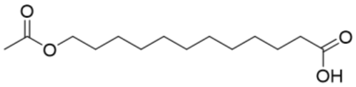 |
| **4e** | 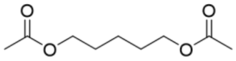 | **5e** | 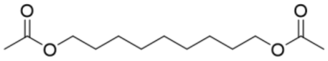 | **6e** | 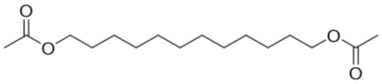 |
| **4g** | 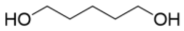 | **5g** | 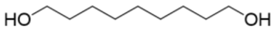 | **6g** | 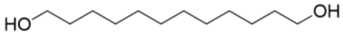 |

**Table S5.** List of products derived from conversions of branched fatty acid and fatty alcohol esters, and *n*-alkanes used in this work.

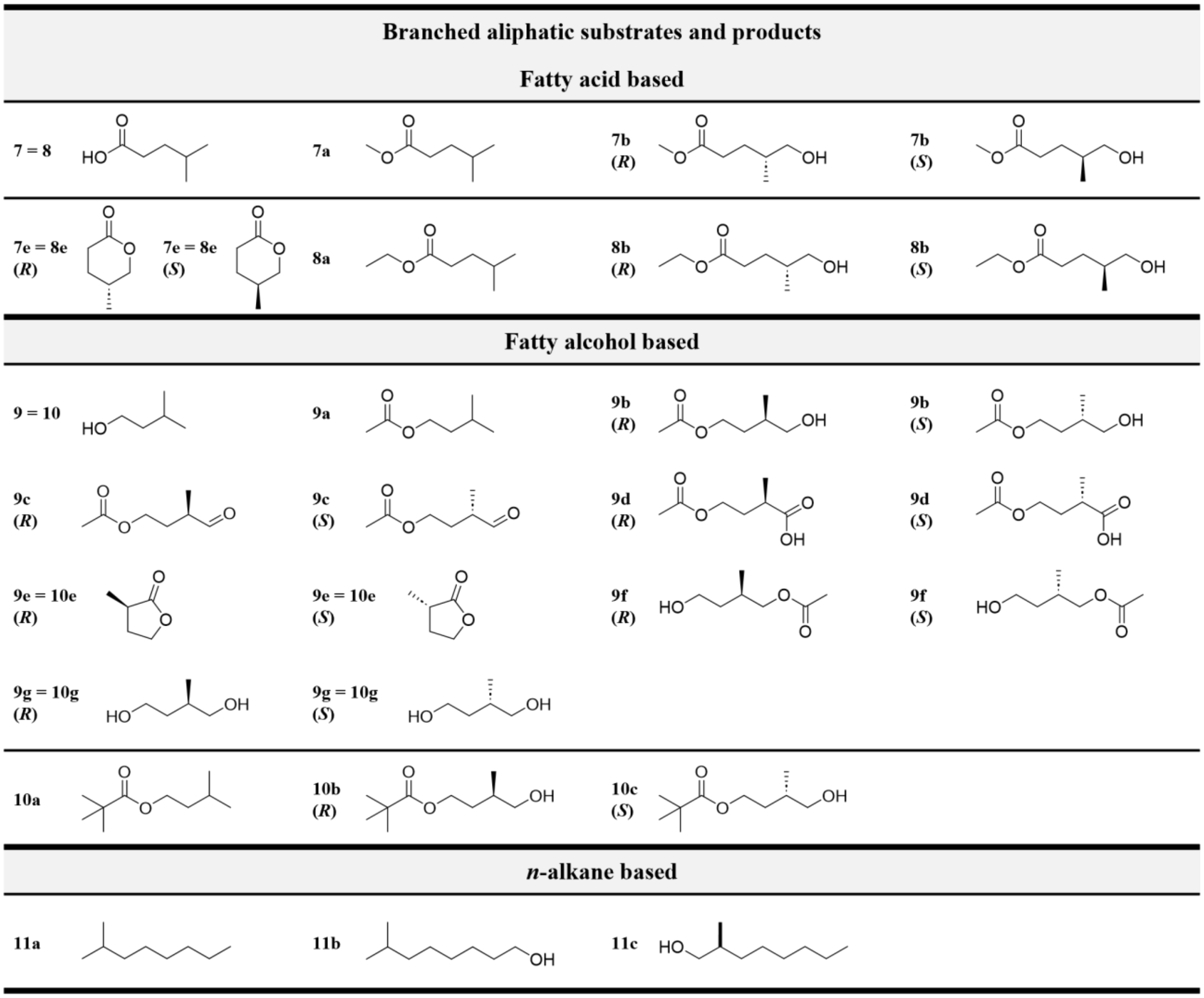

SwissADME was used to predict logP values of the investigated substrates (Daina, Michielin, & Zoete, 2017). SMILES inputs for each compound generated five logP models (iLOGP, XLOGP3, WLOGP, MLOGP, and SILICOS-IT). Among these, XLOGP3, known for its reliability in common organic structures and was selected for the comparison (Table S6).

**Table S6.** Predicted logP values of substrates tested in this study using SwissADME (Daina et al., 2017).

| Substrate | Canonical SMILES | XLOGP3 | Substrate | Canonical SMILES | XLOGP3 |
|---|---|---|---|---|---|
| **1a** | CCCCC(=O)OC | 1.96 | **7a** | COC(=O)CCC(C)C | 2.08 |
| **2a** | CCCCCCCCCC(=O)OC | 4.32 | **8a** | CCOC(=O)CCC(C)C | 2.44 |
| **3a** | CCCCCCCCCCCCC(=O)OC | 5.41 | **9a** | CC(CCOC(=O)C)C | 2.25 |
| **4a** | CCCCCOC(=O)C | 2.30 | **10** | CCCCCCC(C)C | 4.80 |
| **5a** | CCCCCCCCCCOC(=O)C | 3.95 | | | |
| **6a** | CCCCCCCCCCCCCOC(=O)C | 5.58 | | | |

### 3 Strains, plasmids and primers

Table S7 lists all strains and plasmids used in this work. Primers for site-directed mutagenesis are provided in Table S8. An example plasmid map of the construct used for the expression of M_AlkB variants is shown in Figure S5.

**Table S7.** All strains used in this work. All plasmids for the expression of M_AlkB(wt/mutant) were based on the broad-host vector pCom10 (Smits, Seeger, Witholt, & van Beilen, 2001) using the alkane-inducible promoter $P_{alkB}$ system. The sequences encoding *alkFGTL* are derived from *Ectopseudomonas oleovorans* (PpAlkFGTL) as described in previous work (Nigl et al., 2025).

| Strain | Enzyme variant encoded | Ref. |
|---|---|---|
| *E. coli* BL21 (DE3) pPpAlkB-FGTL | PpAlkB | (Julsing et al., 2012; Nigl et al., 2025; van Nuland, Eggink, & Weusthuis, 2016) |
| *E. coli* BL21 (DE3) pM_AlkB-FGTL | M_AlkB | (Nigl et al., 2025) |
| *E. coli* BL21 (DE3) pM_AlkB(I238V)-FGTL | M_AlkB I238V | (Nigl et al., 2025) |
| *E. coli* BL21 (DE3) pM_AlkB(W60)-FGTL | M_AlkB W60S | This study |
| *E. coli* BL21 (DE3) pM_AlkB(F169L)-FGTL | M_AlkB F169L | (Nigl et al., 2025) |
| *E. coli* BL21 (DE3) pM_AlkB(H278A)-FGTL | M_AlkB H278A | This study |
| *E. coli* BL21 (DE3) pM_AlkB(W60S_I238V)-FGTL | M_AlkB W60S I238V | This study |

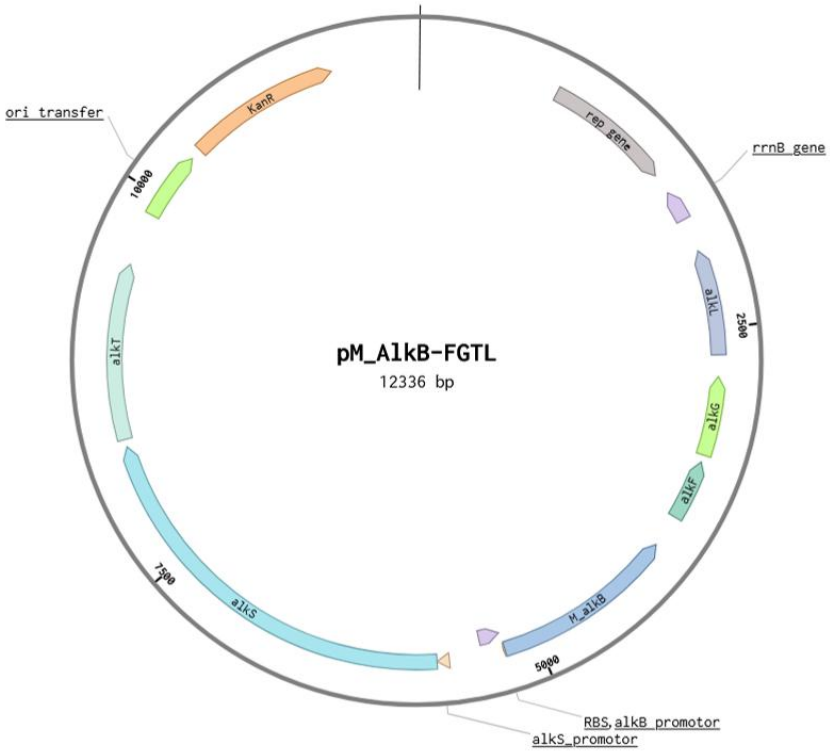

**Figure S5.** An example plasmid map of pM_AlkB-FGTL. For pPpAlkB-FGTL M_AlkB was replaced by the coding sequence of PpAlkB.

**Table S8.** Forward (FP) and reverse (RP) primers used to generate single and double mutants used in this work.

| Mutation | Primers |
|---|---|
| **W60S** | FP: ATCCTTAGCTACGGCGCAGTTCCGC<br>RP: GCCGTAGCTAAGGATCAACACCAAGCC |
| **F169L** | FP: GATGAACAGATGACCATAGCCTACC<br>RP: GGTCATCTGTTCATCGAACACAATAAG |
| **I238V** | FP: CGCGGTTAAAACCATCGGCTGAAGAAC<br>RP: CCGATGGTTTTAACCGCGGTGCTTT |
| **H278A** | FP: TCCGTAGGCCTCAATGTAGTTGGCGCTG<br>RP: ATTGAGGCCTACGGACTACTGCGCGAGAAGATG |

### 4 Analytical methods

#### 4.1 Gas chromatography

GC-MS was used for qualitative analysis and compound identification. The measurements were performed on a Shimadzu GC-MS-QP2010 SE equipped with an AOC-20i/s autosampler and injector unit. The method parameters are summarized in Table S9. The concentrations of the analytes in the biotransformation samples were quantified via GC-FID (Shimadzu Nexis GC-2030 and AOC-20i/s autosampler and injector unit) using the methods presented in Table S10. To determine the *ee* of chiral products and quantify product formation, samples were analyzed via GC-FID (Shimadzu Nexis GC-2030 and AOC-20i/s autosampler and injector unit) equipped with chiral columns as described in

Table **S11** and Table S12.

**Table S9.** GC-MS parameters of method used to analyze substrates and products formed in reactions with **1a** – **11a**.

| AC-M1 (1a-11a) | | |
|---|---|---|
| **Column** | **Name** | **ZB-5MS/ZB-5*PLUS*** |
| | Phase | 5% Phenyl-Arylene / 95% Dimethylpolysiloxane |
| | Dimensions | Length: 30.0 m<br>Inner diameter: 0.25 mm<br>Film thickness: 0.25 µm |
| **Oven temperature** | Profile | Hold: 50°, 3 min<br>Heating: to 300°C, 30°C/min<br>Hold: 300°C, 3 min |
| **Injection port** | Volume | 1 µL |
| | Carrier Gas | He |
| | Temperature | 250°C |
| | Split Ratio | 9.1 |
| **MS Detector** | Temperature | Ion source: 250°C<br>Interface: 320°C |
| | Total flow | 15.0 mL/min |

| | | |
|---|---|---|
| | Column flow | 1.19 mL/min |
| | Pressure | 67.2 kPa |

**Table S10.** Parameters of the GC-FID (achiral column) methods used to quantify substrates and products from biotransformation samples.

| | | AC-M2 (2a, 4a, 5a, 6a) | AC-M3 (9a) | AC-M4 (11a) |
|---|---|---|---|---|
| **Column** | **Name** | **Zebron ZB-5** | | |
| | Phase | 5% Phenyl / 95% Dimethylpolysiloxane | | |
| | Dimensions | Length: 30 m<br>Inner diameter: 0.32 mm<br>Film Thickness: 0.25 µm | | |
| **Oven temperature** | Profile | Hold: 50 °C, 1 min<br>Heating:<br>to 150 °C, 40 °C/min<br>to 250 °C, 20 °C/min<br>Hold: 250 °C, 2 min | Hold: 50 °C, 2 min<br>Heating: to 250 °C, 15 °C/min<br>Hold: 250 °C, 2 min | Hold: 50 °C, 2 min<br>Heating: to 180 °C, 5 °C/min<br>to 250 °C, 40 °C/min<br>Hold: 250 °C, 2 min |
| **Injection port** | Volume | 1 µL | | |
| | Carrier Gas | $N_2$ | | |
| | Temperature | 250 °C | | |
| | Split Ratio | 10.0 | | |
| **FID Detector** | Temperature | 320 °C | | |
| | Air flow rate | 400 mL/min | | |
| | $H_2$ flow rate | 40 mL/min | | |
| | $N_2$ flow rate | 32 mL/min | | |

**Table S11.** Parameters of the GC-FID (chiral Hydrodex βTBDAc column) methods used to determine *ee*-values of analytes from biotransformation samples.

| | | C-M1 (7a, 8a) | C-M2 (9a, 10a) |
|---|---|---|---|
| **Chiral column** | Name | Hydrodex β-TBDAc column | |
| | Phase | Heptakis-(2,3-di-O-acetyl-6-O-t-butyldimethylsilyl)-β-cyclodextrin | |
| | Dimensions | Length: 50 m<br>Inner diameter: 0.25 mm<br>Film Thickness: 0.25 µm | |
| **Oven temperature** | Profile | Hold: 90 °C, 3 min<br>Heating:<br>to 220 °C, 2 °C/min,<br>Hold: 220 °C, 3 min | Hold: 120°C, 10 min<br>Heating:<br>to 137°C, 0.5°C/min,<br>to 220°C, 10°C/min<br>Hold: 220 °C, 2 min |
| **Injection port** | Volume | 1 µL | |
| | Carrier Gas | $N_2$ | |
| | Temperature | 230 °C | |
| | Split Ratio | 2.0 | |
| **FID Detector** | Temperature | 250 °C | |
| | Air flow rate | 200 mL/min | |
| | $H_2$ flow rate | 32 mL/min | |
| | $N_2$ flow rate | 24 mL/min | |

**Table S12.** Parameters of the GC-FID (chiral Hydrodex β-6TBDM column) methods used to determine *ee*-values of analytes from biotransformation samples.

| | | C-M3 (9b) |
|---|---|---|
| **Chiral column** | Name | Hydrodex β-6TBDM |
| | Phase | Heptakis-(2,3-di-O-methyl-6-O-t-butyldimethyl-silyl)-β-cyclodextrin |
| | Dimensions | Length: 25 m<br>Inner diameter: 0.25 mm<br>Film thickness: 0.25 μm |
| **Oven temperature** | Profile | Hold: 100 °C, 10 min<br>Heating:<br>to 115 °C, 0.5 °C/min,<br>to 220 °C, 20 °C/min<br>Hold: 220 °C, 2 min |
| **Injection port** | Volume | 1 μL |
| | Carrier Gas | $N_2$ |
| | Temperature | 230 °C |
| | Split Ratio | 10.0 |
| **FID Detector** | Temperature | 250 °C |
| | Air flow rate | 200 mL/min |
| | $H_2$ flow rate | 32 mL/min |

**Table S13.** Retention times (Rt) of substrates and products on GC-MS using method AC-M1.

| GC-MS (ZB-5MS/ZB-5PLUS) | | | | | | | | |
|---|---|---|---|---|---|---|---|---|
| **Compound** | **Method** | **Rt (min)** | **Compound** | **Method** | **Rt (min)** | **Compound** | **Method** | **Rt (min)** |
| **1** | AC-M1 | 4.8 | **4a** | AC-M1 | 5.0 | **7e** | AC-M1 | 6.7 |
| **1a** | AC-M1 | 4.1 | **4b** | AC-M1 | 6.9 | **8a** | AC-M1 | 5.7 |
| **1b** | AC-M1 | 6.4 | **5** | AC-M1 | 6.8 | **8b** | AC-M1 | 7.3 |
| **1d** | AC-M1 | 6.8 | **5a** | AC-M1 | 7.6 | **9a** | AC-M1 | 4.9 |
| **2** | AC-M1 | 7.4 | **5b** | AC-M1 | 8.8 | **9b** | AC-M1 | 6.9 |
| **2a** | AC-M1 | 7.1 | **5g** | AC-M1 | 8.3 | **9c** | AC-M1 | 6.4 |
| **2b** | AC-M1 | 8.5 | **6** | AC-M1 | 8.4 | **9d** | AC-M1 | 7.5 |
| **2i** | AC-M1 | 8.7 | **6a** | AC-M1 | 8.9 | **9e** | AC-M1 | 5.6 |
| **3** | AC-M1 | 8.8 | **6b** | AC-M1 | 10.0 | **10a** | AC-M1 | 6.3 |
| **3a** | AC-M1 | 8.6 | **7** | AC-M1 | 5.7 | **11a** | AC-M1 | 4.8 |
| **3b** | AC-M1 | 9.8 | **7a** | AC-M1 | 5.1 | **11b** | AC-M1 | 6.8 |
| **3i** | AC-M1 | 9.9 | **7b** | AC-M1 | 6.9 | **11c** | AC-M1 | 6.7 |

**Table S14.** Retention times (Rt) of substrates and products on achiral GC-FID using methods AC-M2 and AC-M3.

| GC-FID achiral (Zebron ZB-5) | | | | | | | | |
|---|---|---|---|---|---|---|---|---|
| **Compound** | **Method** | **Rt (min)** | **Compound** | **Method** | **Rt (min)** | **Compound** | **Method** | **Rt (min)** |
| **2** | AC-M2 | 6.3 | **5a** | AC-M2 | 6.6 | **9b** | AC-M3 | 9.3 |
| **2a** | AC-M2 | 6.2 | **5b** | AC-M2 | 8.1 | **11a** | AC-M4 | 8.8 |
| **2b** | AC-M2 | 7.6 | **6a** | AC-M2 | 8.2 | **11b** | AC-M4 | 16.9 |
| **4a** | AC-M2 | 4.6 | **6b** | AC-M2 | 10.0 | **11c** | AC-M4 | 17.0 |
| **4b** | AC-M2 | 6.1 | **9a** | AC-M3 | 6.1 | | | |

**Table S15.** Retention times (Rt) of substrates and products based on chiral GC-FID using the methods C-M1 to M3.

| GC-FID chiral | | | | | | | | |
|---|---|---|---|---|---|---|---|---|
| **Compound** | **Method** | **Rt (min)** | **Compound** | **Method** | **Rt (min)** | **Compound** | **Method** | **Rt (min)** |
| **7a** | C-M1 | 7.6 | **8a** | C-M1 | 9.7 | **9g** | C-M2 | 30.9 |
| **7b** | C-M1 | 32.1 | **8b** | C-M1 | 32.2 | **(*S*)-9b** | C-M3 | 26.7 |
| **(*R*)-7e** | C-M1 | 49.5 | **9a** | C-M2 | 4.6 | **(*R*)-9b** | C-M2 | 27.5 |
| **(*S*)-7e** | C-M1 | 49.7 | **9b** | C-M2 | 25.5 | | | |

External calibration curves on GC-FID were created by measuring samples with known concentrations of commercially available compounds. The samples were prepared in resting cell buffer at concentrations ranging from 0 mM to 6 mM. They were extracted and prepared for GC-FID measurement in the same manner as the biotransformation samples. The curves were fitted using the linear regression model: y = ax + b (y: peak area; a: slope; x: analyte concentration; b: y-intercept was forced through 0). $R^2$ (coefficient of determination) is provided as a measure of variation. For compounds which were not commercially available, the concentrations were calculated based on calibration curves of structurally similar compounds.

**Table S16.** Calibration curve values for calculating analyte concentrations.

| **Calibration curves** | **Compound** | ***a*** | ***R²*** |
|---|---|---|---|
| **Achiral compounds** | **2a** | 256345 | 0.9985 |
| | **2b** | 260953 | 0.9997 |
| | **2** | 282056 | 0.9997 |
| | **4a** | 179469 | 0.9986 |
| | **5a** | 315542 | 0.9998 |
| | **6a** | 377382 | 0.9979 |
| **(Pro)chiral compounds** | **7a** | 750537 | 0.9958 |
| | **8a** | 848359 | 0.9922 |
| | **9a** | 591204 | 0.9999 |
| | **(*R*)-7e** | 309286 | 0.9842 |
| | **11b** | 233586 | 0.9850 |

##### 4.2 NMR-spectroscopy

NMR spectra were recorded on a Bruker AVANCE III 300 spectrometer ($^1$H: 300.36 MHz) with an autosampler. Chemical shifts δ are referenced to the residual proton signal of the deuterated solvent ($CDCl_3$: δ = 7.26 ppm ($^1$H). For the analysis, 5-10 mg of analyte was dissolved in $CDCl_3$. The $^1$H-NMR spectra were compared to the reported spectra.

### 5 Synthesis of product standards and product analysis

All commercially available chemicals and solvents were purchased from Merck, Roth, Sigma-Aldrich, TCI, and VWR, and used without further purification unless otherwise noted. Anhydrous dichloromethane was obtained by pre-drying ethanol-stabilized dichloromethane over $P_4O_{10}$, then heating it under reflux with $CaH_2$ for 24 hours under an argon atmosphere. It was distilled into an amber 1 L Schlenk bottle over activated 4 Å molecular sieves and under an argon atmosphere.

##### 5.1 Synthesis of isoamyl pivalate (10a)

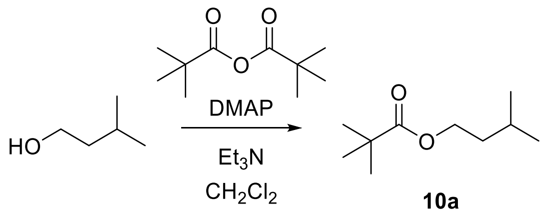

**10a**

To a 200 mL flame-dried Schlenk flask, anhydrous $CH_2Cl_2$ (35 mL) and isoamyl alcohol (2.5 mL, 22.7 mmol, 1 eq.) were added. The solution was cooled to 0 °C using a water-ice bath, and DMAP (0.14g, 2.3 mmol, 0.1 eq.) was added. Subsequently, $Et_3N$ (6.5 mL, 90.8 mmol, 4 eq.) and pivalic anhydride (2.8 mL, 27.2 mmol, 1.2 eq.) were slowly added. The yellow solution was left stirring at room temperature. After 24 h, the reaction was quenched with saturated $NaHCO_3$ (15 mL). After separating the organic and the aqueous phase, the aqueous phase was extracted with $CH_2Cl_2$ (20 mL, 3x). The combined organic layers were washed with brine (30 mL), dried over $Na_2SO_4$, filtered, and concentrated under reduced pressure. To remove the remaining DMAP, the extract was diluted with $CH_2Cl_2$, washed with HCl (1 M, 15 mL, 3x), and concentrated under reduced pressure. The crude product was further purified via flash column chromatography (cyclohexane:EtOAc, 20:1 (v/v)), affording the desired product as a colorless oil (1.35 g, 35%).

$C_{10}H_{20}O_2$ [172.27 g·mol$^{-1}$]; $R_f$ = 0.57 (cyclohexane:EtOAc = 20:1 (v/v), $KMnO_4$)

$^1$H NMR (300 MHz, $CDCl_3$): δ 4.08 (t, $J$ = 6.7 Hz, 2H), 1.79 – 1.60 (m, 1H), 1.51 (dd, $J$ = 13.5, 6.8 Hz, 2H), 1.19 (s, 9H), 0.92 (d, $J$ = 6.6 Hz, 6H).

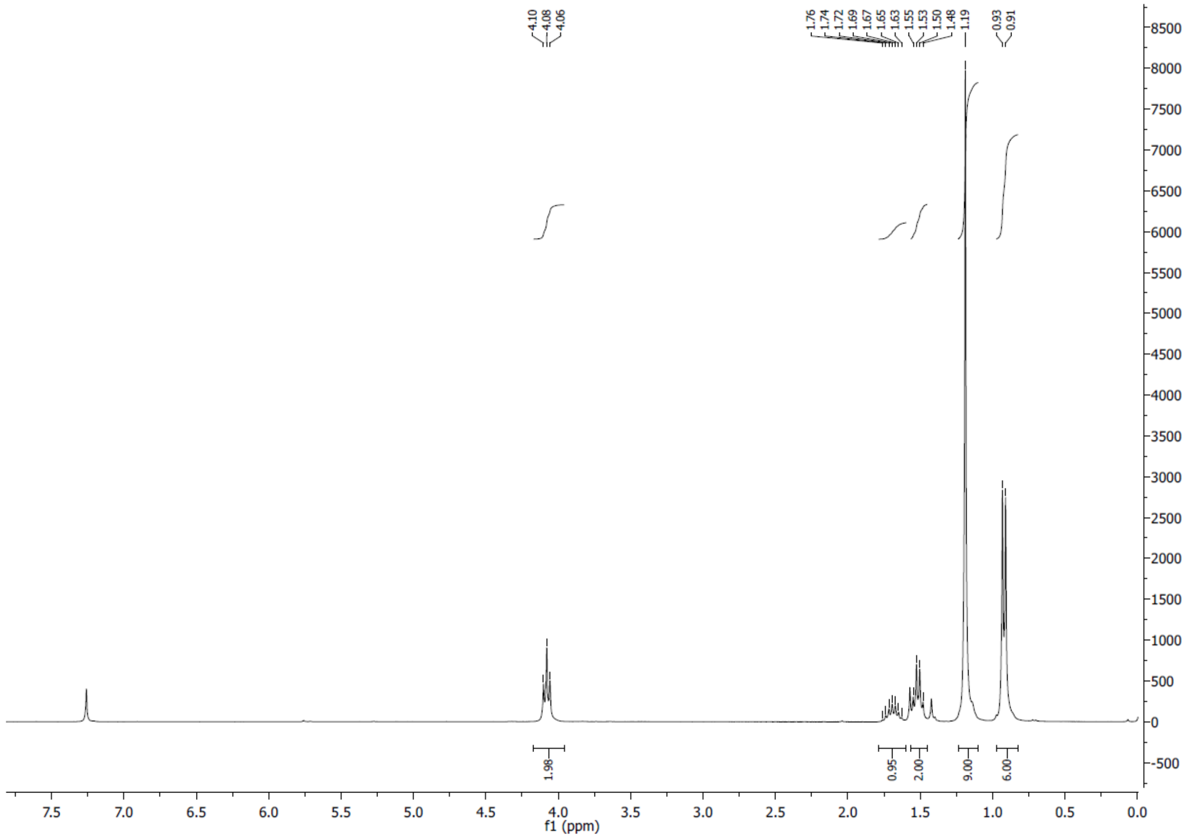

**Figure S6.** $^1$H-NMR of **10a**.

#### 5.2 Synthesis of standards for 4b, 5b, 6b, 9b

To obtain authentic standards of hydroxylated fatty alcohol acetates, standards were synthesized by acetylation of the corresponding diol. Therefore, 1,5-pentane diol **4g**, 1,9-nonane diol **5g**, 1,12-dodecane diol **6g**, and 2-methyl 1,4-butanediol **9g** were acetylated using *Pseudomonas cepacia* lipase (LPS, 38.6 U/mg, Sigma-Aldrich, Buchs, Switzerland) following previously published methods (Ferraboschi, Grisenti, Manzocchi, & Santaniello, 1994; Grisenti, Ferraboschi, Casati, & Santaniello, 1993). Briefly, to a solution of 0.48 mmol of diol in anhydrous methyl tert-butyl ether (MTBE, 1 mL) and LPS (312 U/mmol substrate), vinyl acetate (one eq. for linear diols and 3 eq. for branched diols) was added. The reaction mixture was stirred for 24 h in round-bottom flasks at 30 °C in an oil bath. After 24 h, the reaction mixture was filtered, centrifuged (16,000 x *g*, 4 °C, 15 min), and successful acetylation was confirmed by GC-MS (Figure S7).

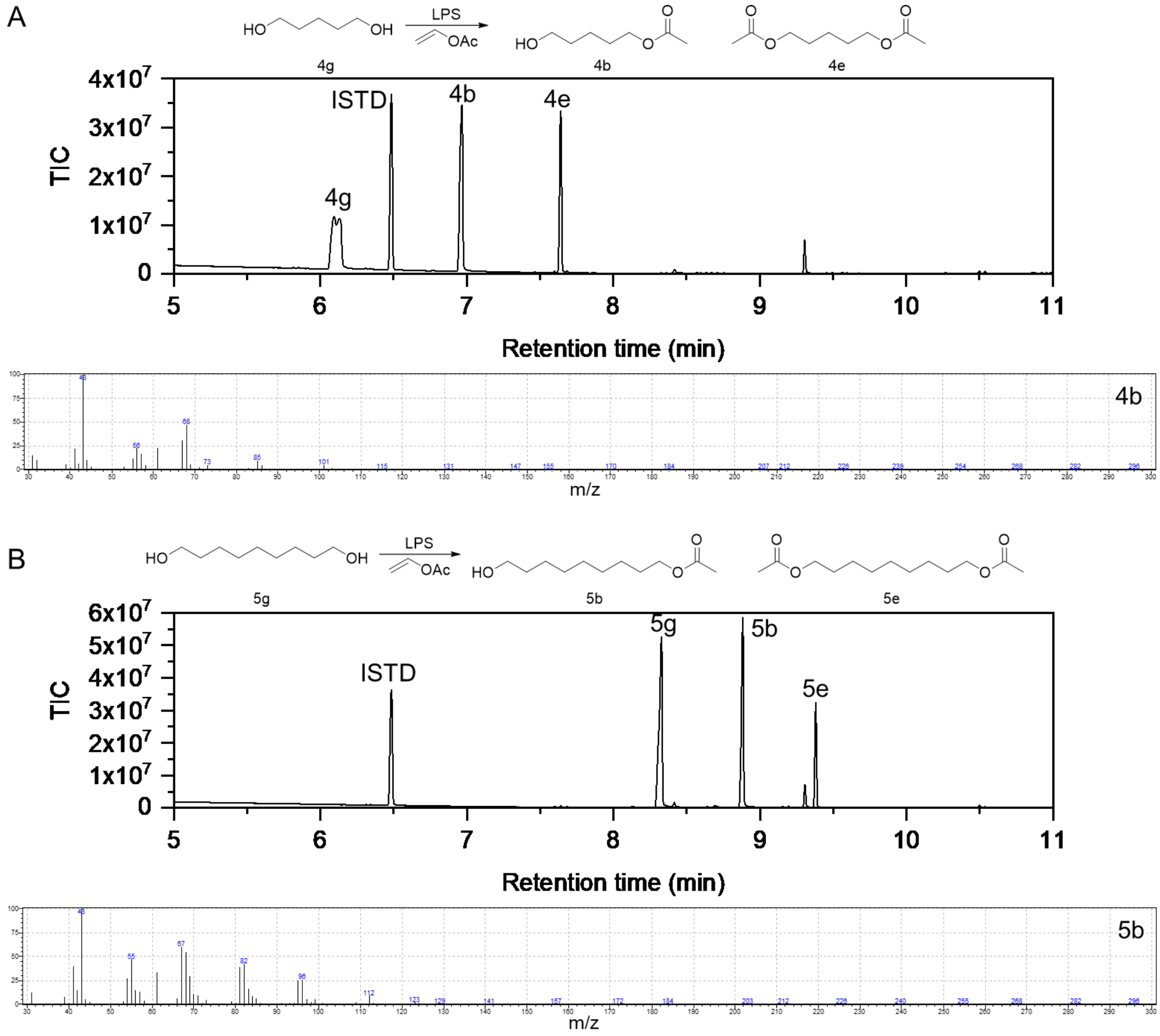

**Figure S7.** GC-MS results of *Pseudomonas cepacia* lipase (LPS)-catalyzed acetylation of A) 1,5-pentane diol **4g**, B) 1,9-nonane diol **5g**, C) 1,12-dodecane diol **6g**, and D) 2-methyl 1,4-butanediol **9g**.

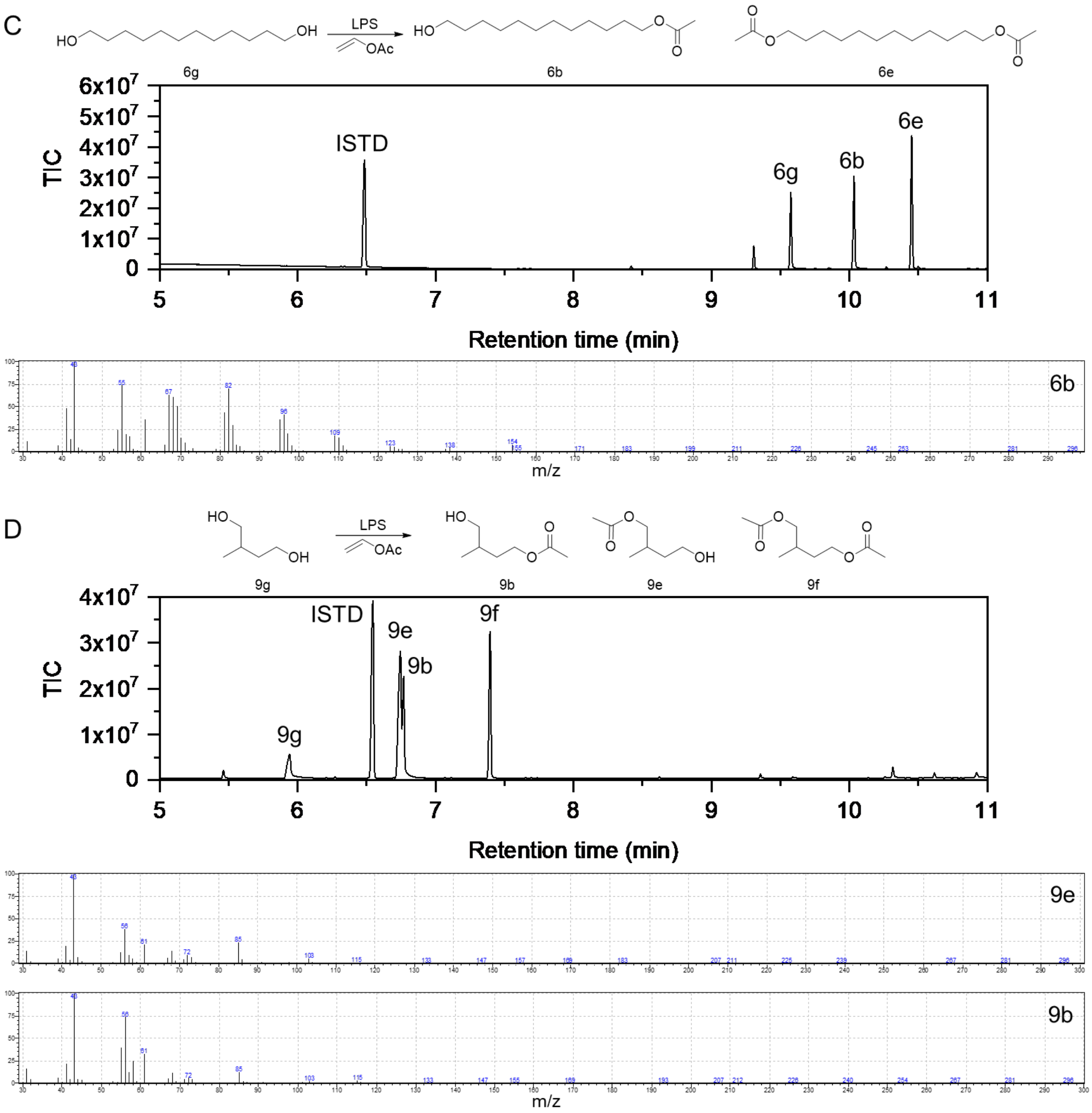

**Figure S5.** – continued.

### 6 Heterologous AlkB expression

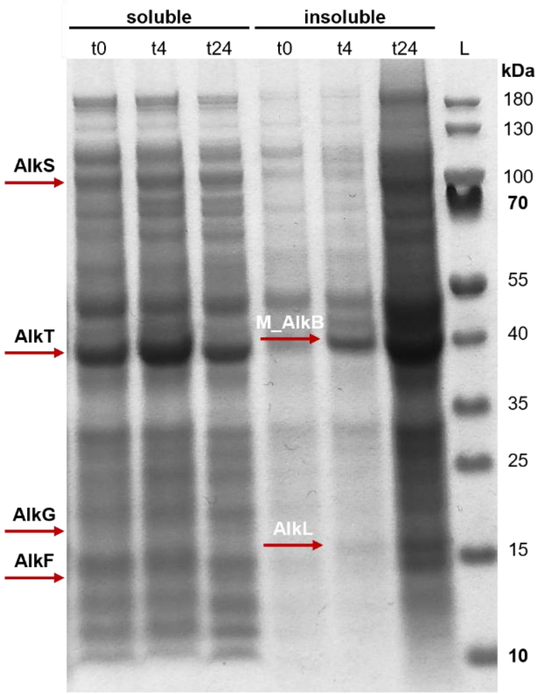

**Figure S8.** SDS-PAGE analysis of *M_alkBFGTL* expression. Samples were taken 0 h, 4 h, and 24 h after inducer addition (0.05% dicyclopropyl ketone). Soluble and insoluble fractions were obtained by cell lysis using BugBuster® Master Mix. The soluble proteins AlkS, AlkT, AlkG, and AlkF were expected at ~99 kDa, ~41 kDa, ~19 kDa, and ~15 kDa, respectively. The membrane-associated proteins M_AlkB (~ 40 kDa) and AlkL (~17 kDa) were expected in the insoluble fraction. The sizes were compared to the PageRuler™ Prestained protein Ladder (L) from Thermo Fisher Scientific.

### 7 GC analysis of AlkB reactions and controls

The peaks of the hydroxylated esters formed by AlkB were assigned by comparing the monoacetates formed by the LPS-catalyzed acetylation of the diols on GC-MS and GC-FID. Direct analysis via achiral GC-FID allowed rate quantification. Chiral GC-FID analysis was used to determine the % *ee* of the hydroxy products by comparing the absolute peak areas of the two enantiomers and to quantify the product formation rate for reactions using **7a** and **8a** as substrates. Hydrolysis of the hydroxylated fatty alcohol acetate to the corresponding diol allowed for comparison with an authentic standard of the diol and determination of *ee*-values and enantiomeric configuration. The *ee*-values of the hydroxylated branched fatty acid esters were determined by direct GC-FID measurements, and comparison of the corresponding lactone with authentic standards allowed the determination of the absolute configuration.

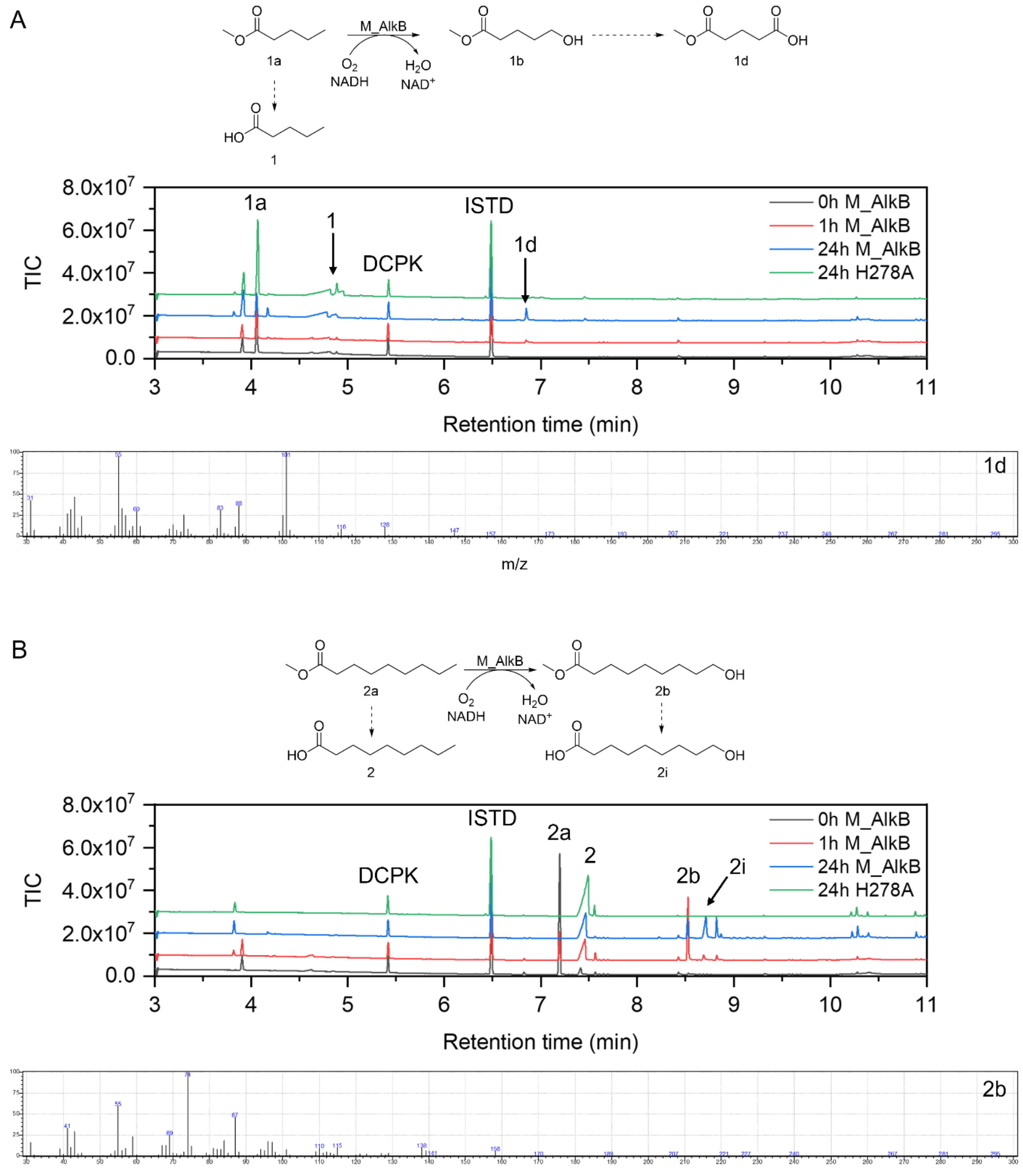

**Figure S9.** GC-MS analysis of the initial screening of substrate acceptance of M_AlkB toward **1a** - **6a** (A-F). The biotransformations were performed using 3.1 $g_{CDW}$/L for 24 h at 25 °C, 180 rpm and compared with a strain expressing an inactive AlkB variant (H278A) as negative control.

C

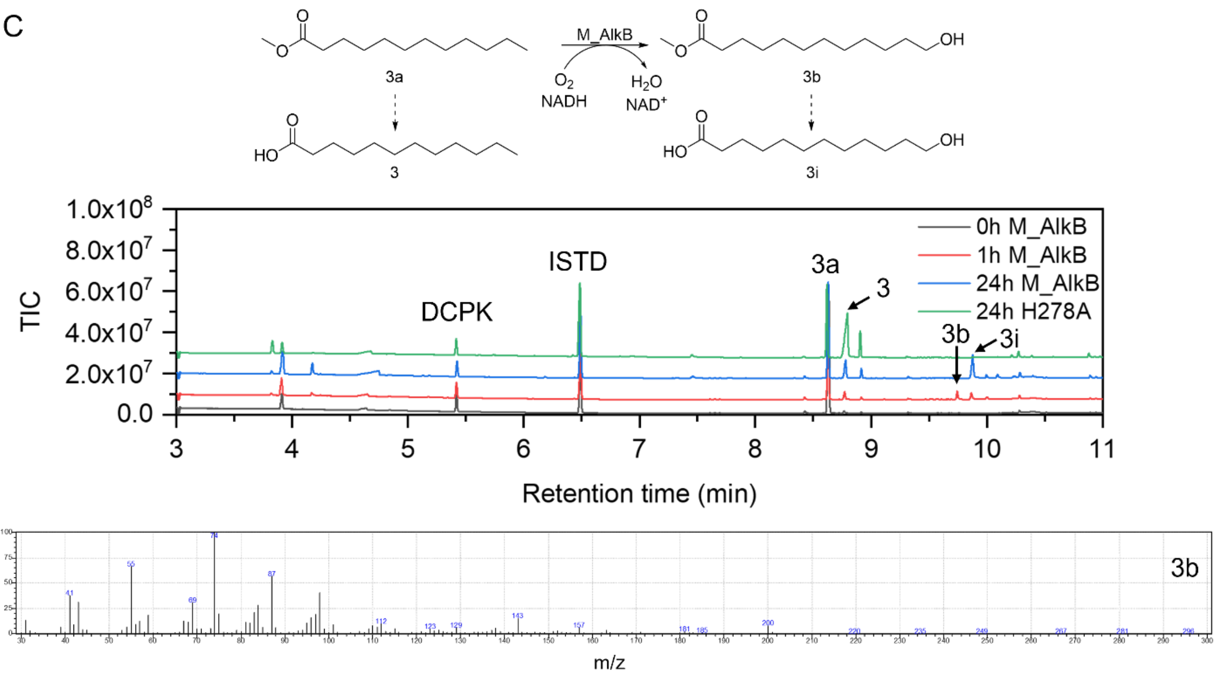

D

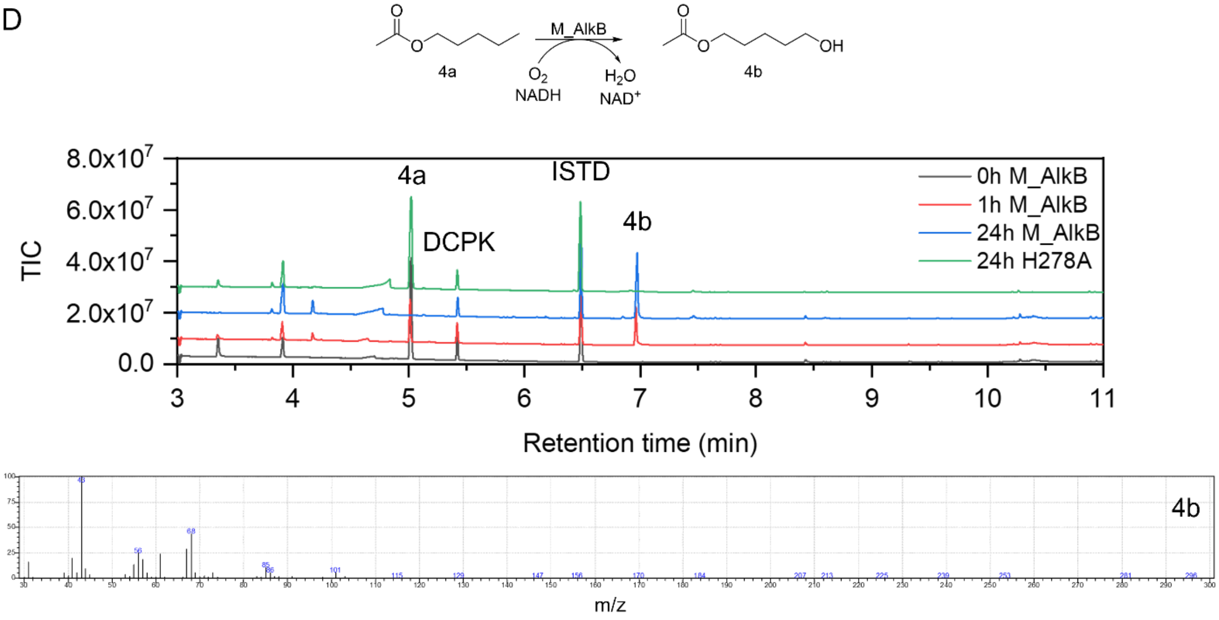

**Figure S9.** – continued.

E

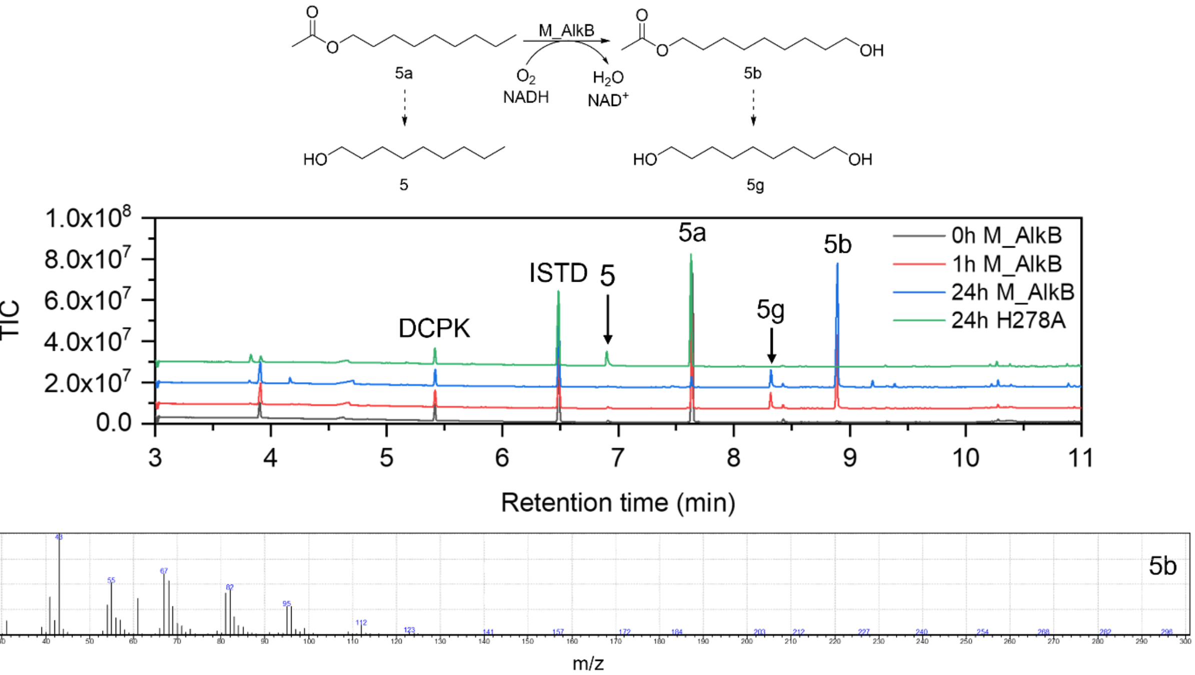

F

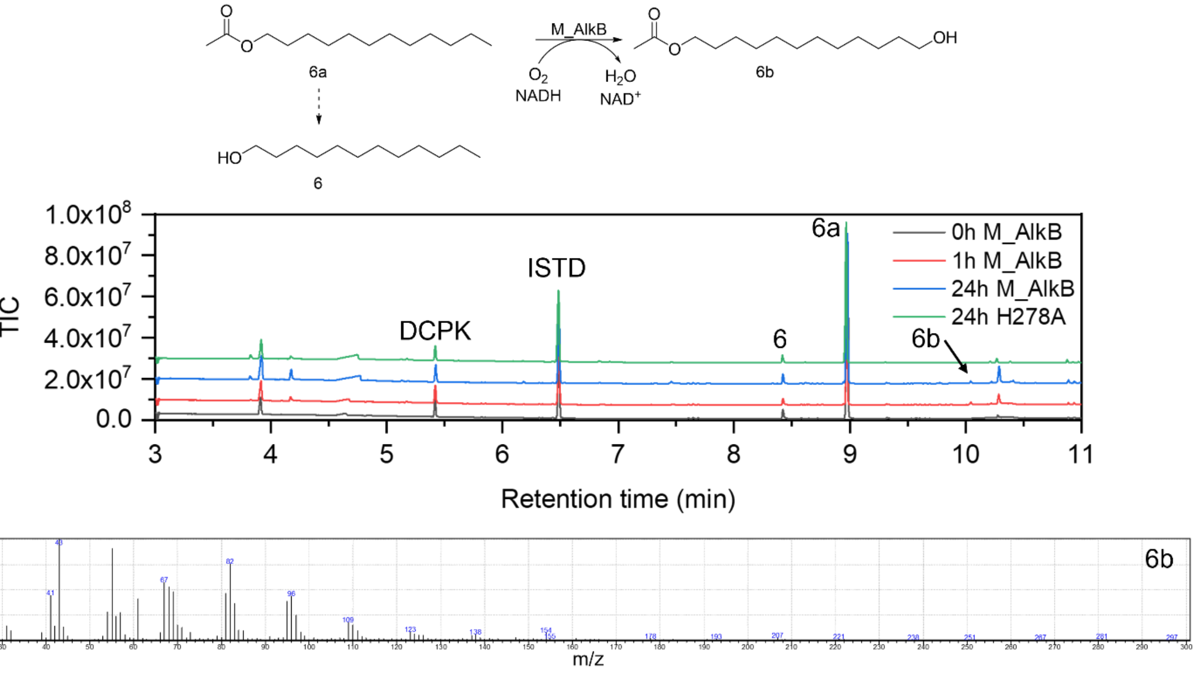

**Figure S9.** – continued.

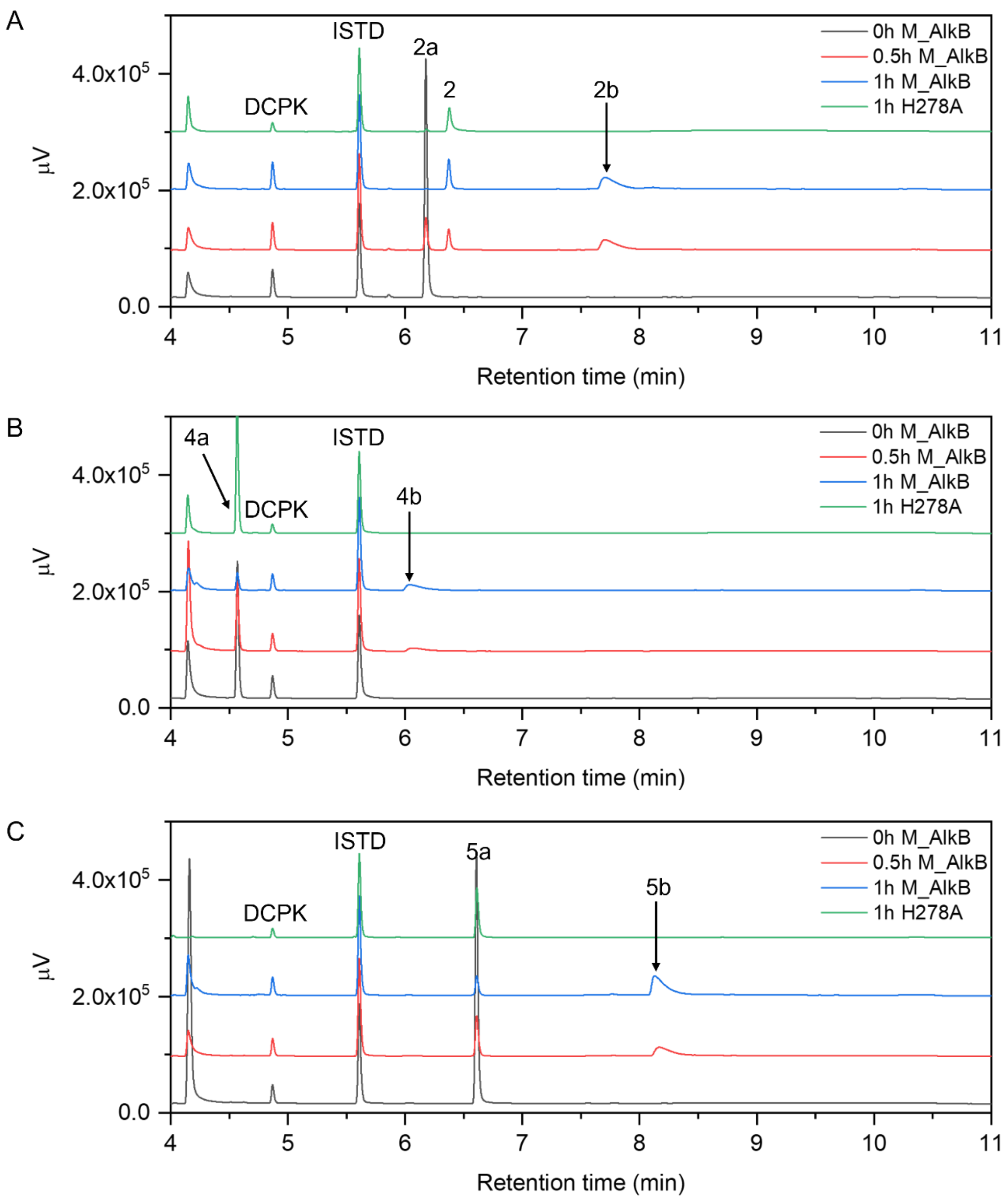

**Figure S10.** Exemplary GC-FID chromatograms of reactions with the linear esters A) **2a**, B) **4a**, C) **5a**, D) **6a** as substrates. The transformations were performed using 3.1 $g_{CDW}$/L for 24 h at 25 °C, 180 rpm, and compared with a strain expressing an inactive M_AlkB variant H278A as a negative control.

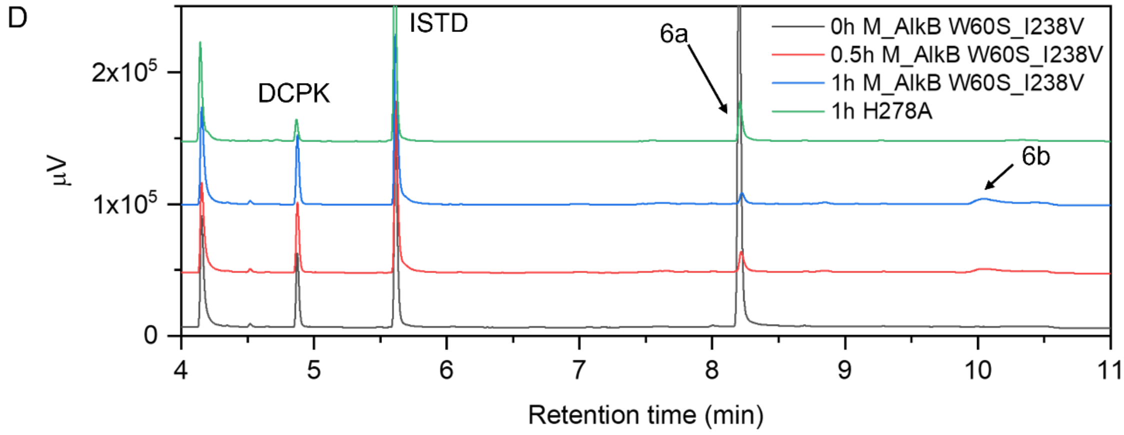

**Figure S10.** – continued.

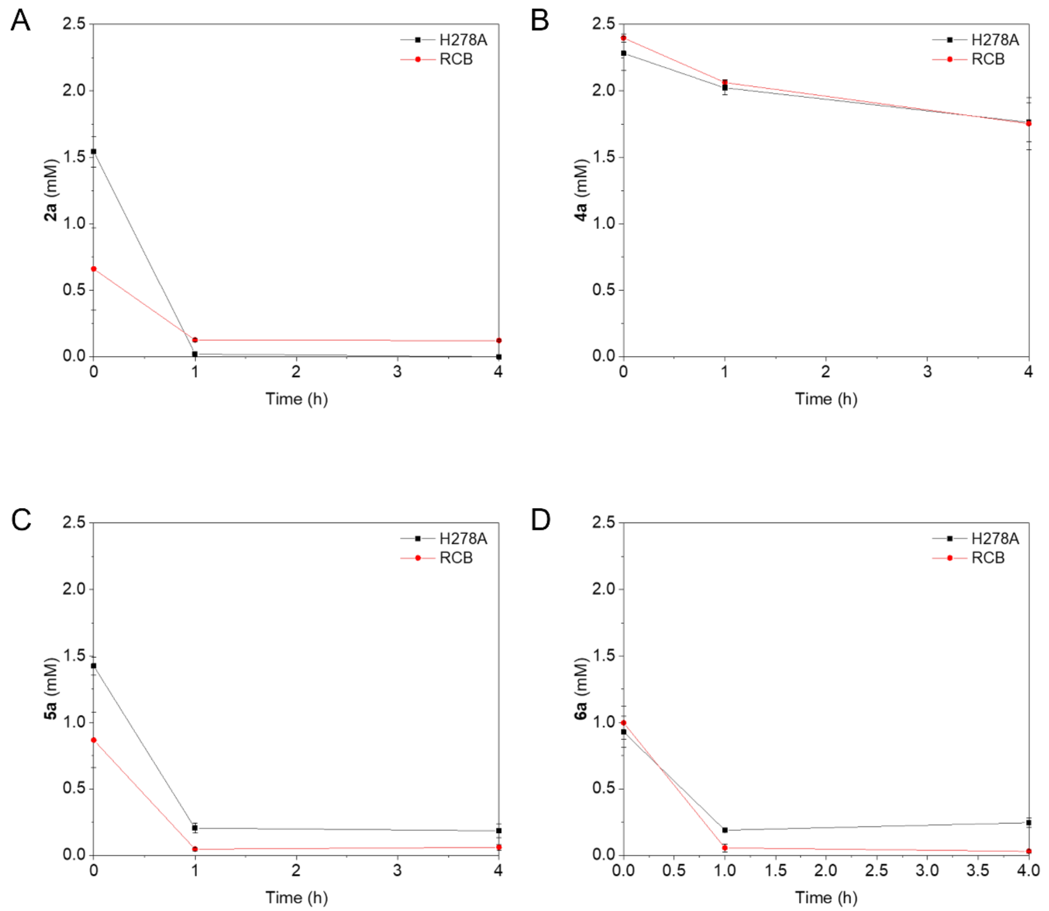

**Figure S11.** Substrate A) **2a**, B) **4a**, C) **5a,** and D) **6a** depletion in negative controls. The substrates were incubated with resting cells expressing the inactive variant M_AlkB H278A or resting cell buffer (RCB) only. Data represented as mean ± SD ($n$ = 3).

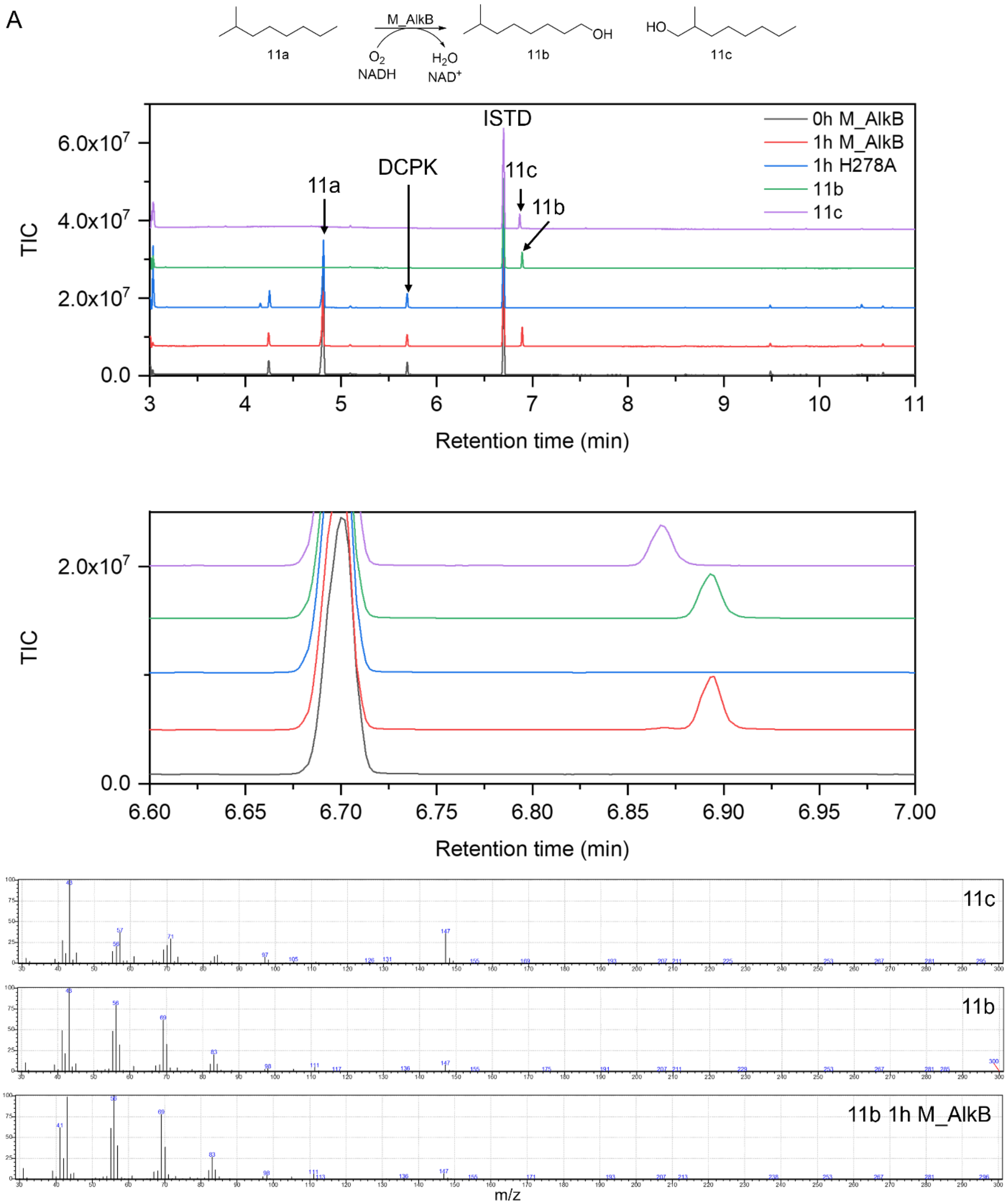

**Figure S12.** GC-MS analysis of M_AlkB-catalyzed reactions with **11a** and 3.1 $g_{CDW}$/L for 1 h at 25 °C, 180 rpm, and compared with a strain expressing an inactive M_AlkB variant (H278A) as negative control.

**Table S17.** Obtained activities of M_AlkB using **11a** as a substrate (mean ± SD; *n* = 3 biological replicates).

| **Product** | **Activity U/$g_{CDW}$** | **Activity ratio 11b/11c** |
|---|---|---|
| **11b** | 0.42 ± 0.01 | - |
| **11c** | 0.012 ± 0.001 | 36.1 |

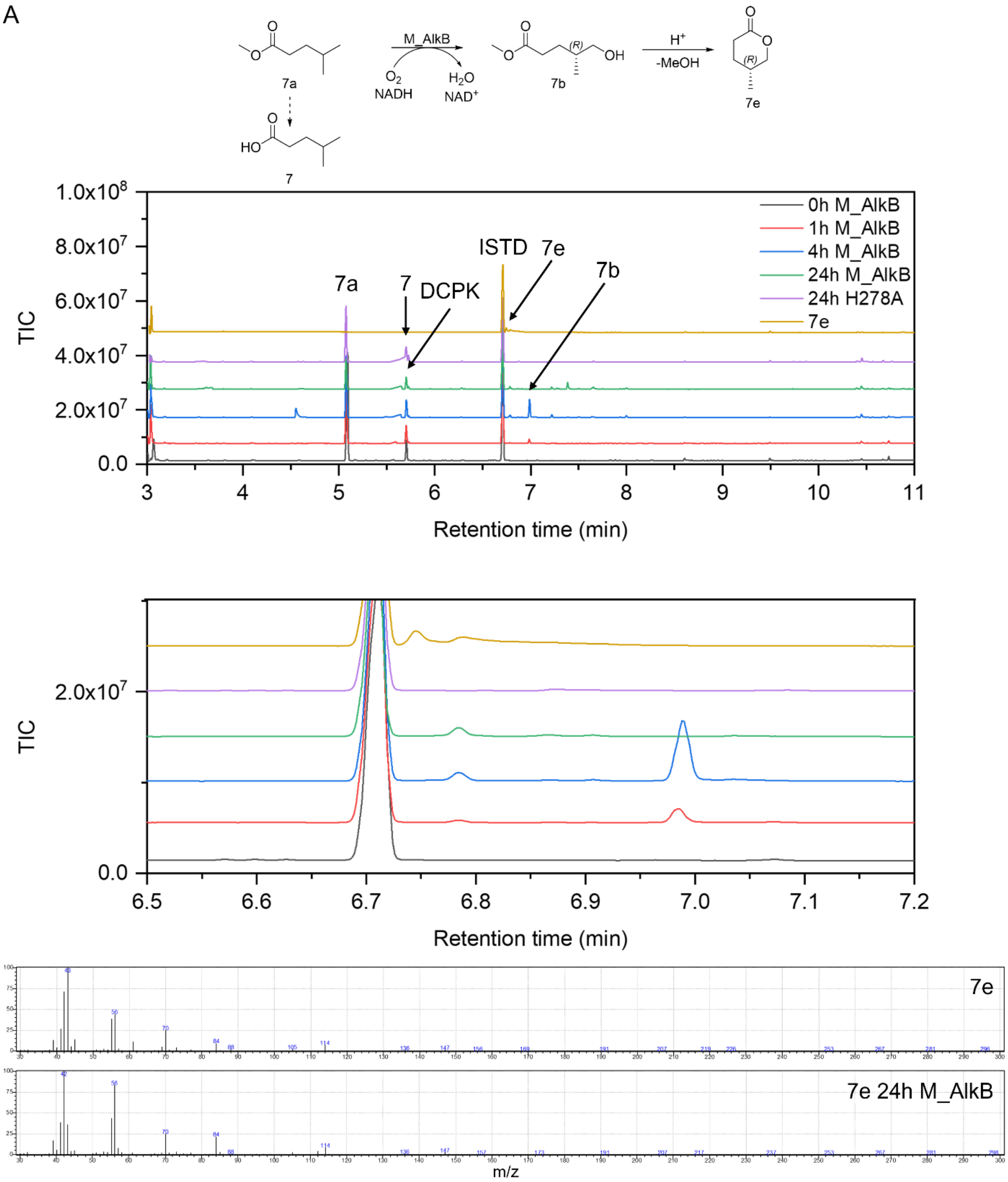

**Figure S13.** GC-MS analysis of initial activity screening of M_AlkB using A) **7a** and B) **8a**. The reactions were performed using 3.1 $g_{CDW}$/L at 25 °C, 180 rpm for 24 h, and compared to a strain expressing an inactive M_AlkB variant (H278A) as a negative control.

B

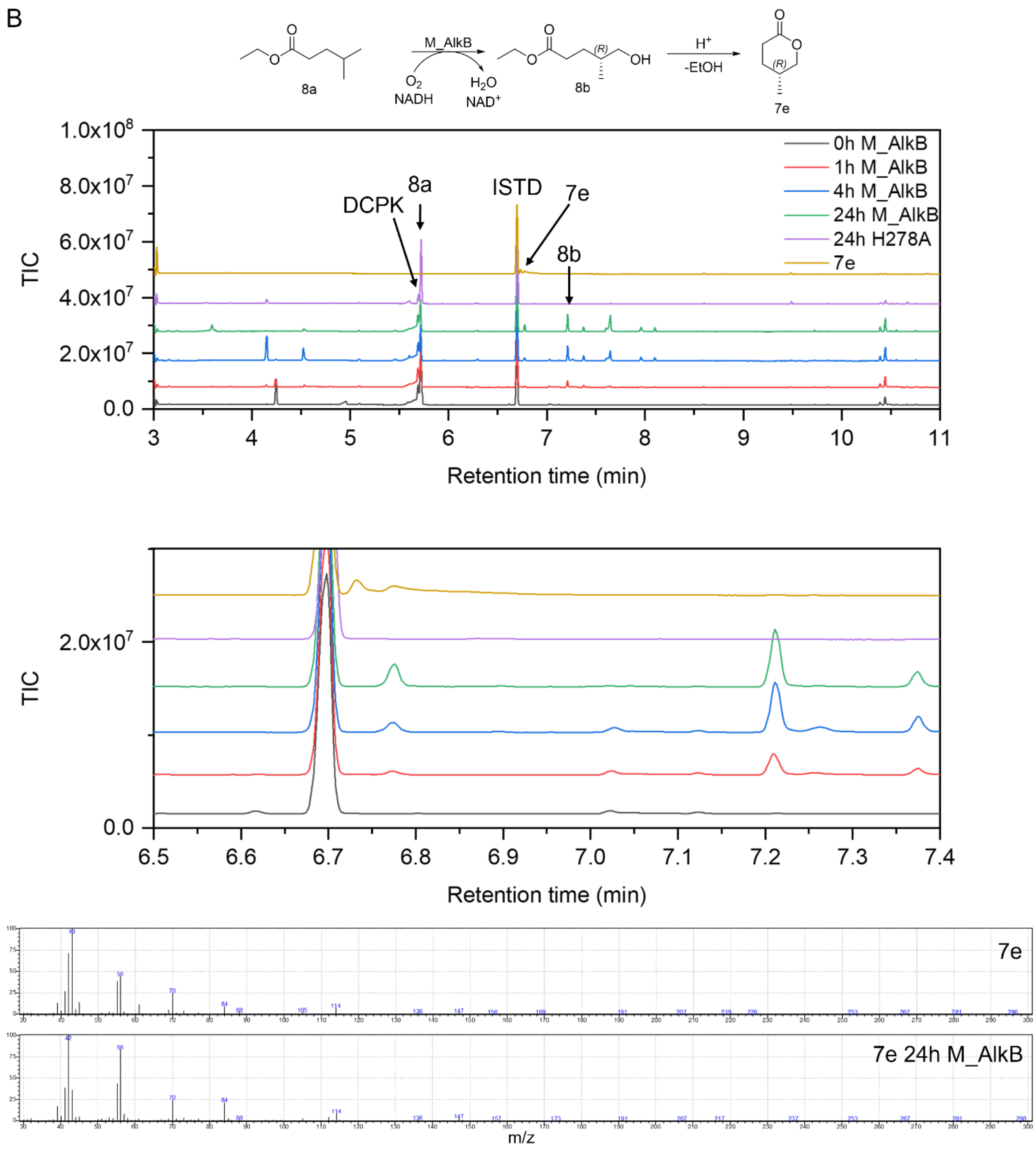

**Figure S13.** – continued.

A

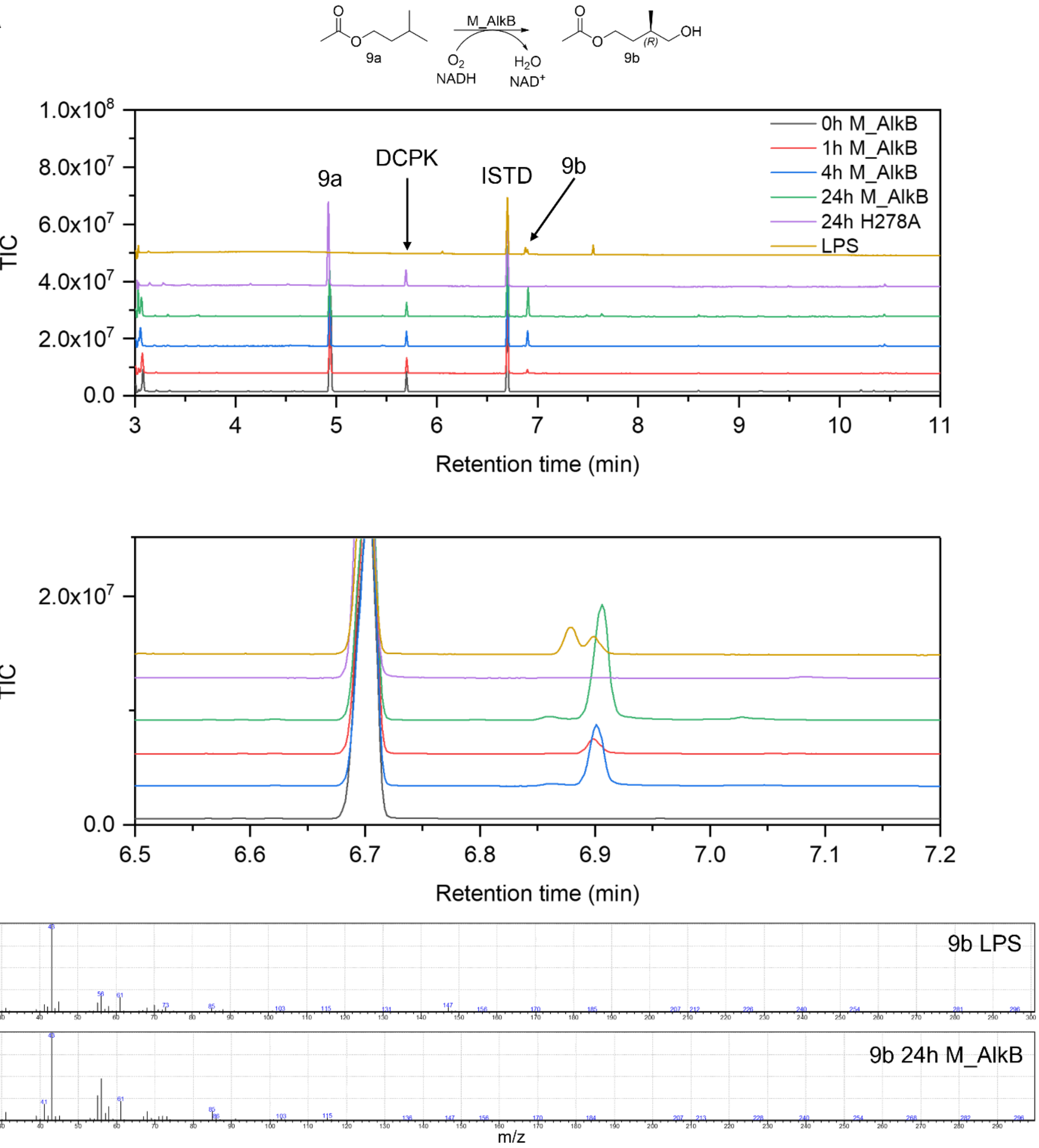

**Figure S14.** GC-MS analysis of M_AlkB-catalyzed reactions with **9a** using A) ethanol as co-solvent, which hindered the overoxidation of **9a**, while the use of B) DMSO as solvent shows overoxidation of **9b** to **9c** and **9d**. Upon acidification, **9d** was lactonized (L) to **9e**. The reactions were compared with a strain expressing an inactive M_AlkB variant (H278A) as a negative control.

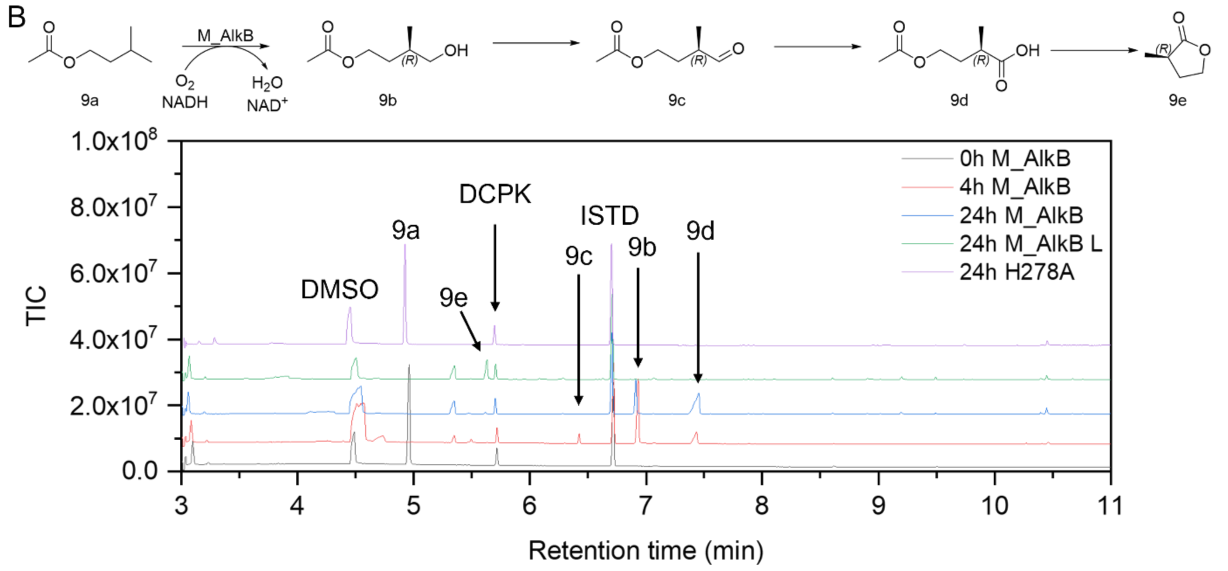

**Figure S14.** – continued.

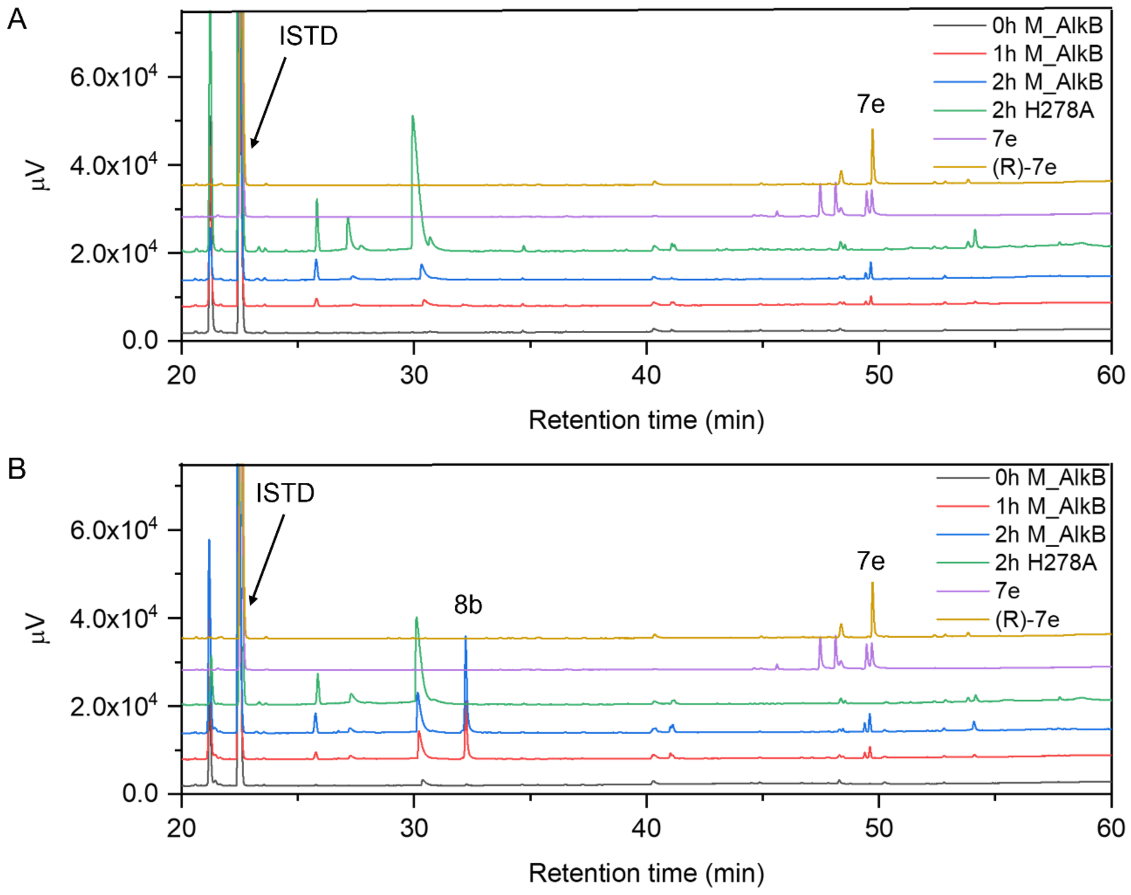

**Figure S15.** Exemplary chiral GC-FID chromatograms of reactions with the branched esters A) **7a** and B) **8a**. The transformations were performed using 3.1 $g_{CDW}$/L for 24 h at 25 °C, 180 rpm, and compared with a strain expressing an inactive variant M_AlkB (H278A) as a negative control and authentic standards of racemic **7e** and (*R*)-**7e**.

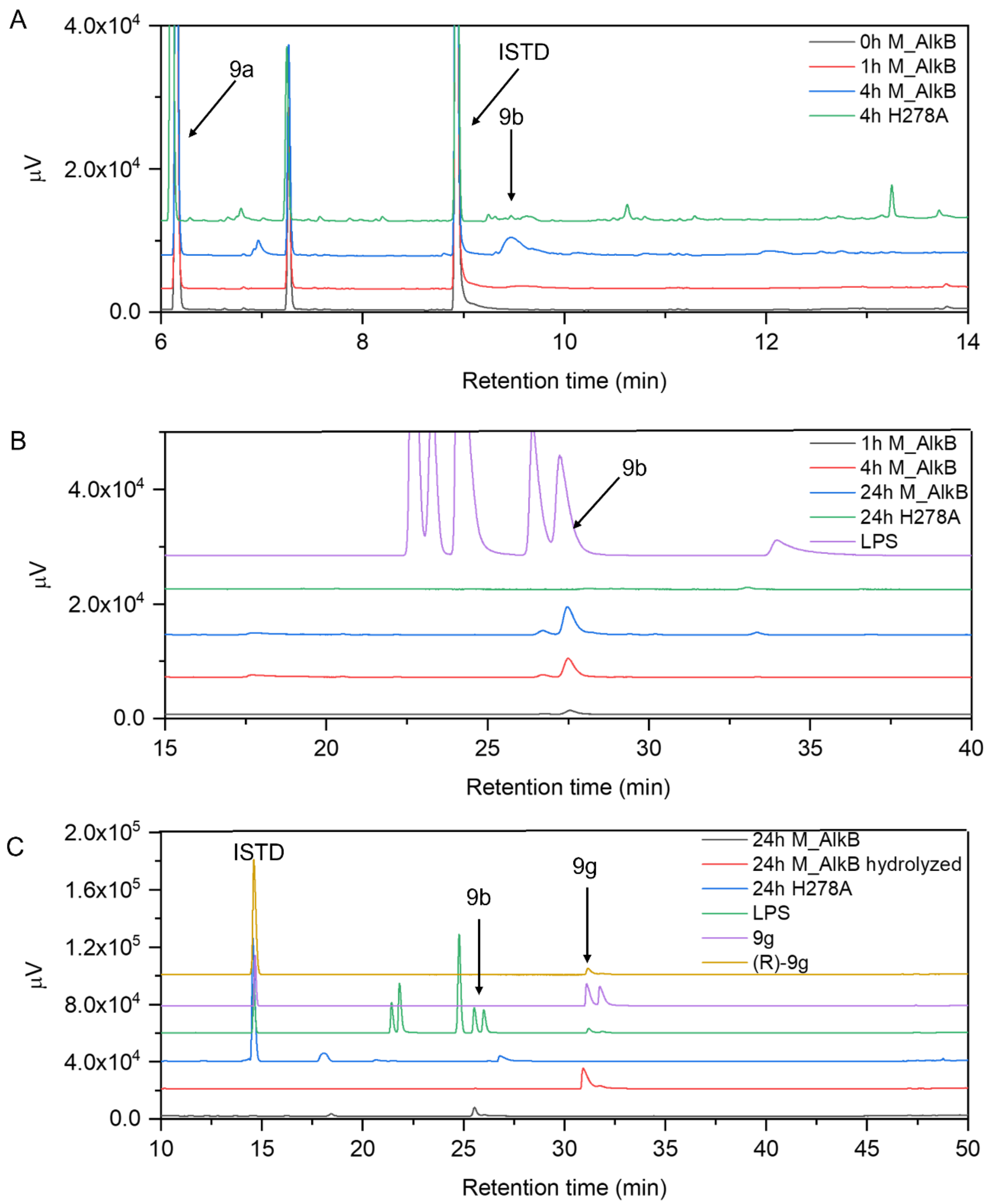

**Figure S16.** A) Exemplary achiral GC-FID chromatograms of reactions with the branched ester **9a** used for rate quantification B) Exemplary chiral GC-FID (β-TBDM column) analysis of reaction samples of M_AlkB catalyzed hydroxylation of **9a**, compared with the in-house synthesized sample **9b** (LPS); C) Exemplary chiral GC-FID (β-TBDAc column) analysis of reaction samples of M_AlkB catalyzed hydroxylation of **9a**, hydrolyzed samples compared with the in-house synthesized sample **9b** (LPS), and to authentic standards of racemic **9g** and (*R*)-**9g**; D) Exemplary chiral GC-FID (β-TBDAc column) analysis of reaction samples of M_AlkB catalyzed hydroxylation of 9a using DMSO as a cosolvent, lactonized sample (L) and authentic standard of **9e**. All reactions were compared with a strain expressing an inactive variant M_AlkB (H278A) as a negative control. The biotransformations were performed using 3.1 $g_{CDW}$/L for 24 h at 25 °C, 180 rpm.

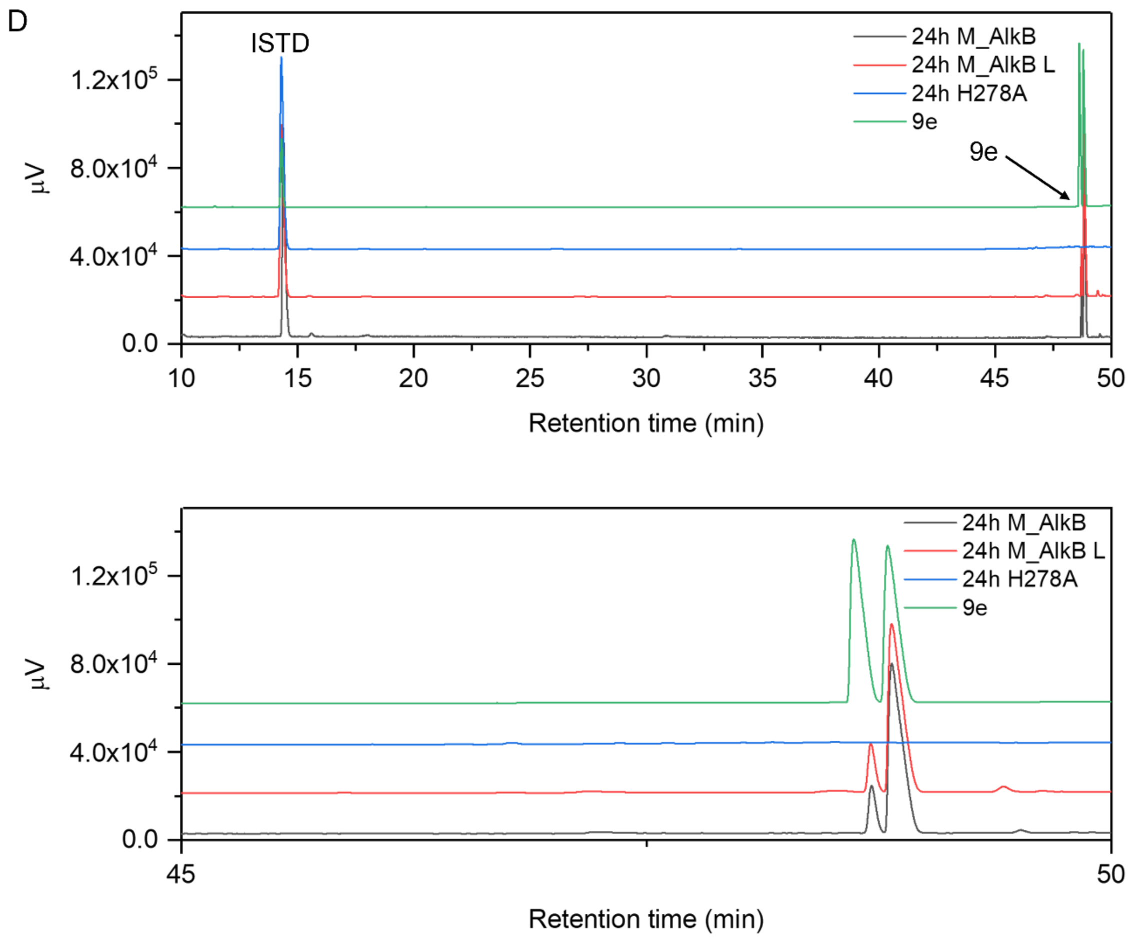

**Figure S16**. – continued.

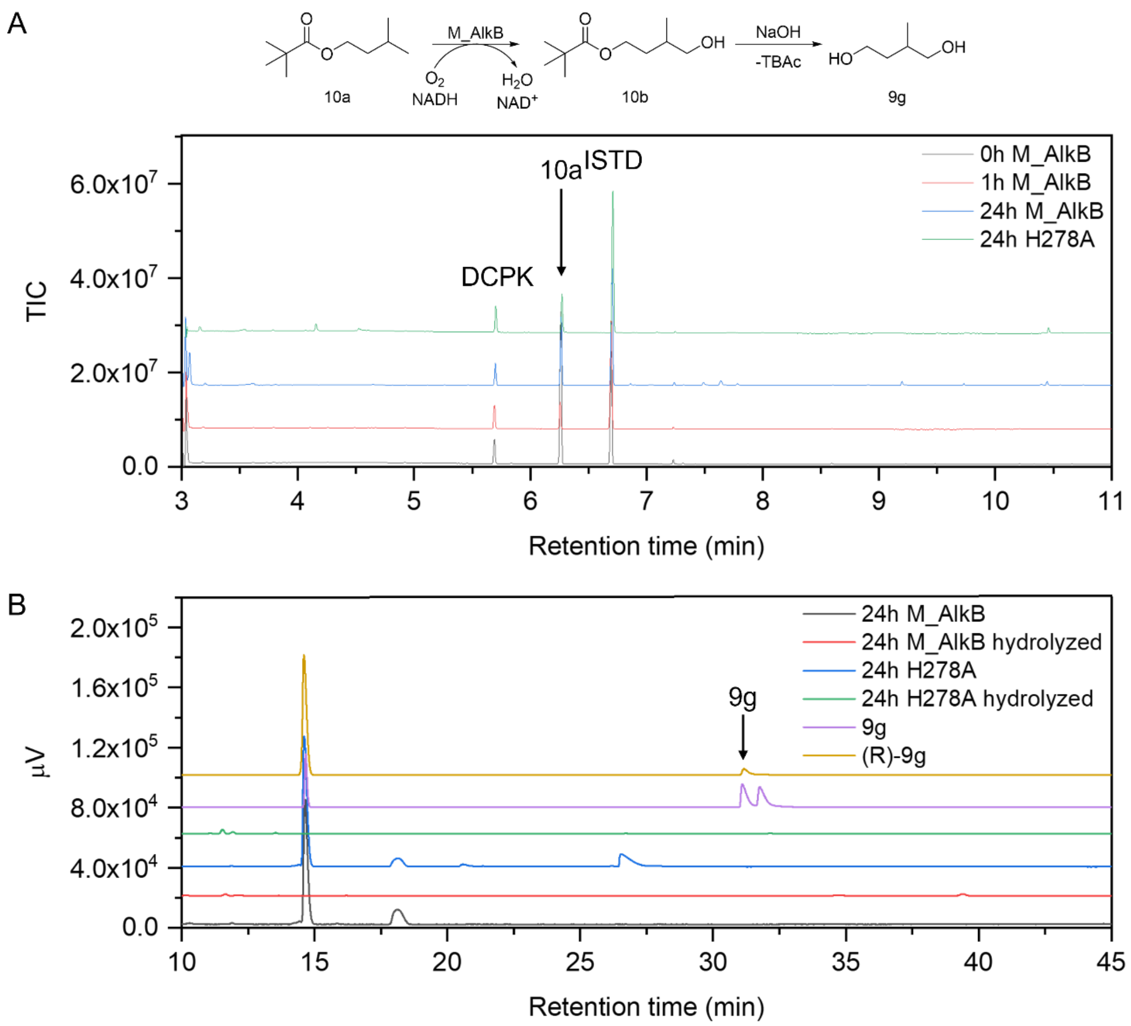

**Figure S17.** A) GC-MS analysis of initial activity screening of M_AlkB with **10a** and B) chiral GC-FID analysis of hydrolyzed 24 h reaction sample. The reaction samples were compared to an authentic standard of racemic **9g** and (*R*)-**9g**. The transformations were performed using 3.1 $g_{CDW}$/L for 24 h at 25 °C, 180 rpm.

### 8 Summary of activity screening of AlkB variants

**Table S18.** Summary of activity screening of AlkB wt and variants with linear esters **2a**, **4a**, **5a**, and **6a**. Data shown as arithmetic mean ± SD (n = 3).

| Enzyme | Variant | | Activity (U/$g_{CDW}$) | | | | Product (mM)[a] | | | |
|---|---|---|---|---|---|---|---|---|---|---|
| | | $g_{CDW}$/L | **2a** | **4a** | **5a** | **6a** | **2b** | **4b** | **5b** | **6b** |
| **PpAlkB** | wt | 1 | 7.70 ± 1.76 | 0.34 ± 0.24 | 0.80 ± 0.46 | n. d. | 0.61 ± 0.001 | 0.01 ± 0.001 | 0.12 ± 0.07 | n. d. |
| **M_AlkB** | wt | 1 | 5.84 ± 1.16 | 2.65 ± 0.64 | 4.77 ± 0.69 | n. d. | 0.61 ± 0.12 | 0.32 ± 0.04 | 0.56 ± 0.07 | n. d. |
| | | 3.1 | 4.86 ± 0.13 | 3.24 ± 0.28 | 4.72 ± 0.29 | n. d. | 0.47 ± 0.06 | 0.51 ± 0.15 | 0.74 ± 0.05 | n. d. |
| | I238V | 1 | 9.73 ± 1.46 | 3.51 ± 0.63 | 5.57 ± 0.77 | n. d. | 0.92 ± 0.13 | 0.51 ± 0.09 | 0.67 ± 0.14 | n. d. |
| | | 3.1 | 6.67 ± 0.97 | 6.42 ± 0.99 | 7.39 ± 0.99 | n. d. | 0.43 ± 0.02 | 0.52 ± 0.14 | 0.73 ± 0.15 | n. d. |
| | F169L | 1 | 10.53 ± 0.19 | 4.70 ± 0.22 | 6.40 ± 0.43 | n. d. | 0.83 ± 0.15 | 0.59 ± 0.01 | 0.63 ± 0.03 | n. d. |
| | | 3.1 | 7.05 ± 0.54 | 4.20 ± 0.17 | 4.42 ± 0.59 | n. d. | 0.32 ± 0.05 | 0.27 ± 0.20 | 0.60 ± 0.07 | n. d. |
| | W60S | 1 | 3.40 ± 0.88 | 3.17 ± 0.23 | 2.10 ± 0.29 | n. d. | 0.36 ± 0.07 | 0.38 ± 0.07 | 0.25 ± 0.06 | n. d. |
| | | 3.1 | 3.20 ± 0.31 | 3.47 ± 0.47 | 3.38 ± 0.07 | 0.63 ± 0.17 | 0.32 ± 0.05 | 0.89 ± 0.02 | 0.82 ± 0.13 | 0.26 ± 0.08 |
| | W60S + I238V | 1 | n. d. | n. d. | n. d. | 0.26 ± 0.05 | n. d. | n. d. | n. d. | 0.03 ± 0.01 |
| | | 3.1 | n. d. | n. d. | n. d. | 0.79 ± 0.07 | n. d. | n. d. | n. d. | 0.20 ± 0.01 |

[a]obtained after 2 h; n. d.: not determined.

### 9 Reported C-H oxyfunctionalization by AlkB of sterically demanding substrates

**Table S19.** Overview of previously reported C-H oxyfunctionalization of sterically demanding substrates by AlkB homologs and variants.

| Enzyme | Variant | Substrate | Product(s) | Activity ($U/g_{CDW}$) | Reaction system | Ref. |
|---|---|---|---|---|---|---|
| **PpAlkB** | **wt** | **2- methyl pentane** | 2-methyl-1-pentanol[b] | 0.8 | *Pseudomonas putida* S81 pGEc41Δ*alkJ* (Bosetti, van Beilen, Preusting, Lageveen, & Witholt, 1992)<br>batch reaction; 25 mL Erlenmeyer flask;<br>4 mg/mL resting cells in 50 mM KPi, pH 7.0, 10 mM MgSO4, 0.05% Triton X-100<br>1 % (v/v) substrate; 3 mL reaction volume<br>30 °C, 200 rpm, 30 min<br>Activities relative to *n*-nonane[a] | (van Beilen, Kingma, & Witholt, 1994) |
| | | | 4-methyl-1-pentanol[b] | 1.3 | | |
| | | **3-methyl pentane** | 3-methyl-1-pentanol[b] | 1.9 | | |
| | | **2,4-dimethylpentane** | 2,4-dimethyl-l-pentanol[b] | 1.4 | | |
| | | **2-methylhexane** | 2-methyl-1-hexanol[b] | 6.0 | | |
| | | | 5-methyl-1-hexanol | 17.0 | | |
| | | **3-methylhexane** | 3-methyl-1-hexanol[b] | 14.0 | | |
| | | | 4-methyl-1-hexanol[b] | 3.3 | | |
| | | **2,5-dimethylhexane** | 2,5-dimethyl-l-hexanol[b] | 6.7 | | |
| | | **2-methylheptane** | 2-methyl-l-heptanol[b] | 5.1 | | |
| | | | 6-methyl-l-heptanol[b] | 30.0 | | |
| | | **3-methylheptane** | 3-methyl-l-heptanol[b] | 5.7 | | |
| | | | 5-methyl-l-heptanol | 4.7 | | |
| | | | 4-methyl-l-heptanol | 9.3 | | |
| | | **2-methyloctane** | 2-methyl-l-octanol[b] | 3.3 | | |
| | | | 7-methyl-l-octanol[b] | 38.0 | | |
| | | **3-methyloctane** | 3-methyl-l-octanol[b] | 2.2 | | |
| | | | 6-methyl-l-octanol[b] | 15.0 | | |
| | | **4-methyloctane** | 4-methyl-l-octanol[b] | 10.0[c] | | |
| | | | 5-methyl-l-octanol[b] | | | |
| | | **2,6-dimethyloctane** | 3,7-dimethyl-l-octanol[b] | 4.0[c] | | |
| | | | 2,6-dimethyl-l-octanol[b] | | | |
| | | **2-methylnonane** | 2-methyl-l-nonanol[b] | 1.7 | | |
| | | | 8-methyl-l-nonanol[b] | 6.8 | | |
| | | **methylcyclohexane** | *trans*-4-methylcyclohexanol | 48.0 | | |
| | | **ethylcyclohexane** | *trans*-4-ethylcyclohexanol | 31.0 | | |
| | | **toluene** | benzyl alcohol | 15.0 | | |
| | | **ethylbenzene** | 2-phenyl-l-ethanol | 80.0 | | |
| | | ***n*-propylbenzene** | 3-phenyl-l-propanol | 5.0 | | |
| | | ***n*-butylbenzene** | 4-phenyl-l-butanol | 26.0 | | |
| | | **isopropyl benzene** | 2-phenyl-l-propanol | 28.0 | | |

**Table S19.** – continued.

| Enzyme | Variant | Substrate | Product(s) | Activity ($U/g_{CDW}$) | Reaction system | Ref. |
|---|---|---|---|---|---|---|
| PpAlkB | wt | **2-diethylbenzene** | 2-(2-ethylphenyl)-ethanol | 6.0 | *Pseudomonas putida* S81 pGEc41Δ*alkJ* (Bosetti et al., 1992)<br>batch reaction; 25 mL Erlenmeyer flask;<br>4 mg/mL resting cells in 50 mM KPi, pH 7.0, 10 mM MgSO4, 0.05% Triton X-100<br>1 % (v/v) substrate; 3 mL reaction volume<br>30 °C, 200 rpm, 30 min<br>Activities relative to *n*-nonane[a] | (van Beilen et al., 1994) |
| | | **3-diethylbenzene** | 2-(3-ethylphenyl)-ethanol | 107.0 | | |
| | | **4-diethylbenzene** | 2-(2-ethylphenyl)-ethanol | 85.0 | | |
| | | **2-ethyltoluene** | 2-(2-tolyl)-ethanol | 6.0 | | |
| | | **3-ethyltoluene** | 2-(3-tolyl)-ethanol | 134.0 | | |
| | | **4-ethyltoluene** | 2-(4-tolyl)-ethanol | 95.0 | | |
| | | ***o*-xylene** | 2-methylbenzylalcohol | 0.4 | | |
| | | ***m*-xylene** | 3-methylbenzylalcohol | 4.3 | | |
| | | ***p*-xylene** | 4-methylbenzylalcohol | 2.8 | | |
| | | **allylbenzene** | 3-phenyl-l,2-epoxypropane | 22.0 | | |
| | | **allyl phenyl ether** | pheny1g1ycidylether | 11.0 | | |
| PpAlkB | wt<br>(or variant) | **isobutene** | 2-methyl-2-propen-1-ol<br>methacrylic acid | n. r. | *E. coli* W3110 pBT10 (Schrewe, Magnusson, Willrodt, Bühler, & Schmid, 2011)<br>continuous reaction, 300 mL fermenter,<br>600 mg/mL resting cells in 70 mM ammonium phosphate buffer pH 7, 0.5 g/L NaCl, 0.49 g/L $MgSO_4$ x $7H_2O$, 1 mL/L trace element solution US3, 50 g/l kanamycin<br>Glc either as batch (10 g/L) or fed (1.8 g/h L)<br>Substrate feed: 15 Nl/h gas mixture (25% isobutene and 75% synthetic air); 50 mL start volume<br>40 °C, 900 rpm, 420 min | (Patent No. US20150010968A1, 2015) |
| PpAlkB | wt | **Isoprenyl acetate** | 4-acetoxy-2-methylene-butan-1-ol | 0.5 | *E. coli* BL21(DE3) pPpAlkB(mut)-FGTL or pM_AlkB(mut)-FGTL<br>Batch reaction, 1.5 mL glass vial<br>3.1 $g_{CDW}$/L resting cells, 50 mM Kpi, pH 7.4; 2 mM $MgSO_4$, 1% Glc;<br>5 mM substrate; 0.3 mL total reaction volume<br>25 °C; 180 rpm, 60 min | (Nigl et al., 2025) |
| | F164L | | | 1.1 | | |
| | I233V | | | 0.8 | | |
| | F164L_I233V | | | 0.9 | | |
| M_AlkB | wt | | | 1.0 | | |
| | F169L | | | 0.9 | | |
| | I238V | | | 1.8 | | |
| | F169L_I238V | | | 0.6 | | |

[a]normalization to *n*-nonane activity (10-15 $U/g_{CDW}$) of individual experiment; [b]products tentatively identified; [c]only one product peak but two products expected; n. r. not reported.

### 10 References

Abramson, J., Adler, J., Dunger, J., Evans, R., Green, T., Pritzel, A., … Jumper, J. M. (2024). Accurate structure prediction of biomolecular interactions with AlphaFold 3. *Nature*, *630*(8016), 493–500. doi: 10.1038/s41586-024-07487-w

Bosetti, A., van Beilen, J. B., Preusting, H., Lageveen, R. G., & Witholt, B. (1992). Production of primary aliphatic alcohols with a recombinant *Pseudomonas* strain, encoding the alkane hydroxylase enzyme system. *Enzyme Microb Technol*, *14*(9), 702–708. doi: 10.1016/0141-0229(92)90109-2

Chai, J., Guo, G., McSweeney, S. M., Shanklin, J., & Liu, Q. (2023). Structural basis for enzymatic terminal C–H bond functionalization of alkanes. *Nat Struct Mol Biol*, *30*(4), 521–526. doi: 10.1038/s41594-023-00958-0

Daina, A., Michielin, O., & Zoete, V. (2017). SwissADME: a free web tool to evaluate pharmacokinetics, drug-likeness and medicinal chemistry friendliness of small molecules. *Sci Rep 2017 7:1*, *7*(1), 1–13. doi: 10.1038/srep42717

Engel, P., Haas, T., Pfeffer, J. C., Thum, O., & Gehring, C. (2015). *Patent No. US20150010968A1*. United States. Retrieved from https://patents.google.com/patent/US20150010968/ar

Expasy - SIM Alignment Tool. (n.d.). Retrieved from https://web.expasy.org/sim/

Ferraboschi, P., Grisenti, P., Manzocchi, A., & Santaniello, E. (1994). Regio- and enantioselectivity of *Pseudomonas cepacia* lipase in the transesterification of 2-substituted-1,4-butanediols. *Tetrahedron: Asymmetry*, *5*(4), 691–698. doi: 10.1016/0957-4166(94)80031-6

Gasteiger, E. (2003). ExPASy: the proteomics server for in-depth protein knowledge and analysis. *Nucleic Acids Res*, *31*(13), 3784–3788. doi: 10.1093/nar/gkg563

Grisenti, P., Ferraboschi, P., Casati, S., & Santaniello, E. (1993). Studies on the enantioselectivity of the transesterification of 2-methyl-1,4-butanediol and its derivatives catalyzed by *Pseudomonas fluorescens* lipase in organic solvents. *Tetrahedron: Asymmetry*, *4*(5), 997–1006. doi: 10.1016/S0957-4166(00)80144-5

Groves, J. T., Feng, L., & Austin, R. N. (2023). Structure and function of alkane monooxygenase (AlkB). *Acc Chem Res*, *56*(24), 3665–3675. doi: 10.1021/acs.accounts.3c00590

Guo, X., Zhang, J., Han, L., Lee, J., Williams, S. C., Forsberg, A., … Feng, L. (2023). Structure and mechanism of the alkane-oxidizing enzyme AlkB. *Nat Commun*, *14*(1), 2180. doi: 10.1038/s41467-023-37869-z

Julsing, M. K., Schrewe, M., Cornelissen, S., Hermann, I., Schmid, A., & Bühler, B. (2012). Outer membrane protein AlkL boosts biocatalytic oxyfunctionalization of hydrophobic substrates in *Escherichia coli*. *Appl Environ Microbiol*, *78*(16), 5724–5733. doi: 10.1128/aem.00949-12

Kuipers, R. K., Joosten, H.-J., van Berkel, W. J. H., Leferink, N. G. H., Rooijen, E., Ittmann, E., … Schaap, P. J. (2010). 3DM: Systematic analysis of heterogeneous superfamily data to discover protein functionalities. *Proteins: Struct Funct Bioinf 78*(9). doi: 10.1002/prot.22725

Marques, S. M., Borko, S., Vavra, O., Dvorsky, J., Kohout, P., Kabourek, P., … Bednar, D. (2025). Caver Web 2.0: analysis of tunnels and ligand transport in dynamic ensembles of proteins. *Nucleic Acids Res*, *53*(W1), W132–W142. doi: 10.1093/nar/gkaf399

Mikulska-Ruminska, K., Licht, M., Ertem, M. Z., Shanklin, J., Liu, Q., & Bahar, I. (2025, June 27). Substrate binding and channeling allosterically modulate the interactions within the AlkB-AlkG electron transfer complex. *BioRxiv*, p. 2025.06.23.661152. Cold Spring Harbor Laboratory. doi: 10.1101/2025.06.23.661152

Nigl, A., Delsoglio, V., Sovic, L., Grgić, M., Malihan-Yap, L., Myrtollari, K., … Kourist, R. (2025). Engineering of transmembrane alkane monooxygenases to improve a key reaction step in the synthesis of polymer precursor tulipalin A. *Angew Chem Int Ed*, *64*(25). doi: 10.1002/anie.202503464

Schrewe, M., Magnusson, A. O., Willrodt, C., Bühler, B., & Schmid, A. (2011). Kinetic analysis of terminal and unactivated C–H bond oxyfunctionalization in fatty acid methyl esters by monooxygenase-based whole-cell biocatalysis. *Adv Synth Catal*, *353*(18), 3485–3495. doi: 10.1002/adsc.201100440

Schrödinger LLC. (2025). *Schrödinger Release 2025-2: Maestro*. New York.

Smits, T. H. M., Seeger, M. A., Witholt, B., & van Beilen, J. B. (2001). New alkane-responsive expression vectors for *Escherichia coli* and *Pseudomonas*. *Plasmid*, *46*(1), 16–24. doi: 10.1006/plas.2001.1522

Stourac, J., Vavra, O., Kokkonen, P., Filipovic, J., Pinto, G., Brezovsky, J., … Bednar, D. (2019). Caver Web 1.0: identification of tunnels and channels in proteins and analysis of ligand transport. *Nucleic Acids Res*, *47*(W1), W414–W422. doi: 10.1093/nar/gkz378

van Beilen, J. B., Kingma, J., & Witholt, B. (1994). Substrate specificity of the alkane hydroxylase system of *Pseudomonas oleovorans* GPo1. *Enzyme Microb Technol*, *16*(10), 904–911. doi: 10.1016/0141-0229(94)90066-3

van Beilen, J. B., Smits, T. H. M., Roos, F. F., Brunner, T., Balada, S. B., Röthlisberger, M., & Witholt, B. (2005). Identification of an amino acid position that determines the substrate range of integral membrane alkane hydroxylases. *J Bacteriol*, *187*(1), 85–91. doi: 10.1128/jb.187.1.85-91.2005

van Nuland, Y. M., Eggink, G., & Weusthuis, R. A. (2016). Application of AlkBGT and AlkL from *Pseudomonas putida* GPo1 for selective alkyl ester ω-oxyfunctionalization in *Escherichia coli*. *App Environ Microbiol*, *82*(13), 3801–3807. doi: 10.1128/aem.00822-16

Waterhouse, A. M., Procter, J. B., Martin, D. M. A., Clamp, M., & Barton, G. J. (2009). Jalview Version 2—a multiple sequence alignment editor and analysis workbench. *Bioinformatics*, *25*(9), 1189–1191. doi: 10.1093/bioinformatics/btp033
